# “Cold” Orthogonal Translation: A Psychrophilic Pyrrolysyl-tRNA Synthetase Boosts Genetic Code Expansion in *E. coli*

**DOI:** 10.1101/2023.05.23.541947

**Authors:** Nikolaj G. Koch, Peter Goettig, Michael A. Nash, Juri Rappsilber, Nediljko Budisa

## Abstract

Using orthogonal translation systems (OTSs) is one of the most efficient strategies for producing unnatural proteins through the incorporation of non-canonical amino acids (ncAAs) into the genetic code. Traditionally, efforts to expand substrate specificity start with a (hyper-)stable enzyme capable of withstanding the structural changes induced by necessary mutations. In contrast, we propose a radically different approach for the PylRS system: by starting with enzymes that evolved to cope with instability in order to adapt to cold conditions, potentially offering greater resilience to mutational changes. By finding and further engineering a psychrophilic (“cold”) OTS from *Methanococcoides burtonii*, we developed an alternative to the widely used mesophilic and thermophilic systems. This novel OTS demonstrated exceptional *in vivo* efficiency for a broad range of substrates, even at very low ncAA concentrations and low cultivation temperatures. The general versatility of the PylRS system across a wide range of applicable host organisms suggests that the Cold-OTS has the potential to also improve protein yields in these hosts and help to drive the transformation of the expanded genetic code from an academic pursuit into a high-value, chemistry-driven biotechnology.

## Introduction

New-to-nature non-canonical amino acids (ncAAs) expand the functional repertoire of proteins.^1–4^ Their specific incorporation relies on orthogonal translation systems (OTSs) composed of engineered orthogonal aminoacyl-tRNA synthetase/tRNA pairs. To date, over 500 ncAAs have been incorporated into proteins using various genetic code expansion (GCE) strategies, with more than 300 achieved using the PylRS system alone.^1–5^ The most common method for site-specific incorporation is amber stop codon suppression (SCS). In SCS, the ncAA is incorporated in response to an in-frame stop codon placed at a predefined position within the target protein coding sequence, allowing ribosomal expression either *in vivo* or *in vitro*. Among GCE systems those derived from pyrrolysyl-tRNA synthetases (PylRS) are the most widely used due to their natural orthogonality in both prokaryotes and eukaryotes.^2,6,7^

The ideal OTS combines high catalytic efficiency with sufficient versatility or promiscuity, enabling straightforward engineering through minimal mutations. While known PylRS variants fulfill the requirement for versatility, they lack catalytic efficiency necessary for high *in vivo* activity.^8^ This limitation likely contributes to their modest performance in cellular contexts. Significant efforts have been made to progressively increase the in-cell efficiency of PylRS-based OTS for both wild-type and engineered systems. These efforts include optimizing OTS plasmid copy number and promotor strength of aaRSs and/or tRNA genes^9^, engineering tRNA^Pyl 10^, rational and semi-rational engineering of aaRS variants^11^, optimizing sequence context around the target codon^12^, host cell engineering^13^, engineering parts of the translational machinery (e.g., elongation factor TU)^14^ and increasing the solubility of the aaRS^15^. Despite these strategies, the yield of target protein production consistently falls short compared to that of the wild-type reference protein.

To shift the natural substrate recognition of PylRS or any other OTS to a desired ncAA, the enzyme must be engineered. A general strategy in enzyme engineering, including the GCE field, is to use (hyper-)thermophilic enzymes as starting scaffolds. These highly stable enzymes provide the necessary stability to tolerate the mutations required to remodel the active site. For example, the commonly used hyperthermophilic tyrosyl-tRNA synthetase of *Methanocaldococcus jannaschii* (*Mj*TyrRS) has highly specific substrate interactions, requiring a significant number of mutations – up to 10 – to acquire new substrate specificities.^16^ However, the *Mj*TyrRS’s high level of substrate specificity complicates engineering and reduces the likelihood and ease of successfully identifying desired variants.^16,17^ Another drawback of thermophilic enzymes is their typically lower catalytic activity under standard cultivation conditions compared to their mesophilic homologs, necessitating further engineering to enhance efficiency.^17–19^

Fortunately, the unique structure of Pyl has resulted in minimal evolutionary pressure for PylRS to develop a highly specific recognition mechanism. Instead, it relies on relatively nonspecific hydrophobic interactions.^20^ This recognition mode often leads to a high probability of encoding new substrates with only a few mutations, typically requiring one to four mutations (and never more than three in this study).^20,21^ This unique property among aminoacyl-tRNA synthetases (aaRSs) eliminates the need for an extremely stable starting scaffold during engineering campaigns.

Given these circumstances, we hypothesized that the PylRS OTS could enable a shift away from the prevalent engineering approaches to genetic code expansion. Specifically, this shift involves moving away from using mesophilic and thermophilic aaRSs. Enzymes derived from cold-adapted organisms are known to exhibit higher activity at ambient temperatures, provided they can tolerate these conditions. However, psychrophilic enzymes are often characterized by high flexibility, which is accompanied by structural instability at elevated temperatures.^18,22,23^ Despite this, we identified a psychrophilic homolog of the PylRS system with superior activity under standard laboratory cultivation conditions (18-37 °C). Remarkably, its performance is even greater at lower temperatures (18 °C) and with suboptimal substrates. We demonstrate that the *Methanococcoides burtonii* (*M. burtonii*) PylRS (Mbur) outperforms other variants in terms of both *in vivo* incorporation efficiency and substrate recognition scope.

## Results

### Initial screening of psychro-, meso- and thermophilic PylRS variants

To overcome the efficiency limitations of the current PylRS system, we turned our attention towards utilizing psychrophilic enzyme scaffolds. Simplified, the PylRS exists in two classes: the +N class, which includes an N-terminus, and the ΔN class, which lacks the N-terminus. To determine which +N PylRS to prioritize, we evaluated the optimal growth temperature (OGT) of a range of organisms (see **Figure 1A** and discussion in Supplement Section 1 for details). For the psychrophilic PylRS set, we included four variants from organisms with an OGT of 26°C or lower (additional information is provided in **Table S3** of the supplements). These were compared with two mesophilic and two thermophilic +N PylRS variants.

**Figure 1.**
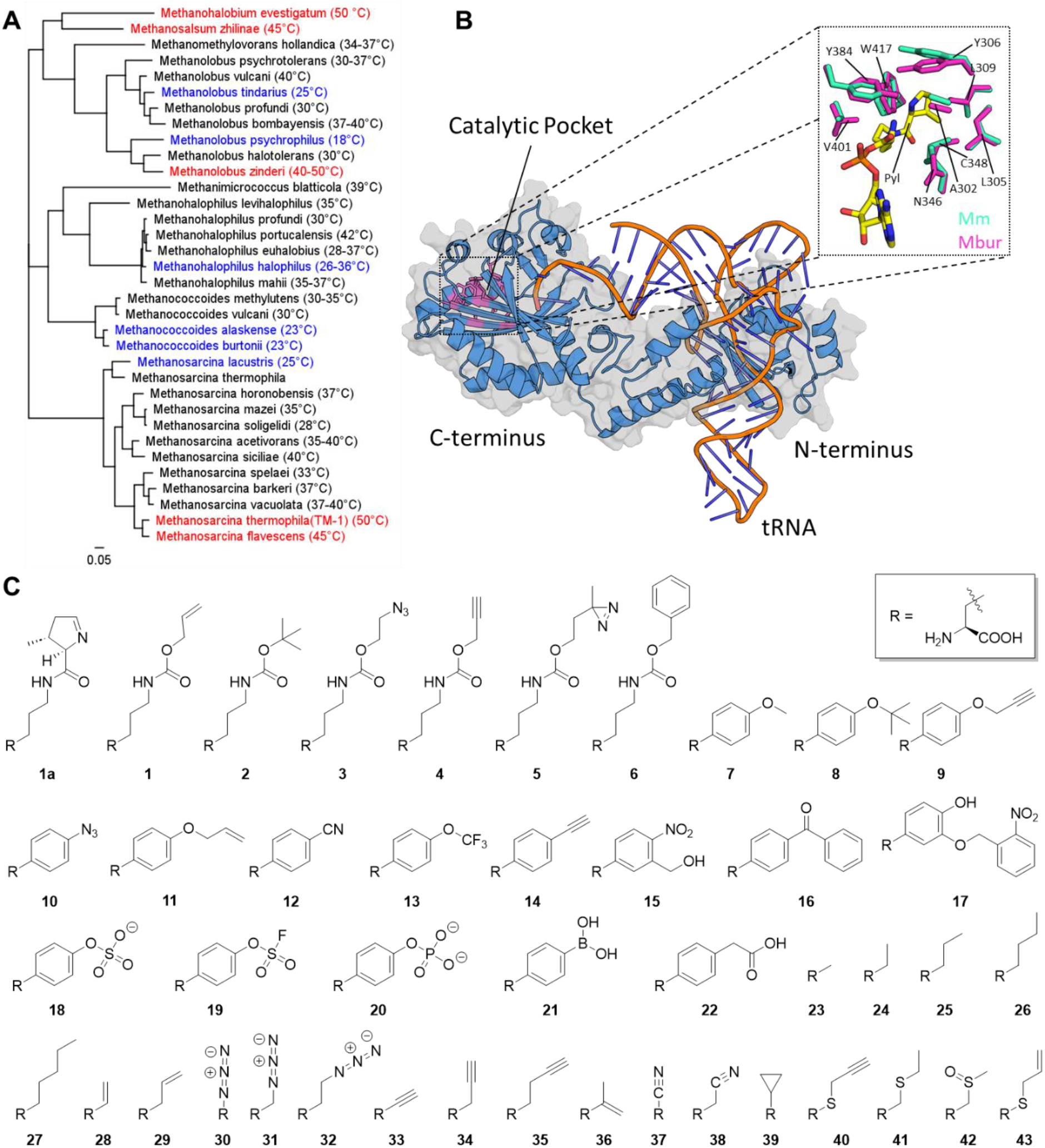
Integrating evolutionary phylogeny, structural biology, and non-canonical amino acid chemistry led to the discovery of a novel psychrophilic orthogonal translation system (“Cold-OTS”) based on PylRS. **A**) Phylogenetic tree based on all N-terminus containing PylRS enzymes of *Methanosarcinales* and *Methanomassiliicoccales* generated using the Approximate-Maximum-Likelihood method. The FastTree algorithm (version 2.1.11) was used to build the tree based sequences available in NCBI and *M. alaskense* from the JGI database.^32^ The numbers in parentheses represent the optimal growth temperature (OGT). Coloring indicates the origin of the psychrophilic (blue, equal to and below 26°C) and thermophilic (red, equal to and above 45°C) PylRSs. The scale bar indicates 5% sequence divergence. **B**) The 3D model structure (shown as cartoon) of *M. burtonii* PylRS (cyan, without linker) was created by aligning the N- and C-termini with a superposition of the corresponding termini of *M*.*mazei* (PDB ID 5UD5) and *Desulfitobacterium hafniense* (PDB ID 2ZNI) (see Supplementary Material for detailed discussion). The catalytic pocket with bound Pyl-AMP (yellow) is from *M. mazei* (PDB ID: 2Q7H) while the *M. burtonii* structure was calculated with ColabFold.^21,33^ The conserved active site residues are shown as sticks, with *M. burtonii* PylRS in red and *M. mazei* PylRS in blue (*M. mazei* notation). For clarity and because the relative position is not clear, the linker region has been omitted but can be found in the Supplementary Material. **C**) Non-canonical amino acid substrates used in this study. The systematic and trivial names of these ncAAs are listed in the Supplementary Material (**Table S1**). The compounds **1, 3, 4, 9**-**11, 14, 23**-**30, 35**, and **38** are suitable for click reactions either with 1,3 dipolar cycloaddition, inverse electron-demand Diels–Alder or the photo-catalyzed thiol-ene coupling reaction.^34^

Certain members of the ΔN PylRS class have been reported to exhibit high activity ^24,25^, at least comparable to *Methanosarcina mazei* (Mm)^26^. While no psychrophilic ΔN PylRS is currently known, our primary goal was to identify the most efficient variant overall. Therefore, we also included the 15 best performing ΔN PylRS and their corresponding tRNAs, as identified in a recent study^27^. These selections were based on performance and representation across the ΔN subgroups (A, B, C or S)^27^ to ensure a diverse set. Additionally, for *Methermicoccus shengliensis* (Sheng, OGT 65 °C) we considered its thermophilic origin.^28^

In total, we tested a set of 24 PylRS variants in combination with six different Pyl analogs (**1, 2, 3, 4, 5, 6**), each with known differences in substrate activity (**Figure 1C**). These substrates were categorized into three groups based on activity data for the +N class: good (**1, 2**)^29,30^, moderate (**3, 4, 5**^31^) and poor (**6**; which exhibits very low activity even at 3.5 mM concentrations^30^). We have previously shown that adding an SmbP-tag to wild-type Mb significantly improves its activity; therefore, this construct was used throughout our study.^15^ **Figure 1A** presents the phylogenetic tree for the +N class. Additional discussions on the amino acid composition of the +N class an the unique features of the *Methanosarcina* PylRSs variants can be found in the supplements (section 1.2-1.4).

All the selected +N PylRS were functional under standard cultivation conditions, albeit with varying levels of activity (**Figure 2**, upper half). As expected, the poor substrate **6** resulted to minimal activity, even at the highest ncAA concertation (9 mM, see **Figure S5**). Encouragingly, none of the psychrophilic PylRSs showed significantly higher background incorporation, indicating that they can discriminate against canonical amino acids (cAAs) with fidelity comparable to commonly used PylRS systems.

**Figure 2.**
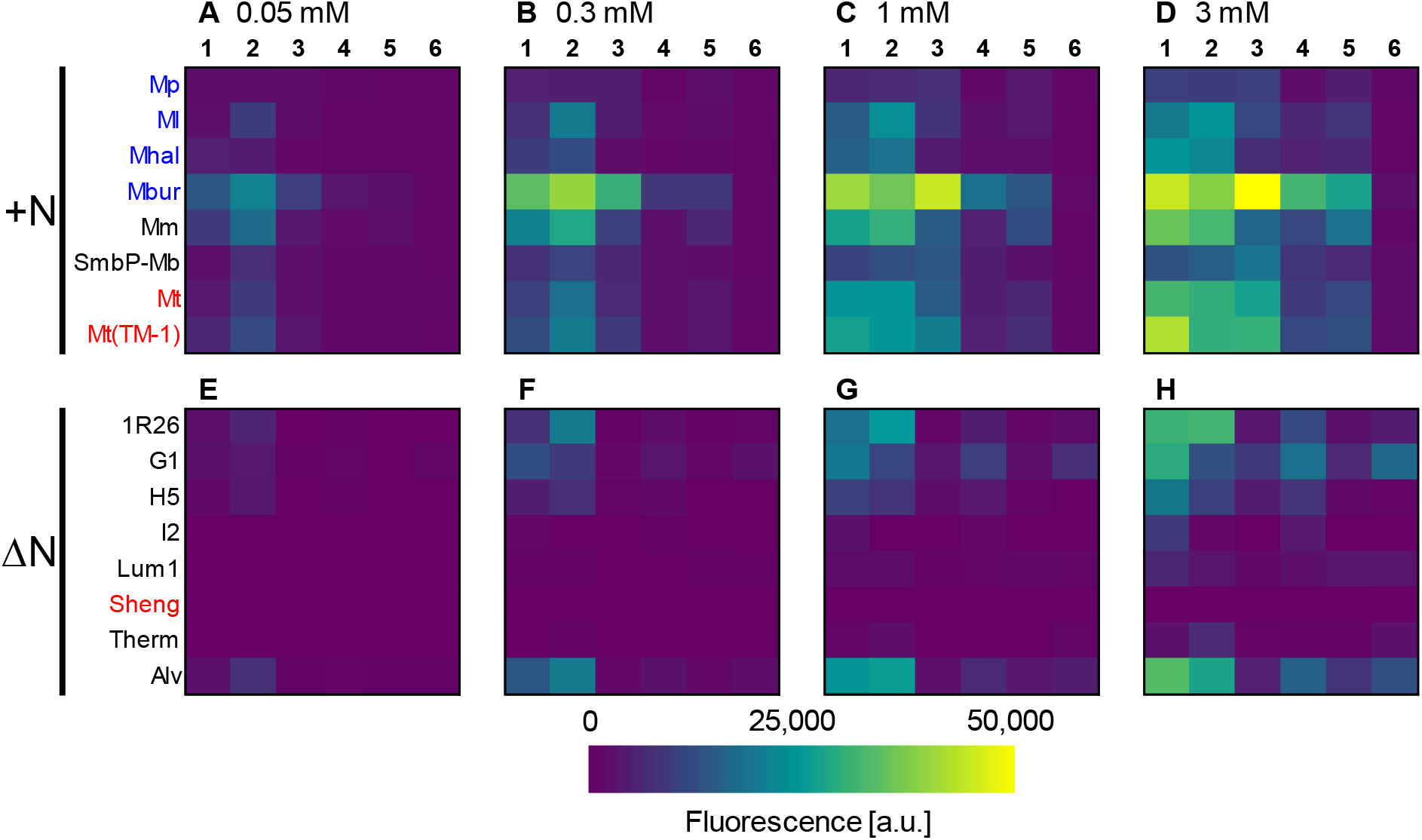
Heatmaps showing the fluorescence of sfGFP(1x amber) expression with *E. coli* BL21(DE3) for the eight selected PylRS variants, as a function of varying ncAA concentrations. The concentrations of ncAA supplied are: **A)&E), B)&F), C)&G), D)&H)**, are 0.05, 0,3, 1, 3 mM. Heatmaps for 0.1 and 9 mM are provided in the supplements (**Figure S5**&**6**). Bar charts with the number of replicates and error bars representing the standard deviation can be found in the supplements (**Figures S20-S37**). The Heatmaps for all 15 ΔN PylRS and the full ncAA concentration range can be found in the supplements (**Figure S6**). Substrate numbering corresponds to that in **Figure 1**. Background suppressions ranged between 900 and 1,300 [a.u.] depending on the construct.

Among the four tested psychrophilic PylRS variants, Mbur stood out as a clear outlier, consistently demonstrating superior OTS efficiency for all substrates across all tested concentrations. The maximal efficiency of Mbur for substrate **2** was achieved at just 0.3 mM. Overall, Mbur displayed exceptional performance at lower ncAA concentrations, indicating a more efficient OTS. For example, with substrate **2**, its performance at 0.05 mM was 30% higher than other variants, while at 0.3 and 1 mM the Mbur shows between 300% (for **3**) and 400% (for **4**) higher efficiency than the second-best variant. In contrast, the other psychrophilic PylRSs performed less effectively, with *Mp* barely functional, producing signals only marginally above background suppression levels. Since *Mp* is the most psychrophilic variant, 37°C cultivation temperature might be too high for this variant.

For the two mesophilic PylRS variants, the results were mixed. Mm exhibited higher efficiency than SmbP-Mb at low ncAA concentrations, consistent with previous findings.^11^ However, SmbP-Mb outperformed Mm at high concentrations (3 and 9 mM) of ncAA **3**. The thermophilic Mt(TM-1) performed similarly to Mm at ncAA concentrations of 1 mM and above. Unlike highly thermophilic enzymes, which often show little to no activity at 37 °C, Mt(TM-1) maintained activity under these conditions, likely due to its origin from a mildly thermophilic organism.

Among the 15 ΔN PylRS variants tested, the Nitra, RumEn, Sheng, Tron and Clos constructs were completely nonfunctional in our OTS setup (**Figure S6**). This was unexpected, as all these 15 ΔN constructs had previously been reported as functional. Additionally, our OTS setup, based on the pTECH vector, is the most efficient in our hands - outperforming even the widely used pUltra setup^9^ (**Figure S7**). Therefore, any OTS with *in vivo* activity should have been detectable in this setup.

From the 15 ΔN PylRS variants, we selected the seven best performing ΔN PylRS constructs for inclusion in **Figure 2** (lower half), along with the thermophilic Sheng variant. As shown in **Figure 2**, even the best-performing ΔN constructs (1R26, G1, H5 and Alv) exhibited, at most, only half the *in vivo* activity of the Mbur construct - and this was observed only at ncAA concentrations of 1 mM or higher (for substrates **1**,**2** and **4**). Other constructs, such as 030, Deb and Int, displayed minimal activity and required very high ncAA concentrations to produce signals distinguishable from background suppression (**Figure S6**).

### Efficiency characterization of selected PylRS variants by increasing the number of ncAAs incorporated

As discussed in a recent review^2^, using an *E. coli* strain with release factor 1 (RF1) suppression of a single stop codon (as in **Figure 2)** provides a basic indication of the potential *in vivo* efficiency of an OTS. However, the dynamic range of the reporters used in normal strains is often too narrow to fully capture efficiency differences, particularly when reporter constructs with multiple stop codons are included (see **Figure S14** and **S15**). This limited dynamic range makes it difficult to appropriately resolve efficiency differences.

To address this limitation, we used an RF1 knock-out strain in conjunction with reporter constructs containing 1, 3 and 5 stop codons.^35^ In the absence of the amber termination machinery, multiple incorporations of a desired ncAA can be achieved, enabling the detection of even smaller efficiency differences *in vivo*. In this setup, not only does the peak y-value provide valuable information, but also the relative distances between the y-values for the different reporters also offer insights into the efficiency of the system.

We selected the four best performing +N and ΔN variants and tested their ability to perform multiple ncAA incorporations using substrates **1**,**2**,**3** and **4**, in the B-95.ΔA strain (**Figure S8&9**). From this dataset, we identified the two best performing +N and ΔN variants, alongside the *Mbur*, for further analysis (**Figure 3**).

**Figure 3.**
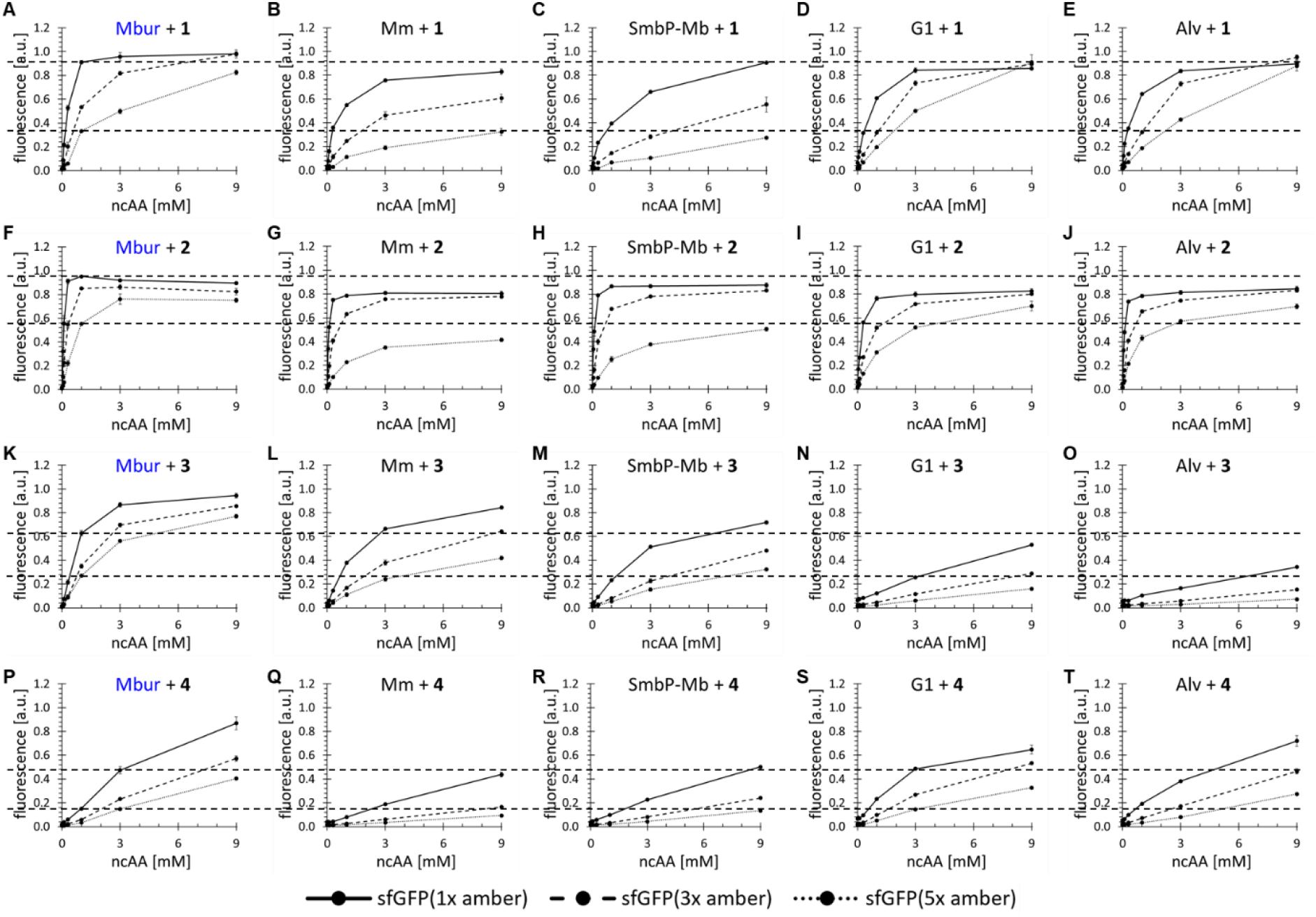
Concentration-dependent unnatural protein production using the two best performing +N and ΔN PylRS variants from **Figure 2**, alongside the Mbur variant. Protein production was conducted in RF1 deficient *Escherichia coli* B-95.ΔA. Endpoint measurements were performed at ncAA concentrations of 0.05, 0.1, 0.3, 1, 3 and 9 mM. Fluorescence values were normalized to their corresponding wild-type sfGFP reporter constructs (without in-frame stop codon). Error bars represent the standard deviation of three replicates (n=3). The upper dashed line indicates the 1 mM y-value for the Mbur/1x amber combination, while the lower one indicates the Mbur/5x amber, except for substrate **4**, there it is at 3 mM, due to the lower efficiency. Substrates and PylRS constructs are indicated in the headline.

All constructs performed best with substrate **2**, except for H5, which showed slightly better performance with substrate **1** when suppressing a single stop codon. For substrate **2**, Mbur clearly outperforms the other variants, particularly with the 5x amber construct. Mbur achieved near wild-type signal levels and exhibited only ∼20% signal loss at 3 mM, compared to ∼40-50% signal loss for the other variants (**Figure 3F-J**).

For substrate **1**, the two ΔN variants (G1 and Alv) perform comparably to Mbur at 3 mM but were clearly outperformed at lower concentrations, as shown on both single and multiple ncAA incorporation experiments (**Figure 3A-E**). For example, at 1 mM, the two ΔN variants reached only 60% of the wild-type signal, whereas Mbur exceeded >90%.

For substrate **3**,Mbur demonstrated significantly better performance in terms of both wild-type signal levels and efficiency at lower concentrations. For substrate **4**, G1 and Alv are clearly better than Mm and SmbP-Mb and perform on par with the Mbur until reaching 9 mM where the Mbur showed a higher maximum signal (**Figure 3P-T**). This aligns with observations from **Figure 1**, as substrate **4** is consistently the weakest performer among the +N variants.

### Flow cytometry analysis of the best performing PylRS variants for temperature dependent performance

In recombinant protein production with *E. coli*, many target proteins are either sequestered into inclusion bodies or fail to fold correctly. A common strategy to mitigate these issues is to slow down the protein production of *E. coli* by lowering the cultivation temperature.^36^ While this approach might alleviate problems with specific target proteins, it may also reduce protein yields in the context of GCE due to decreased activity of the OTS.

Since the Mbur construct originates from a psychrophilic organism, we aimed to determine whether its performance differs from other PylRS constructs at lower cultivations temperatures. To test this, we selected the four next best-performing +N and ΔN constructs, alongside Mbur (**Figure 4**). We used the sfGFP(5x amber) reporter construct, which provides a robust dynamic reporter signal range, and tested substrates **1-5** to obtain a comprehensive performance profile. For substrates **1-3**, we assessed activity at both 1 mM and 0.3 mM ncAA concentrations to capture concentration-dependent changes, as these are high-performing substrates.

**Figure 4.**
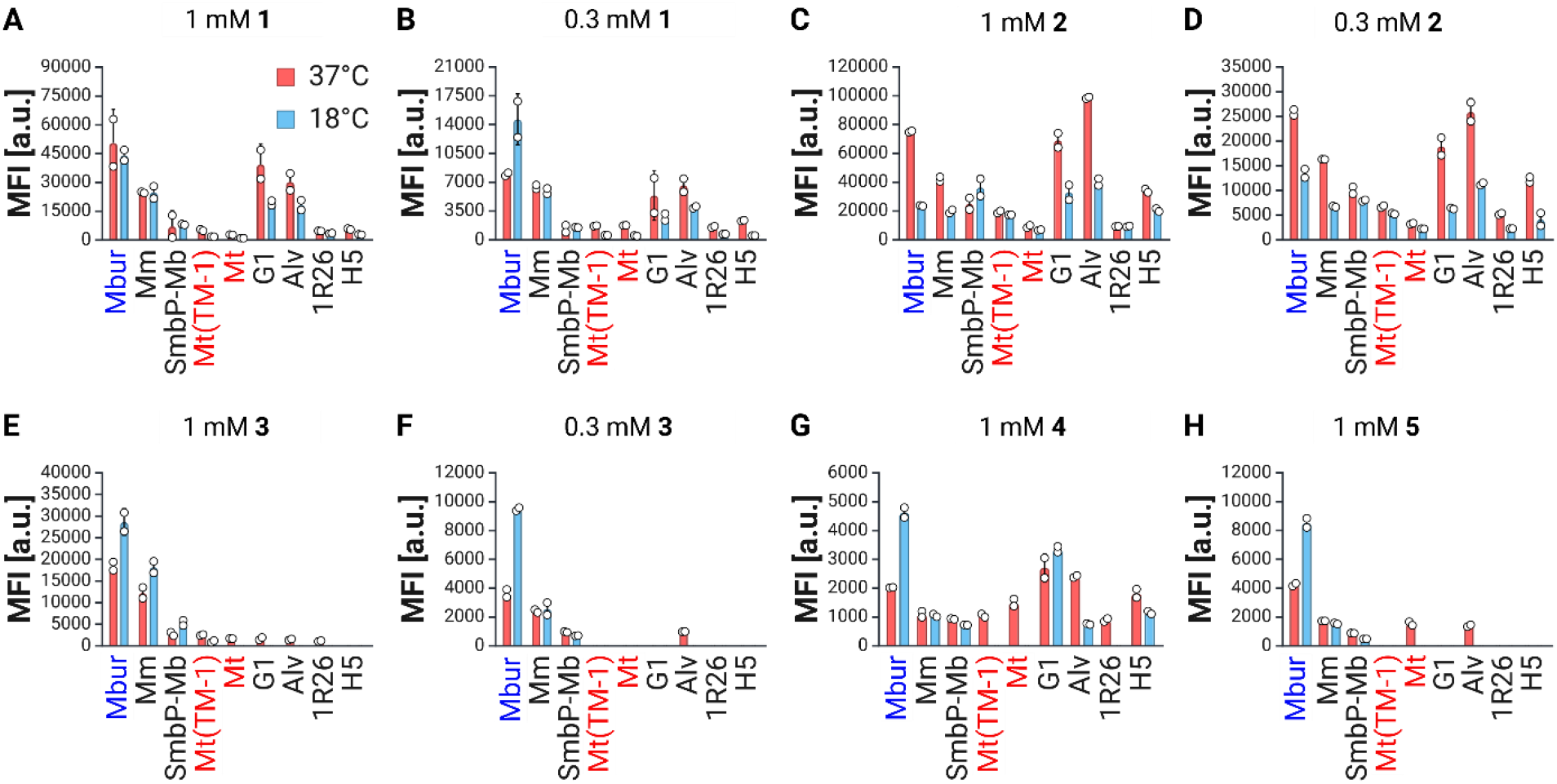
Temperature dependent production of sfGFP(5x amber) in the B-95.ΔA strain at 37°C(red columns) and 18°C(blue columns), supplied with different ncAA concentrations. Panels show results for: **A**) 1 mM **1, B**) 0.3 mM **1, C**) 1 mM **2, D**) 0.3 mM **2, E**) 1 mM **3, F**) 0.3 mM **3, G**) 1 mM **4** and **H**) 1 mM **5**. Constructs were analyzed using flow cytometry in biological duplicates (n=2), with the median fluorescence (MFI) of the target population displayed. Missing bars indicate the absence of a signal above background suppression. Example dotplots for the Mbur construct, from which the data was derived, are provided in the supplements (**Figure S93-S95**).

For all substrates, Mbur consistently performs the best, regardless of temperature, or ranks within the top three performers alongside G1 and Alv for ncAA **2**. For substrate **2**, which previously demonstrated superior performance (**Figure 3**), Mbur shows relatively better efficiency at 0.3 mM to 1 mM, with this advantage being even more pronounced at 18°C. Most notably, for 5 out of these 8 conditions, Mbur exhibited higher absolute performance at 18°C than at 37°C. We observed that for Mbur, the peak signal was reached after 72h at 0.3 mM concentrations for substrates **1**,**2** and **3** as well as at 1 mM for substrates **4** and **5**. However, for substrates **1** and **2** at 1 mM, the peak signal was achieved after 48h. This suggests that substrates with lower recognition benefit from longer cultivations times at lower temperatures. It was also generally observable that at 18°C, the non-expressing population was almost non-existent compared to cultures grown at 37°C, a trend observed across all constructs (**Figure S93-S95**).

It is well established that the overall protein production rates in *E coli* decrease significantly at lower temperatures.^36^ In this study, the general protein production rate at 18°C across all constructs was only between 20%-50% of that observed at 37°C (**Figure S10B**). Consequently, an absolute increase in performance at lower temperatures was unexpected, as an OTS must compensate for the reduced protein production rate to achieve such an effect. Remarkably, for substrate **3** (at 0.3 mM),and substrates **4** and **5**, Mbur exhibited∼2.5 times higher performance at 18°C compared to 37°C. None of the other constructs showed similar absolute performance increase at lower temperatures, except for *Mm*, which displayed slightly improved performance for ncAA **3** at 1 mM.

### Investigating the substrate promiscuities of selected +N variants

As shown in **Figures 1** and **4**, the ΔN variants exhibit lower substrate promiscuity compared to the +N class, which is why this analysis focuses exclusively on the +N variants. Psychrophilic enzymes are often characterized by lower substrate affinity and specificity, attributed to their increased flexibility, or in some cases enhanced active site accessibility. These features result in greater substrate promiscuity, a common trait among psychrophilic enzymes.^23^

There is even an example of an engineered aaRS with increased flexibility which induced higher promiscuity compared to other engineered mutants with lower flexibility.^37^ From an enzyme engineering perspective, higher promiscuity is highly desirable as it facilitates the engineering of enzymes for new substrate recogntion.^38,39^

Our fluorescence heatmaps (**Figure 2**) indicated that the Mbur variant may also exhibit higher promiscuity, enabling more efficient incorporation of the tested ncAAs, compared to other variants. Since all PylRS variants display some degree of promiscuity, we sought to quantify this by deriving corresponding promiscuity scores. A standard method to determine enzyme promiscuity is to compare *in vitro* parameters for new substrates with those of the native substrate.^40^ However, since OTSs are complex systems were *in vitro* parameters might not clearly reflect *in vivo* efficiency^10^, we adapted the concept of *in vitro* enzyme promiscuity determination to cell-based *in vivo* scores (**Figure 5A**). In this approach, incorporation efficiencies for all substrates were normalized to the best-performing substrate for each ncAA concentration and mapped onto a scale from 0 to 100 to yield the substrate promiscuity scores. A perfectly promiscuous enzyme would attain a value of 100, indicating equal incorporation efficiency across all substrates. Conversely, a perfectly specific enzyme recognizing only one substrate would have the lowest possible score of 0. Among the functional constructs, the lowest promiscuity score in **Figure 5A** was 14.5, observed for Ml at 0.1 mM, while the highest score was 57.2 for SmbP-Mb, followed closely by 56 for Mbur.

**Figure 5.**
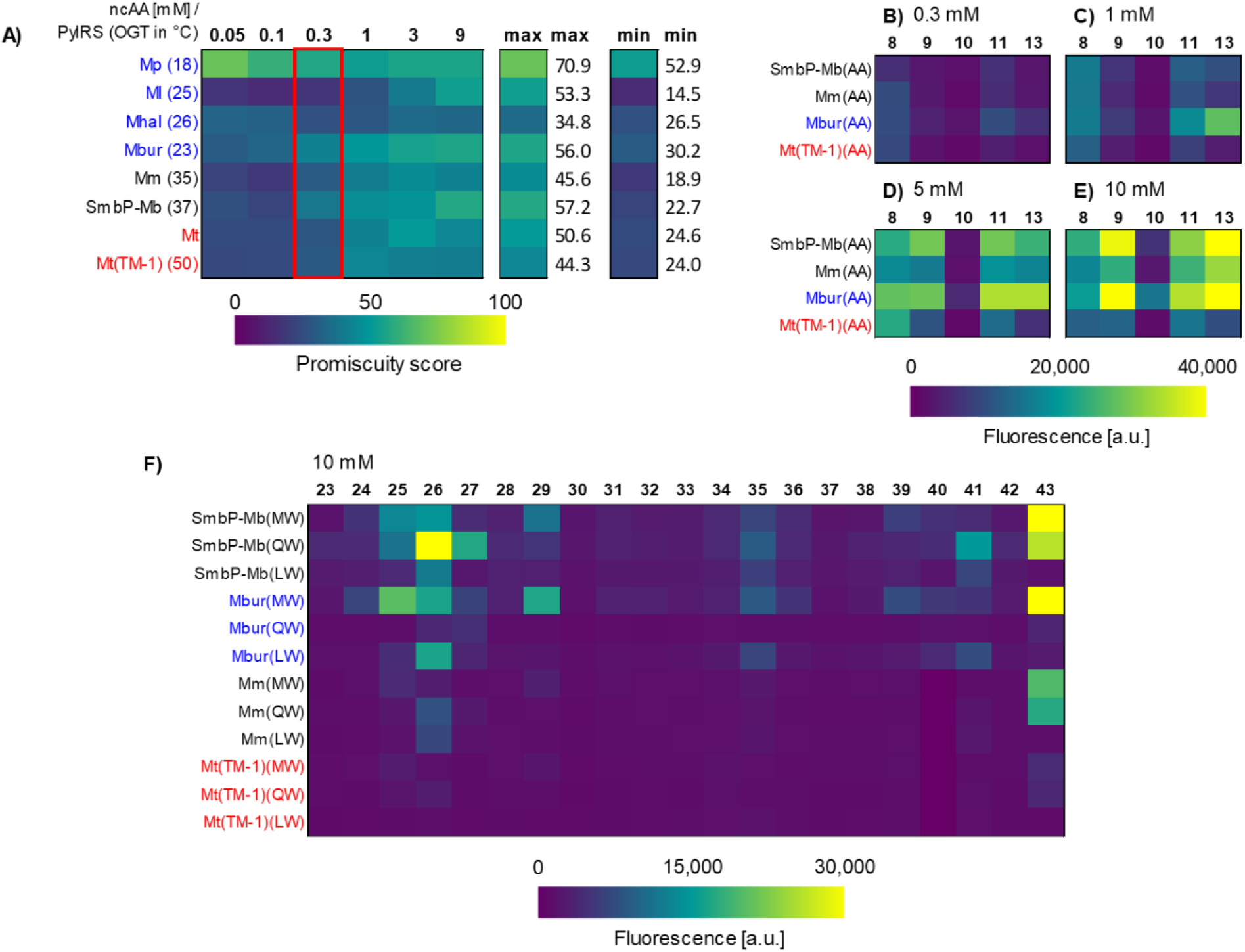
**A)** Heat maps showing the promiscuity score for each PylRS construct at given ncAA concentrations, with the maximum (max) and minimum (min) promiscuity scores in an additional column. The optimal growth temperature (OGT) of each PylRS construct is indicated in brackets. The ncAA concentration of 0.3 mM is highlighted; see text for a detailed explanation. **B), C), D), E)** Heatmaps based on fluorescence of sfGFP(1x amber) expression in *E. coli* BL21(DE3) at ncAA concentrations 0.3, 1, 5 and 10 mM. **F)** Selected PylRS mutants tested with 10 mM ncAA.Figures showing the number of replicates and error bars are provided in the supplements; for **B), C), D), E), Figure S40-S44** and for **F) Figure S45-S53**. Letters in brackets indicate the N311 and the C313 mutations in *M. barkeri* notation. Substrate numbering is detailed in **Figure 1**. Growth for the Mm and Mt(TM-1) constructs with substrate **35** was extremely low, so these values were excluded. Background suppressions were construct dependent and ranged from 1,100 to 2,500 [a.u.].

Unfortunately, data on in-cell ncAA concentrations are generally sparse. However, **2** and **3** show over 90% cell uptake from the medium, for concentrations up to 10 mM.^41^ Given that substrate **2** is highly hydrophobic and substrate **3** very hydrophilic, this provides a useful estimate for the uptake efficiency of the remaining ncAA set. These findings suggest that ncAA uptake is generally not a limiting factor for the Pyl analogs used in this study.

Mp showed the highest promiscuity score at low ncAA concentrations but was unfortunately barely functional under these concentrations (**Figure 2**). No clear trends were observed between promiscuity and the thermal origin of the different PylRS variants. As expected, the promiscuity score increased with rising ncAA concentrations (**Figure 5A**). This is logical, as both good and bad substrates reach their activity plateaus at different increasing concentrations, as reflected in our data (**Figure 2, 3** and **S5**). Therefore, a high promiscuity score at low ncAA concentrations indicates higher general promiscuity, as the activity plateau has not yet been reached. Notably, Mbur and SmbP-Mb exhibited the highest promiscuity scores at low ncAA concentrations (0.3 mM, **Figure 5A**, red box).

To evaluate the predictive power of the promiscuity score, we selected the two OTS variants with the highest promiscuity score (Mbur and SmbP-Mb) at 0.3 mM ncAA concentration and compared their performance to lower scoring variants (Mm and Mt(TM-1)). We used reported mutations for recognizing Phe and Tyr analogs, as well as small aliphatic ncAAs, and transplanted them into the chosen PylRS variants. The mutations responsible for recognizing Phe and Tyr analogs (**7-22**), referred to as AA, were N346A:C348A and were originally reported for Mm^42^ Similarly, the mutations responsible for recognizing small aliphatic ncAAs (**23-43**) included N311M:N313W, N311Q:N313W, N311L:N313W referred to as MW, QW, and LW, respectively, and were reported for Smbp-Mb^43^. Additionally, we tested special-case variants for S-allyl-cysteine (**43**) and S-propargyl-cysteine (**40**), which originated from SmbP-Mb and carried the mutations C313W:W382T.^43,44^

After an initial pre-screen of the Phe and Tyr analogs (**Figure S11**), the ncAAs with good activity were further screened in a concentration-depend manner to gain deeper insights (**Figure 5B-E**). In these tests, Mbur and SmbP-Mb demonstrated the best performance, with Mbur being more effective at lower ncAA concentrations, aligning well with the derived promiscuity scores. Notably, Mbur successfully incorporated **10**, which none of the other PylRS were able to incorporate.

A similar pattern emerged for the second class of ncAAs (**Figure 5F**). While most of the tested ncAAs exhibited relatively low activity, Mbur and SmbP-Mb consistently outperformed the Mm and Mt(TM-1) variants for those ncAAs with detectable activity, again corresponding well to the promiscuity scores.

Given that psychrophilic enzymes, including Mbur, often contain a higher proportion of amino acids with polar or charged side chains, we hypothesized that this might provide an advantage in recognizing ncAAs with similar properties. To test this, we evaluated substrates **18-22** using the AA variants shown in **Figure 5B-E**. Mbur was able to incorporate ncAAs **18-21**, with sulfotyrosine being incorporated most effectively. In contrast, SmbP-Mb produces weaker signals for substrates **18** and **19**, while Mm and Mt(TM-1) failed to incorporate any of these substrates (**Figure S12**).

Since these analogs are most likely synthesized by Tyr derivatization, the purchased ncAAs likely contained some degree of Tyr contamination (up to 5% for substrates 18–22). Therefore, we tested Tyr at various concentrations as a control. Assuming a maximum of 0.5 mM Tyr contamination at a 10 mM substrate concentration, we supplied up to 1.5 mM Tyr to the cultures. However, no Tyr incorporation was detected.

For substrates **40** and **43**, Mbur outperforms the other variants at lower ncAA concentrations and with multiple stop codons, even reaching wild-type levels of sfGFP production (**Figures S13**&**16**).

Overall, more ncAAs served as substrates for Mbur and SmbP-Mb than for Mm and Mt(TM-1) underlining the greater promiscuity of the former two and aligning with our promiscuity scores. In conclusion, our data suggests that the promiscuity scores could be useful in guiding future engineering projects, consistent with findings reported for other enzymes.^45,46^

### Comparing the Mbur variant to other state-of-the-art OTS

To gain insight into how Mbur compares to one of the most efficient OTS currently known for *E. coli*, we performed a side-by-side comparison with the *Mj*TyrRS OTS in an identical plasmid setup (pTECH).^16,47,48^ This comparison included two *Mj*TyrRS variants, selected after a substrate pre-selection to identify the best substrates (**Figure S17** and **S18**).

The first variant, *Mj*ONBYRS, developed by our group, is known for its exceptional efficiency.^47^ The second variant, *Mj*pCNFRS, was engineered to recognize *para*-cyano-phenylalanine and also accommodates various *para*-substituted phenylalanine derivatives.^49^ For Mbur, we selected the two most efficient Mbur variants for the two different substrates, **2** and **43** (**Figure 6**).

**Figure 6.**
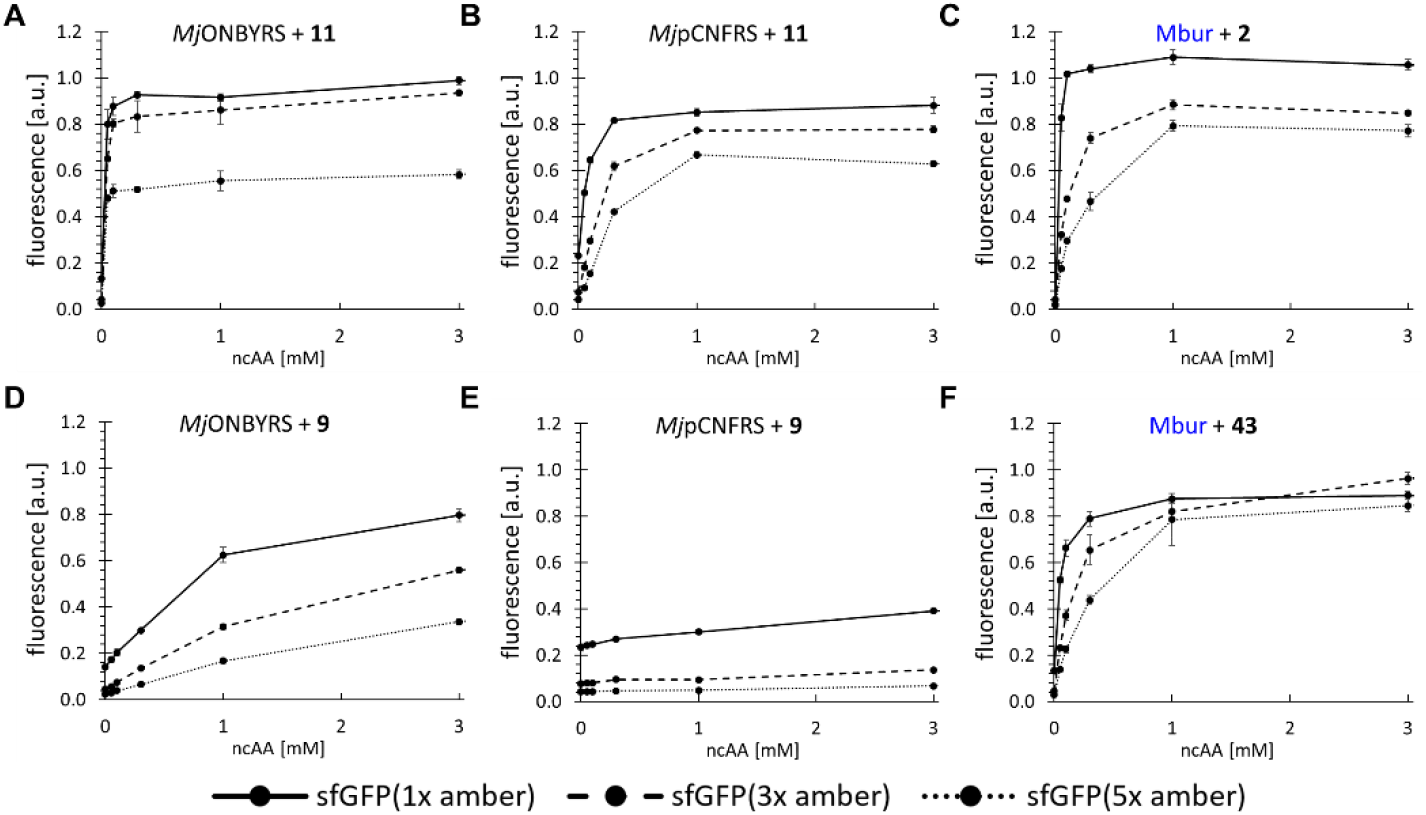
Concentration-dependent production of unnatural proteins by various OTS constructs: **A**) *Mj*ONBYRS with substrate **11, B**) *Mj*pCNFRS with substrate **11, C**) Mbur with substrate **2, D**) *Mj*ONBYRS with substrate **9, E**) *Mj*pCNFRS with substrate **9, F**) Mbur with tRNA from *M. alaskense* and substrate **43, H**) *Mm*PylRS with substrate **3**. Protein production was performed in *Escherichia coli* B-95.ΔA^35^. Endpoint measurements were taken at ncAA concentrations of 0.05, 0.1, 0.3, 1 and 3 mM. Fluorescence values were normalized to those of wild-type sfGFP reporter constructs (without an in-frame stop codon). Error bars represent the standard deviation of three replicates (n=3).

Later in our study, the genome of the psychrophilic organism *M. alaskense* (Mala) became available, allowing us to test the Mala tRNA (**Figure S62**) with Mbur for substrates **40** and **43** (**Figure S19**). Remarkably, this combination increased the incorporation efficiency up to 40% and was therefore included in the results shown in **Figure 6F**.

Overall, the best *Mj*TyrRS (**Figure 3A** and **B**) and Mbur (**Figure 3C** and **F**) systems performed comparably. Although wild-type PylRS systems have occasionally demonstrated similar performance levels^50^, this study is the first to report a mutated PylRS OTS capable of incorporating a non-Pyl derivate, **43**, with such high efficiency (**Figure 3F**), even at very low ncAA concentrations (0.3 mM).

### Validation of small-scale fluorescent assays by scaled-up protein production and purification

The protein yields obtained from scaled-up production and purification (**Figure 7**) aligned well with the trends observed in the small-scale fluorescence experiments, confirming the reliability of these assays. However, an exception was noted for the incorporation of substrate **40**, where the yield of target protein with Mbur exceeded our expectations (**Figure S13B**). Despite this anomaly, within each dataset, fluorescence values and protein yields correlated strongly.

**Figure 7.**
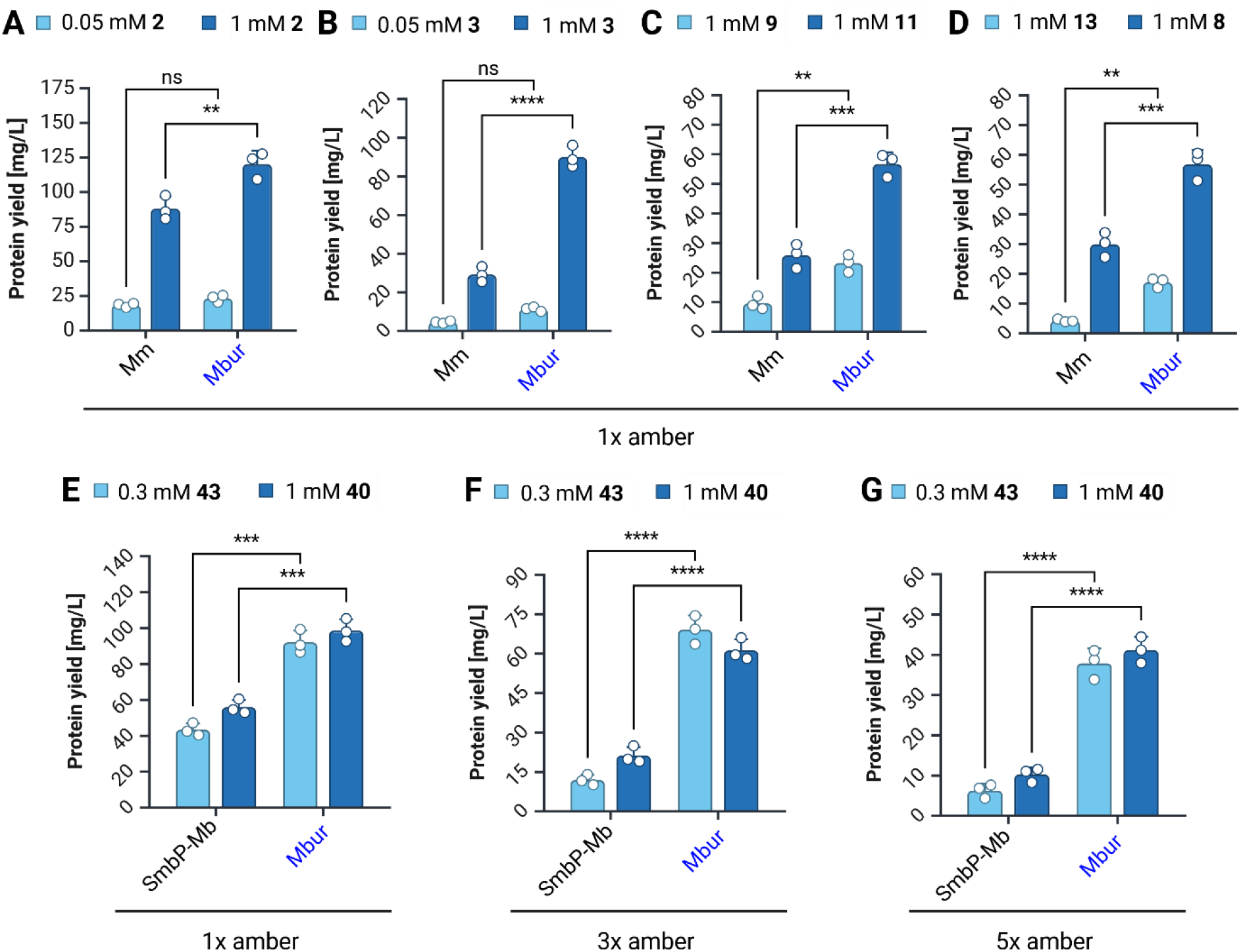
Protein yields of sfGFP from small-scale shake flask cultures. Protein yield is reported as the amount of protein per liter of cell culture. In the upper row *E. coli* BL21(DE3) was used with the indicated reporter constructs, while the bottom row displays results for *E. coli* B-95.ΔA with the indicated reporter. Substrate concentrations are provided in the legends, with numbering corresponding to **Figure 1**. For detailed information about the PylRS variants used and ESI-MS of purified proteins see **Tables S1** and **S2**. Statistical analysis was performed using a Two-way ANOVA with Tukey multiple comparisons test; significance is indicated by asterisks.

Improvements in the incorporation of a single in-frame ncAA ranged between 25% (for 1 mM **2**) and 200% (for 1 mM **3**) (**Figure 7A-4D**). The most remarkable improvements were observed in the multi-site incorporation of five ncAAs (**38** and **35**), with an improvement of over 400% and 300%, respectively (**Figure 7G**). A clear trend emerged, indicating that the efficiency difference between SmbP-Mb and Mbur increases with the number of ncAAs that are incorporated within a target protein (**Figure 7E-4G**).

To estimate the robustness of the 96-well plate assays, we analyzed the small scale 96-well plate sfGFP fluorescence data in dependence to the isolated protein yields of the shaking flask cultures by linear regression and derived the R^2^ values (see supplements **Figure S92**). The analysis revealed a notably strong correlation between the 96-well plate assays for each ncAA, with R^2^ values consistently exceeding 0.97. Even when combining data from cultivations with different ncAAs, the R^2^ values remained high, reaching at least 0.85, indicating a strong correlation of small-scale fluorescence and shake flask protein yields.

However, the combined R^2^ values were slightly lower compared to those from a single ncAA. This suggests that protein yields are influenced by factors beyond the specific ncAA concentration, potentially including ncAA toxicity leading to growth impairment. The high correlation was expected, as our 96-well plate experiments consistently operated within the expected linear range of the measurement space, with cell densities never exceeding OD_600_ = 1. Collectively, these results demonstrate the robustness of the data and indicates that protein yields from 96-well plate assays are reliably translatable to shake flaks cultivations.

## Discussion

Our work demonstrates that psychrophilic enzymes, such as the Mbur variant, offer promising advancements in OTSs. Mbur outperformed established systems like Mm and Alv in both single and multi-site ncAA incorporations, even under challenging conditions such as low ncAA concentrations and reduced cultivation temperatures. It also efficiently incorporated non-Pyl derivatives, showcasing exceptional substrate promiscuity and robustness. These findings validate the use of promiscuity scores for guiding future OTS engineering efforts and open new avenues for applications in synthetic biology and protein design.

We embarked on this study driven by the question of how to address the general issue of low *in vivo* efficiency of the PylRS orthogonal translation system. To tackle this, we conducted a comparative study exploring +N PylRS homologs derived from extremophilic organisms and included 15 ΔN class variants, resulting in a total set of 24 OTSs. Among these, we identified a highly active variant from the psychrophilic (cold-active) organism *Methanococcoides burtonii* (Mbur). Mbur exhibited exceptional efficiency in incorporating various ncAAs into proteins, especially at low temperatures (**Figure 4**).

The superior incorporation efficiency was consistently observed for all major substrate classes, demonstrating the robustness of the results. Particularly remarkable was the multi-site ncAA incorporation with BocK (**2**) and especially S-allyl-cysteine (**43**). To the best of our knowledge, no previous PylRS mutant has achieved such high ncAA incorporation efficiency for non-Pyl-analogs, approaching that of the wild-type reference. This finding is significant, as it has the potential to address the challenges posed by the high cost of certain ncAAs, thereby facilitating and accelerating research projects in the field.

Considering that only two OTSs - *Mj*TyrRS^47,48,51^ and *Af*TyrRS^51^ - are known to approach wild-type recombinant protein production yields, the addition of a third OTS to this group is highly desirable. The inclusion of a third orthogonal translation system enhances the potential for simultaneous incorporation of a greater variety of ncAAs, both identical and different ones, with higher efficiency. The most promising improvements are expected in organisms with liberated codons, as the selection of tRNA^Pyl^ anticodons for orthogonal translation provides a high degree of flexibility.^52,53^ When the Mm is used in a recoded *E. coli* strain, OTS efficiency is higher for some of these newly created codons, but only to a limited extent.^53^ This result suggests that the *in vivo* efficiency of the PylRS OTS remains a bottleneck, even when free codons are available. In contrast, Mbur, demonstrates strong potential to overcome this limitation, as evidenced by its high performance in an RF1 deficient strain (**Figures 3, 4** and **6**).

The high performance of Mbur, even at very low ncAA concentrations, opens new possibilities for coupling with ncAA-producing metabolic pathways, potentially eliminating the need for external ncAA supply, as we recently elaborated.^2^ Efficient OTS activity at intracellular ncAA concentrations below 1 mM is critical, as most organisms maintain intracellular canonical amino acid concentrations below this threshold, and engineering pathways to allow for ncAA concentrations above 1 mM seems very unlikely.^54^

This high efficiency at low ncAA concentration also has further implications. The low cost and minimal concentration required for the efficient incorporation of **43** allow recombinant proteins containing one or even multiple bioorthogonal alkene functions to be produced at wild-type levels with negligible additional cost. The alkene function can be perfectly addressed with the most advanced bioconjugation reaction, the inverse electron demand Diels–Alder (IEDDA) reaction, using a tetrazine moiety.^55^ More generally, click reactions will be greatly facilitated as 17 of the 43 incorporated ncAAs (**Figure 1**) are crucial precursors for establishing covalent links in otherwise transient complexes of biomolecules.

Enzyme promiscuity, while unpredictable, is a crucial property in enzyme engineering.^45,46^ Thus, our promiscuity coefficients could serve as useful indicators of the engineering potential of a PylRS variant. We demonstrated that Mbur exhibits higher substrate promiscuity compared to other PylRS. This is evident from its superior performance across all tested substrate classes, even though the mutations were directly transplanted from reported PylRS variants for these substrates without any additional engineering or optimization.

While it is generally possible to transfer mutations from one PylRS ortholog to another to enable recognition of a specific ncAA^56–58^, it is unusual - and highly unlikely - for the recipient variant to also outperform the donor variant without further engineering^59^. Remarkably, this was observed with the Mbur variant, underscoring its exceptional promiscuity. Since substrate promiscuity is a strongly desirable trait in enzyme engineering, as it facilitates the engineering of recognition for new substrates, Mbur shows great promise as a candidate for further expanding the ncAA repertoire, unlocking new possibilities for advancements in this field.^45,60–62^

### Outlook

Our discoveries suggest that shifting the search for efficient orthogonal translation systems to psychrophilic organisms could be highly advantageous. Psychrophilic enzymes, characterized by their flexible architecture, maintain functionality at low temperatures due to structural adaptations such as reduced hydrophobic core packing, elongated loops, fewer hydrogen bonds, and less dense salt bridges.^63^ This flexibility is linked to substrate promiscuity, which is crucial for experimental evolution, driving the emergence of new enzyme specificities and functions.^64^ Although relatively few psychrophilic enzymes have been extensively studied^64^, viral proteins exhibit similar features, such as elongated loops and less densely packed cores ^65^. Tokuriki et al. developed a mutational gradient robustness model for viral proteins, demonstrating lower fitness costs per mutation compared to thermophilic proteins, thereby enhancing mutational tolerance and adaptability.^65,66^ These characteristics make psychrophilic enzymes and viral proteins excellent models for studying evolutionary innovation and the emergence of novel protein functions.

From a biocatalysis perspective, psychrophilic enzymes exhibit remarkable plasticity in substrate recognition due to their structural adaptations, which enhance catalysis at low temperatures.^23^ Features such as flexible and accessible active sites, smaller side chains, and distinct electrostatic potentials facilitate efficient substrate binding, reduce activation energy, and improve catalytic turnover. Cold-active enzymes often show increased Km values and lower ΔG‡, optimizing reaction rates by loosening substrate binding while stabilizing the transition state. Additionally, the higher entropic contributions (TΔS‡) in these enzymes reflect substantial structural ordering during catalysis. Notable examples include cold-active cellulases and proteases.^22,67^ Our findings align with these observations and underscore the potential benefits of leveraging psychrophilic organisms in the search for efficient OTSs, which could have far-reaching implications for biotechnology and evolutionary studies.

While most research in orthogonal translation has focused on engineering thermophilic and mesophilic aminoacyl-tRNA synthetases (aaRS),^68,69^ our study demonstrates the potential of psychrophilic enzymes as a valuable source for highly efficient OTSs. These properties are not limited to aaRSs and could extend to other components of the host translational machinery, including orthogonal ribosomes or elongation factors.^70–72^ By leveraging the inherent benefits psychrophilic macromolecules - particularly their higher promiscuity - future engineering efforts could unlock new possibilities for synthetic biology and protein design.

In our study, we identified a psychrophilic PylRS enzyme displaying typical properties of psychrophilic enzymes, such as heightened enzyme efficiency and promiscuity. These traits are crucial for synthetic biology and protein engineering, yet these enzymes remain underutilized beyond protease research for industrial applications. We hope our findings will inspire further exploration and utilization of psychrophilic enzymes across diverse scientific and industrial fields.

## Materials and Methods

### Canonical and Non-Canonical Amino Acids

Canonical amino acids were purchased from Carl Roth. Non-canonical amino acids were obtained from Fluorochem, Iris Biotech, Chempur, Sigma-Aldrich (Merck), Chiralix, Toronto Research Chemicals, Carl Roth, ThermoFisher Scientific and TCI Deutschland (see Table S4).

### Plasmid Vector Construction, PylRS and Reporter Sequences

The pTECH vector was a gift from Dieter Söll (Addgene plasmid #104073).^11^ Genes were ordered as codon optimized fragments from Twist Bioscience. All plasmids were assembled by Golden Gate cloning and confirmed by Sanger sequencing. Point mutations were introduced by non-overlapping inverse PCR. All PylRS and reporter sequences used in this study are provided in the supplementary materials.

### Analysis of SUMO-sfGFP Expression by Intact Cell Fluorescence

*E. coli* BL21(DE) cells were used for the small-scale expression of reporter constructs. Electrocompetent cells were transformed with the orthogonal translation system and reporter plasmids. LB agar plates for plating contained 1% glucose and corresponding antibiotics. Single colonies of clones were used to inoculate 2 mL of LB medium (in 14 mL tubes) with 1% glucose and appropriate antibiotics, and cultures were grown to saturation overnight. Assays were conducted in 96-well plate format. Cultures were added to each well at 1:100 dilution in ZYP-5052 auto-induction medium to a final volume of 100 μL, supplemented with antibiotics and ncAAs. Cells were grown in black μ-plates (Greiner Bio-One, Kremsmünster, Austria) covered with a gas permeable foil (Breathe-Easy^®^, Diversified Biotech, Dedham, MA, USA) with orbital shaking for 24 h at 37 °C. For endpoint measurements (Tecan M200, Männedorf, Switzerland), the plate foil was removed, and fluorescence was measured with an 85 gain setting. For OD_600_ measurements, 50 μL of ZYP-5052 medium was pipetted into clear 96-well μ-plates, and 50 μL of culture was added. Excitation and emission wavelengths for fluorescence measurements were set to 481 nm and 511 nm, respectively. Fluorescence values were normalized to the corresponding OD_600_.

### Temperature Dependent *In Vivo* Performance and Flow Cytometry

Cells were transformed with the orthogonal translation system, reporter plasmids and necessary controls. LB agar plates used for plating contained 1 % glucose and corresponding antibiotics. Single colonies of clones were inoculated into 2 mL LB (in 13 mL tubes) supplemented with 1 % glucose and the appropriate antibiotics and grown to saturation overnight. Subsequently, 4 mL ZYP-5052 medium, supplied with antibiotics and the desired ncAA, was inoculated (1:50) and grown to an OD_600_ = 0.4-0.6 (∼5 hours). The culture was then split into 2x 2mL aliquots: one was incubated at 18°C and the other one at 37°C. After 24 hours, cells were diluted to an OD_600_ of ∼0.025 using sterile filtered PBS (0.8 μL of the culture in 500 μL PBS) and analyzed by flow cytometry. For cultures incubated at 18°C, this procedure was repeated at additional timepoints of 48h and 72h

Cytometric analyses were conducted using an Attune NxT flow cytometer (ThermoFisher Scientific, Waltham, MA, USA) equipped with 488 nm and 561 nm lasers. The fraction of cells containing the desired and reporter proteins was determined using a gating strategy targeting single cells while excluding aggregates and other background noise.

For the cultures grown at 18°C, the maximum value observed within the 24-72h interval was used for **Figure 4**.

### Protein Expression and Purification

For expression of SUMO-sfGFP variants, *E. coli* strains were used in 10 mL ZYP-5052 medium supplemented with indicated ncAA concentrations and appropriate antibiotics. The expression medium was inoculated with a starter culture (1:100). Shake flasks were incubated for 24 h at 37 °C while shaking at 200 rpm. Cells were harvested by centrifugation and stored at –80 °C or directly used for protein purification.

Harvested cell pellets were resuspended (50 mM sodium phosphate, 300 mM NaCl, 20 mM imidazole, pH 8.0) and lysed with B PER^®^ Bacterial Protein Extraction Reagent (ThermoFisher Scientific, Waltham, MA, USA) according to their protocol, with addition of phenylmethanesulfonyl fluoride (PMSF, 1 mM final concentration), DNAse, and RNAse. Cleared lysates were loaded onto a equilibrated Ni-NTA column and purified via the P-1 peristaltic pump (Pharmacia Biotech, now Cytiva, Marlborough, MA, USA) or by gravity flow. After washing with 10 column volumes of resuspension buffer, elution buffer (50 mM sodium phosphate, 300 mM NaCl, 500 mM imidazole, pH 8.0) was applied to elute the his-tagged target proteins. The first 1.5 mL (1 mL for gravity flow) covering the void volume was discarded. Afterwards, the eluate (1 mL) was collected and dialyzed in cellulose film tubings against 1 L buffer (50 mM sodium phosphate, 300 mM NaCl, pH 8.0) for at least 2 h with three buffer changes. Concentrations of purified reporter proteins were determined by measuring the sfGFP chromophore absorption at 488 nm.

### Intact Protein ESI-MS

Intact protein mass measurements of purified SUMO-sfGFP variants were performed by electrospray LC-MS on a Waters H-class instrument with a Waters Acquity UPLC protein BEH C4 column (300 Å, 1.7 μm, 2.1 mm × 50 mm). Gradient elution went from 5% MeCN in aqueous 0.01% formic acid to 95% in 6 min at a flow rate of 0.3 mL/min. Mass analysis was conducted with a Waters Xevo G2-XS QTof analyzer. Proteins were ionized in positive ion mode, applying a cone voltage of 40 kV. Raw data were analyzed using the maximum entropy deconvolution algorithm. The data were exported and plotted with QtiPlot (version 0.9.9.7).

## Acknowledgments

This work was performed as part of the “Site-directed cross-linking of KLK proteases from prostate” project funded by the Lead Agency FWF (I 3877-B21) (P.G.) with DFG-D-A-CH (BU1404/12-1) (N.G.K). N.B. thanks Canada Research Chairs Program (Grant Nr. 950-231971) and the Natural Sciences and Engineering Research Council (NSERC) of Canada through the Discovery Grant (RGPIN-05669-2020) for support. This work was supported by the Cluster of Excellence “Unifying Systems in Catalysis” (UniSysCat), funded by the Deutsche Forschungsgemeinschaft (DFG, German Research Foundation) under Germany’s Excellence Strategy–EXC 2008/1–390540038 (N.B. and J. R.). We are very grateful to Christian Stieger (Leibniz-Forschungsinstitut für Molekulare Pharmakologie, Hackenberger Group) for his support in ESI-MS measurements. A portion of the used data were produced by the US Department of Energy Joint Genome Institute (https://ror.org/04xm1d337; operated under Contract No. DE-AC02-05CH11231) in collaboration with the user community. Special thanks go to my bench neighbor Dr. Georg Johannes Freiherr von Sass for inspiration to take this journey.

## Notes

### Competing Interest Statement

N. G. Koch and N. Budisa have filed a european patent application (EP4328308) for this orthogonal translation system.

### Summary of Updates

16 constructs of ΔN PylRS have been included into the test set (Figure 1 and Figure S6). Additional data for multi-site ncAA incorporation was added (Figure 3). Temperature dependent incorporation efficiency was determined via flow cytometry for the 9 best PylRS variants (Figure 4). Several charged ncAAs were tested for incorporation with the known double Ala mutant (Figure S12). Comparison of pUltra vs pTECH setup was added to the SI (Figure S7).

