## Supplementary Information for "“Cold” Orthogonal Translation: A Psychrophilic Pyrrolysyl-tRNA Synthetase Boosts Genetic Code Expansion in *E. coli*"

### 1. Supplementary Results

#### 1.1 Phylogenetic PylRS Analysis

Initial sequencing discrepancy: Phylogenetic analysis was performed for all *Methanosarcinales* and *Methanomassiliicoccales* PylRS sequences (**Figure 1**). These sequences were sourced from genome data from NCBI and for *M. alaskense* from the JGI database. In addition, the sequence of *M. thermophila* was included, which was determined with degenerative primer amplification from genomic DNA before the whole genome was available.[1] Unexpectedly, the sequence of this PylRS did not match the PylRS sequence that was extracted from the later published genome of *M. thermophila* (strain TM-1). Since there are two other sequences also extracted from genomes of different strains (CHTI 55 and MT-1) that match the TM 1 sequence, this is most likely the correct one and is referred to as *Mt*PylRS(TM 1) in this work. The other *Mt*PylRS appears to be more closely related to the only known psychrophilic *Methanosarcinae* PylRS from *M. lacustris* and was therefore retained in this study.

Selection criteria for psychrophilic candidates: As shown in the phylogenetic tree (**Figure 1A**), there are six potential psychrophilic candidates With at least one psychrophilic PylRS selected from each genus. When more than one variant was selectable, the one from the species with the lowest optimal growth temperature (OGT) was used.

Choice of variants: The decision to include the PylRS variants from *Methanococcoides burtonii* and *Methanococcoides alaskense* was made based on their high sequence identity. As for the genus *Methanococcoides*, only the genome of *M. burtonii* was available when the project was planned in December 2019 and was therefore used. In hindsight, the decision on which construct to include would have been the same, as these two psychrophilic PylRS variants from *M. burtonii* and *M. alaskense* have more than 95% sequence identity, but a closer look at the tRNAs revealed clear differences in free energy (**Figure S59**). The free energy of $tRNA^{Pyl}$ from *M. burtonii* is almost 6 kcal/mol higher at 37 °C. This suggests that *M. burtonii* may be even more psychrophilically adapted, which is why this variant would have been chosen either way.

#### 1.2 Determination of the PylRS Variable Regions (Linker)

First, ColabFold[2] was used to predict a structure of one PylRS from each family. This prediction tool provides secondary structure predictions along with a confidence score (pLDDT) for the resulting structure as output. According to the original AlphaFold2 publication, a low score reliably indicates an unstructured region.[3,4] Utilizing these scores, the determination of the linker region between the N- and C-terminus was performed. This involved identifying regions with low confidence scores indicative of potential linker regions. We cross-checked the predicted start and end sequences of the linker region in each family alignment. In cases where the determined start and end sequences were not conserved across the family, additional structure predictions were carried out to refine the linker region. Typically, the C-terminal linker end was characterized by a conserved AA sequence. Minor variations were occasionally observed at the N-terminal start, resulting in occasional shifts in the linker beginning by one AA. The predicted AlphaFold2 structures used in this analysis can be found in Figures S3 and S4, with corresponding PDB files provided in the Supplementary Data.

##### 1.3 Analyses of the Amino Acid composition of PylRS

Clustering and arrangement into a heatmap: To analyze the AA composition for the PylRS of *Methanosarcina, Methanococcoides*, *Methanohalophilus* and *Methanolobus*, the PylRS without linker region and the variable linker region were clustered and arranged into a heat map (**Figure S1**). The heat maps display the percentage of AAs included and ratios of AAs classified according to their physico-chemical properties and the total length of the enzyme and linker.

Comparison of AA usage: Despite analyzing PylRS enzymes from different genera, there are no obvious trends in AA usage for the different PylRS. For example, the percentage of nonpolar AA is within a margin of 2.5%. All PylRS were found to be almost the same length when the linker region is removed within a margin of three AA. The analysis indicates that while there are no discernible patterns in AA usage among the PylRS enzymes studied, their lengths remain fairly consistent, at least when the variable linker region is excluded (vide infra).

Consistency in enzyme length: A more heterogeneous picture emerges for the linker region; therefore, the apparent differences are discussed. In general, the most abundant AA residues in the linker are consistent with empirical data on AA usage in intrinsically disordered proteins (IDPs). The seven AAs with the highest propensity in IDPs are P, E, S, Q, K, A, G (sorted by highest to lowest impact) which is corroborated by the data in **Figure S1**.[5] The largest difference of linker composition within this family is the proportion of K. The genera *Methanococcoides* and *Methanohalophilus* have a far higher K content than the other two groups. A higher K content could promote or fine tune linker-tRNA interactions because K is positively charged at physiological pH and RNAs are negatively charged molecules.[6] In addition, the linker of *Methanosarcina* contains much less acidic and basic AAs than the others. Overall, we calculated the intrinsic disorder propensities according to Theillet and colleagues and found that the calculated scores are relatively similar, despite the difference in AA composition.[5] This suggests that the length of the enzyme itself is relatively consistent across different species or genera.

Understanding the substantial differences in linker composition is challenging in the absence of structural data; however, all available evidence strongly indicates that this region is pivotal in fine-tuning enzymatic activity to suit the specific environmental conditions of the respective organisms. These tuning effects may extend to influencing the interaction between the respective PylRS and its corresponding $tRNA^{Pyl}$, as variations are observed at certain positions depending on the organism's origin within the family.[6] One study clearly shows that altering the linker length can have effects on wild-type and engineered PylRS variants, although the precise mechanisms underlying this phenomenon remain unclear.[7]

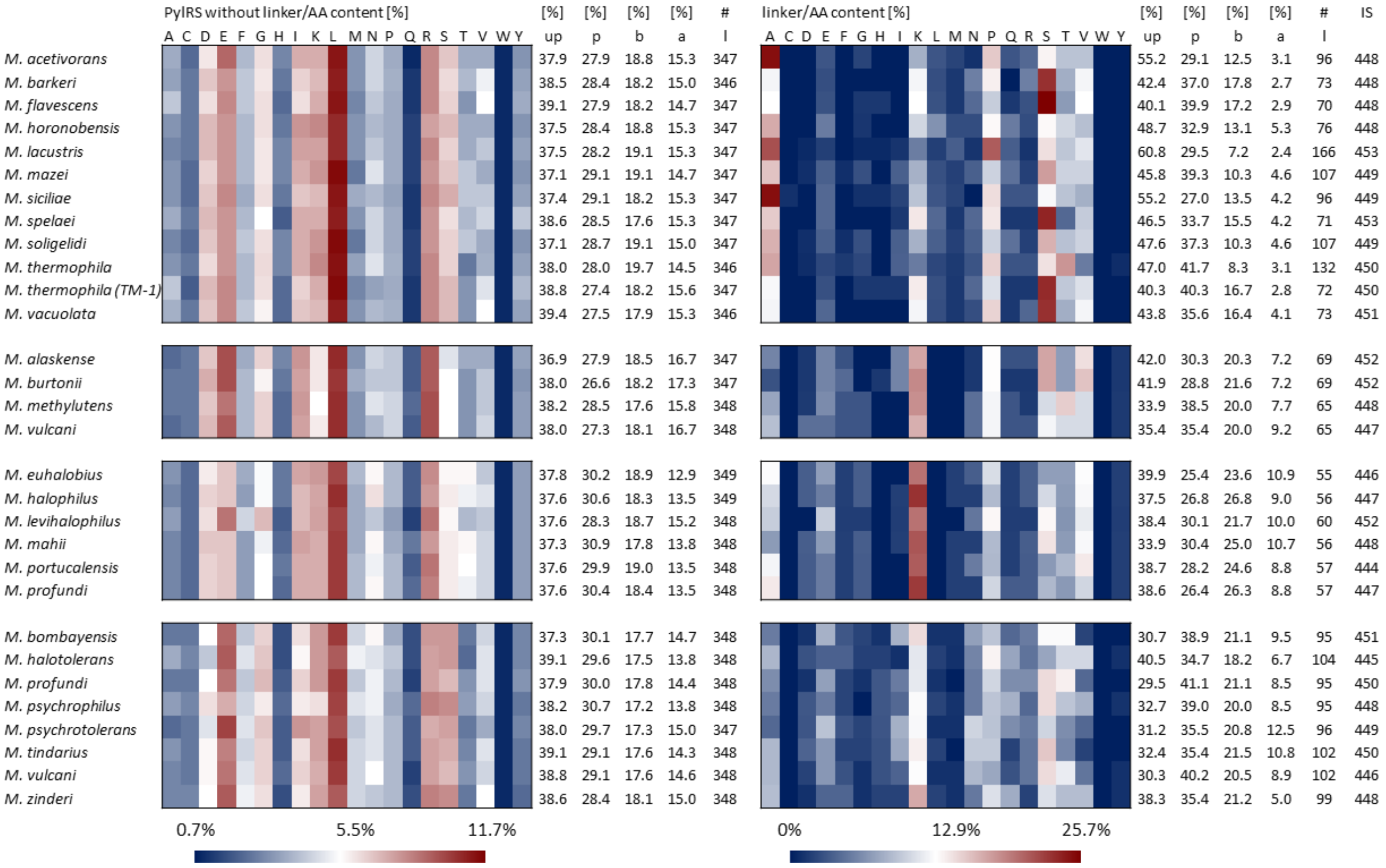

**Figure S1.** Amino acid composition of the PylRS enzymes without linker (left) and with linker (right). The first group is *Methanosarcina*, the second *Methanococcoides*, the third *Methanohalophilus* and the fourth *Methanolobus* (From top to bottom). Abbreviations are up = unpolar, p = polar, b = basic and a = acidic amino acids, in per cent; l = length of the enzyme/linker in number of amino acids; IS = IDP score.

##### 1.3.1 The special case of the PylRS linker of *Methanosarcina*

It is intriguing to note that upon removal of the linker, all PylRS enzymes exhibit nearly identical sequence length, differing only by a single amino acid (as depicted in **Figure S1**). A notable characteristic of *Methanosarcina* PylRS is also that the lengths of the variable region are much more diverse, in contrast to all other PylRS of the same family. In *Methanosarcina* the shortest linker spans 70 amino acids, while the longest extends to 166 amino acids (**Figure S1**, see below for details on variable region determination). This level of variability in linker length is unparalleled among genera within the same family.

Based on this observation, we hypothesized a potential correlation between the thermal origin and the linker length of *Methanosarcina* PylRS enzymes. To test this hypothesis, a sequence identity matrix was constructed based on the PylRS sequences (**Figure S2**). An alignment of the full-length enzyme was conducted, excluding the linker. This exclusion was warranted as it allowed for the establishment of a relationship between the remainder of the enzyme and the linker length. It is worth noting that significant sequence gaps were considered acceptable for a robust relationship estimation only under certain circumstances.[8]

If trends in temperature adaptation were present, they would likely manifest within the enzyme sequence and exhibit some degree of clustering. Based on these findings, we propose that linker length in PylRS enzymes predominantly influences catalytic activity tuning rather than the enzyme sequence itself, although the sequence's proline content may contribute to rigidity.

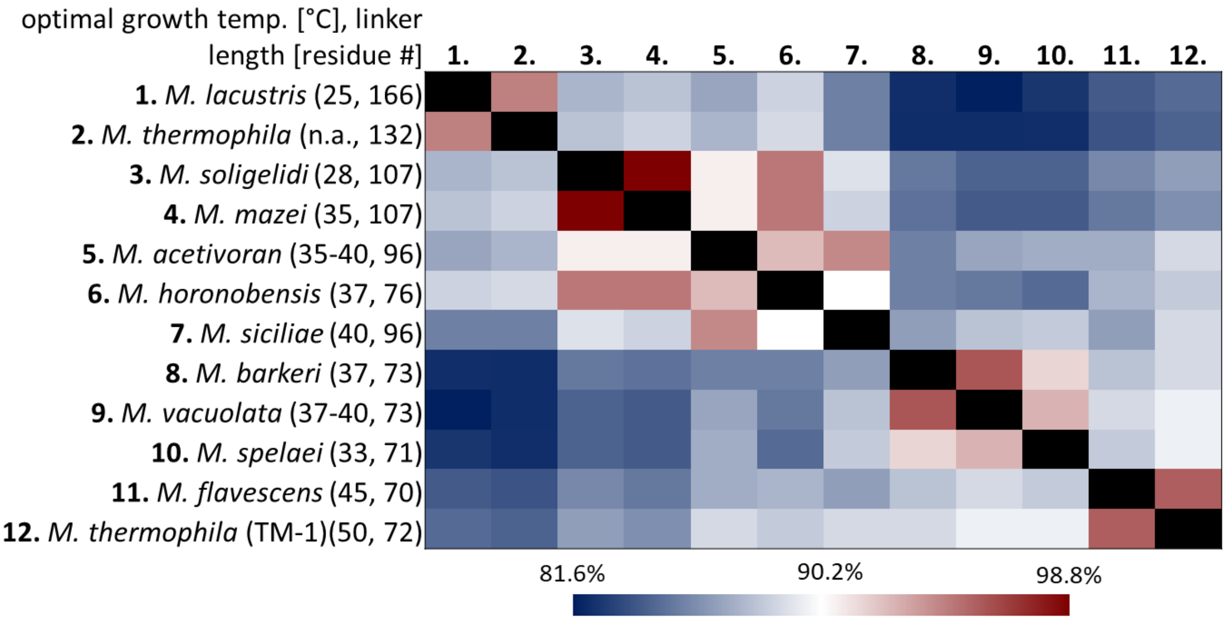

**Figure S2.** Sequence identity matrix of the *Methanosarcina* PylRSs without linker. [AA] = number of amino acids as linker, n.a. = not available.

The sequence identity matrix reveals correlations consistent with those observed in the phylogenic tree (**Figure 1A**). Examining the columns corresponding to *Ml*PylRS (**1.**) and *Mt*PylRS(TM-1) (**12.**), an inverse correlation between OGT and linker length becomes apparent, with the most psychrophilic variant having the longest linker. Given that a longer linker enhances flexibility, this observation aligns with the adaptation of the psychrophilic enzyme. Notably, the Pro content follows the same trend, with high proportions found in enzymes with longer linkers. Psychrophilic homologs typically exhibit higher flexibility of the enzyme due to longer linkers between domains. For example, a psychrophilic cellulase has been observed to have an unusually long linker of 100 AA, which is five times longer than that of the mesophilic homolog.[9,10] In the case of PylRS enzymes, the rise in Pro content accompanying the increase in linker length appears to counteract (mitigate?) this increase in flexibility to some extent.

Based on the accumulated evidence, it is highly plausible that the PylRS referred to as *Mt*PylRS in this work does not posses the AA composition and linker length characteristic of a thermophilic PylRS. This assertion is supported by the fact that *Mt*PylRS shares over 94% sequence identity with *Ml*PylRS, a homolog associated with psychrophilic adaptation. Therefore, despite its nomenclature suggesting thermophilicity, *Mt*PylRS likely exhibits features more consistent with enzymes adapted to colder environments.

#### 1.4 **Structural Biology Considerations on PylRS**

The predicted structures of *M. mazei* (magenta) and *M. burtonii* (red) were compared with the experimental structure of the C-terminus of *M. mazei* (PDB ID: 2Q7H, blue)[11] (**Figure S3A** and **B**). While the predicted structures are almost identical except for V401 the experimental structure deviates mostly for residues Y306 and N346. This difference is easily explained by the fact that the experimental structure was determined with bound adenylated pyrrolysine, and pyrophosphate and it is known that N346 interacts with the ester carbonyl O of pyrrolysine, suggesting an induced fit mechanism. In addition, the catalytic pockets of predicted structures of most of the Methanosarcina PylRS are almost identical, which agrees well with the highly conserved sequences in the Methanosarcinales order (data not shown).

To build a structural model of *M. burtonii* (**Figure 1B**), the crystal structure of the C-terminus of *Desulfitobacterium hafniense* (*D. hafniense*) PylRS (PDB ID: 2ZNI)[12] was superimposed with the

N-terminus (5UD5)[13] of *M. mazei,* with both bound to their respective tRNA$^{Pyl}$. Using the alignment, the N- and C-termini of *M. burtonii* (**Figure S4B**) were modeled based on this superposition (**Figure S4C)**, whereas the *Mbur*PylRS model is shown in **Figure 1B**.

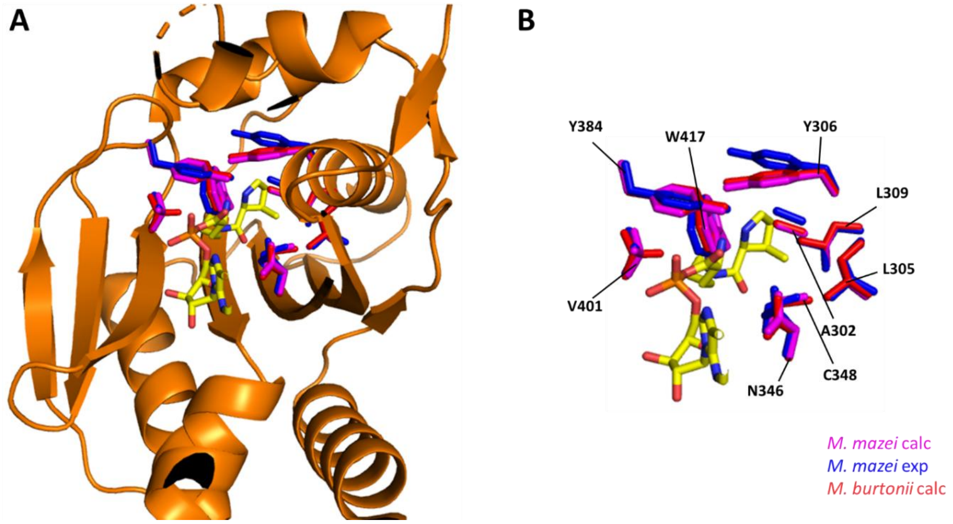

**Figure S3.** Overlay of the C-terminal domain of *M. mazei* (predicted = magenta, experimental determined = blue) and *M. burtonii* (red), with cartoon representation of the enzyme **A**) and without **B**).

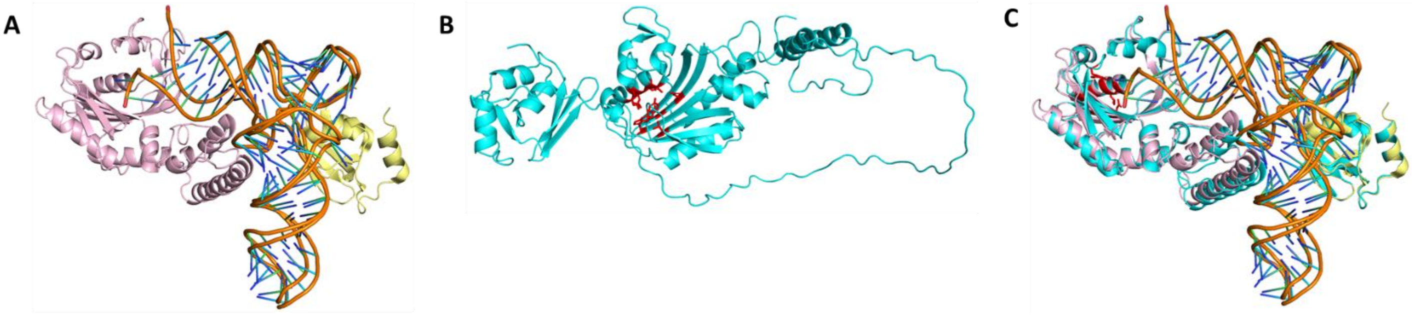

**Figure S4.** Experimental and predicted structures employed for the construction of the structural model depicted in **Figure 1B**. **A**) Superposition of C-terminus of *M. hafniense* (PDB ID: 2ZNI, light pink)[12] bound to tRNA$^{Pyl}$ with the N-terminus of *M. mazei* (light yellow) bound to tRNA$^{Pyl}$. **B**) Predicted structure of *M. burtonii* (cyan). **C**) Overlay of **A**) and **B**) with deleted linker.

#### 1.5 Complete +N and ΔN heatmaps

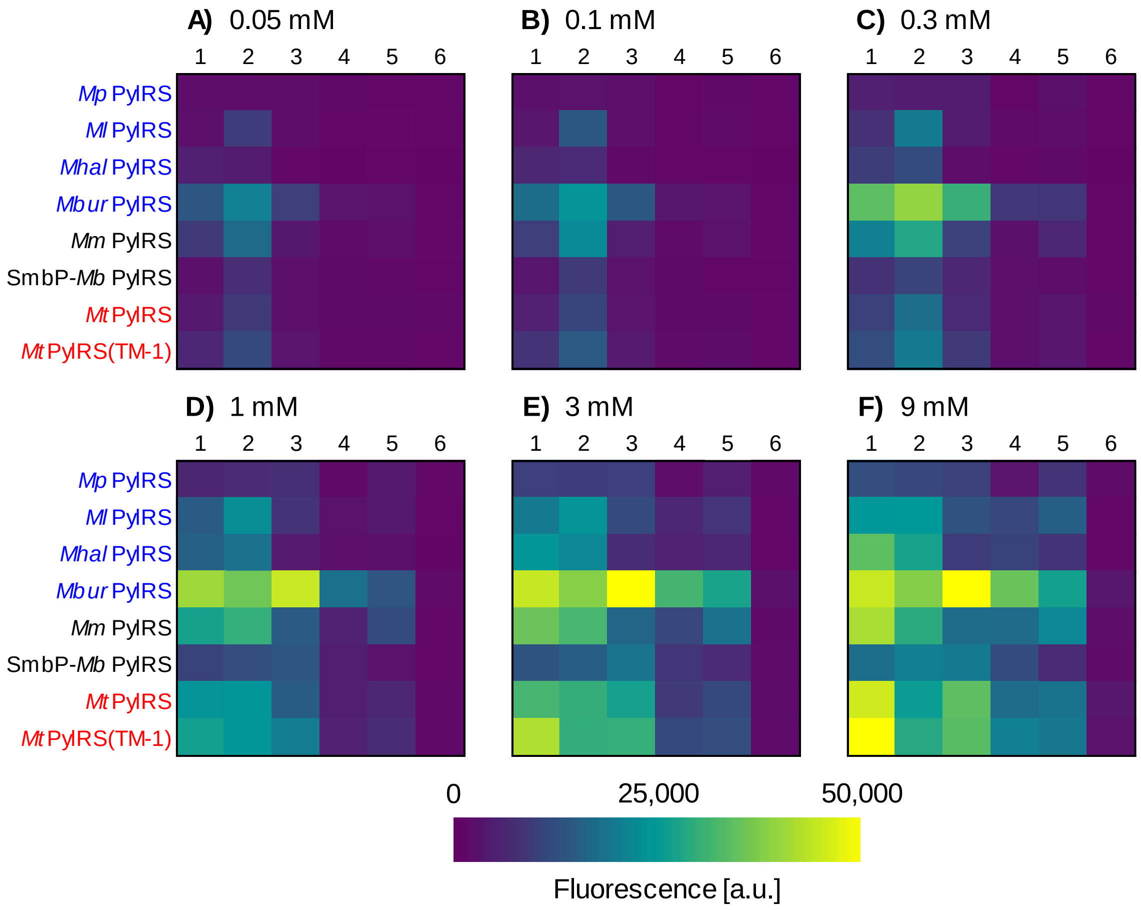

**Figure S5:** Heatmaps based on fluorescence of sfGFP(1x amber) expression with *E. coli* BL21(DE3) for the eight selected +N PylRS as a function of different ncAA concentrations; **A)**, **B)**, **C)**, **D)**, **E)**, **F)** are 0.05, 0.1, 0,3, 1, 3, 9 mM ncAA supplied. Bar charts with the number of replicates and error bars with the standard deviation can be found in the Supplementary Material (**Figures S17-S34**). Substrate numbering corresponds to that in **Figure 1**. Background suppressions were between 900 and 1,300 [a.u.] depending on the construct.

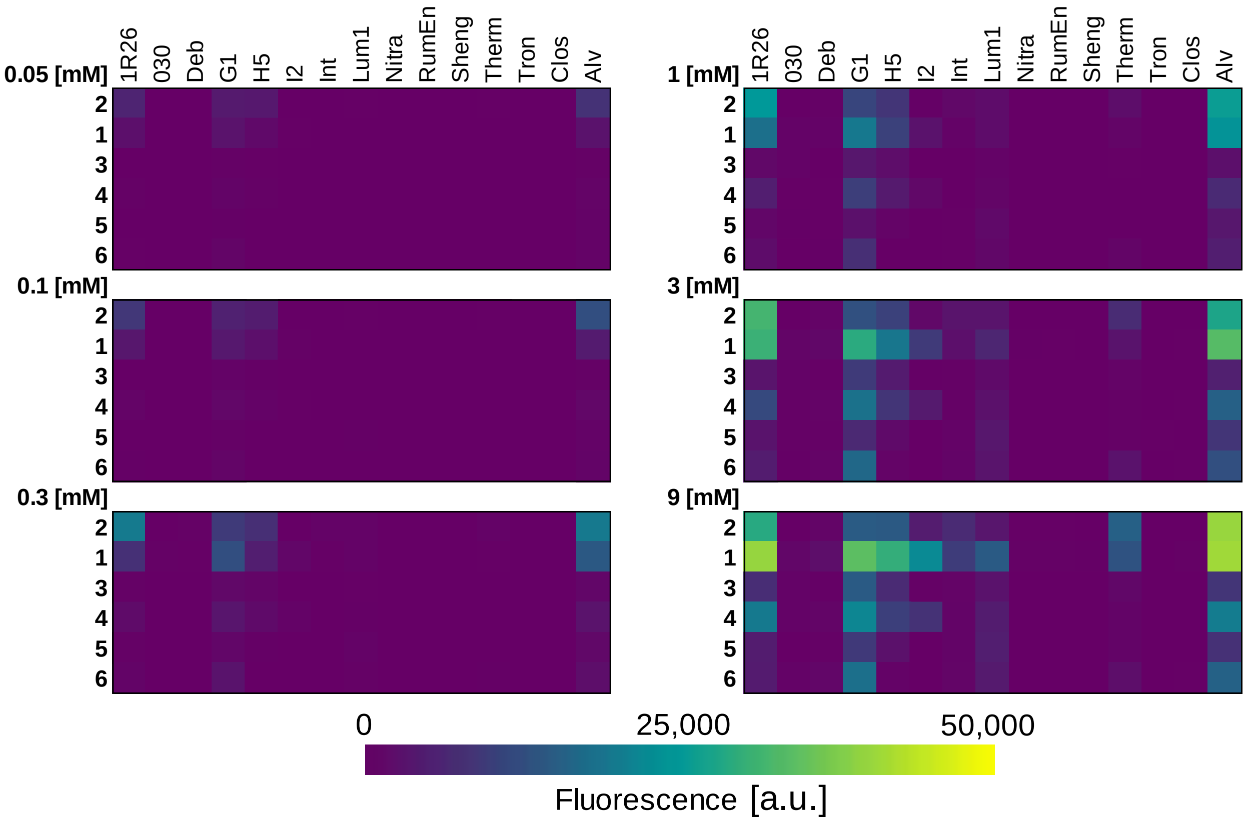

**Figure S6:** Heatmaps based on fluorescence of the sfGFP(1x amber) expression with *E. coli* BL21(DE3) and 15 selected ∆N PylRS variants. Substrate numbering can be found in **Figure 1**. Background suppressions were construct dependent and ranged from 200 to 400 [a.u.].

#### 1.6 **pUltra vs pTECH OTS setup**

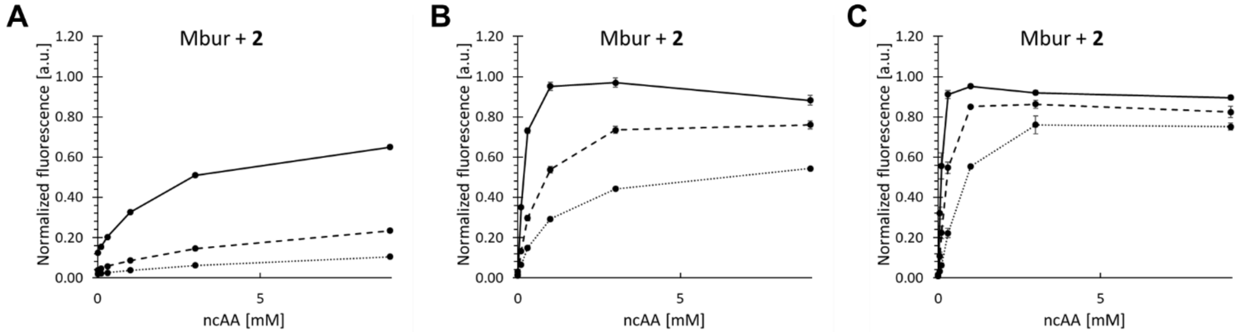

**Figure S7:** Concentration-dependent unnatural protein production with MburPylRS and substrate **2**, using **A**) the pUltra OTS setup, **B**) the pUltra setup with a constitutive lpp promotor instead of $P_{tac}$, **C**) the pTECH OTS setup. The host for protein production was RF1 deficient *Escherichia coli* B-95.ΔA. Fluorescence values were normalized their corresponding wild-type sfGFP reporter constructs (without in-frame stop codon), error bars represent the standard deviation of three replicates.

#### 1.7 Multi-Site ncAA incorporation of +N and ΔN PylRS

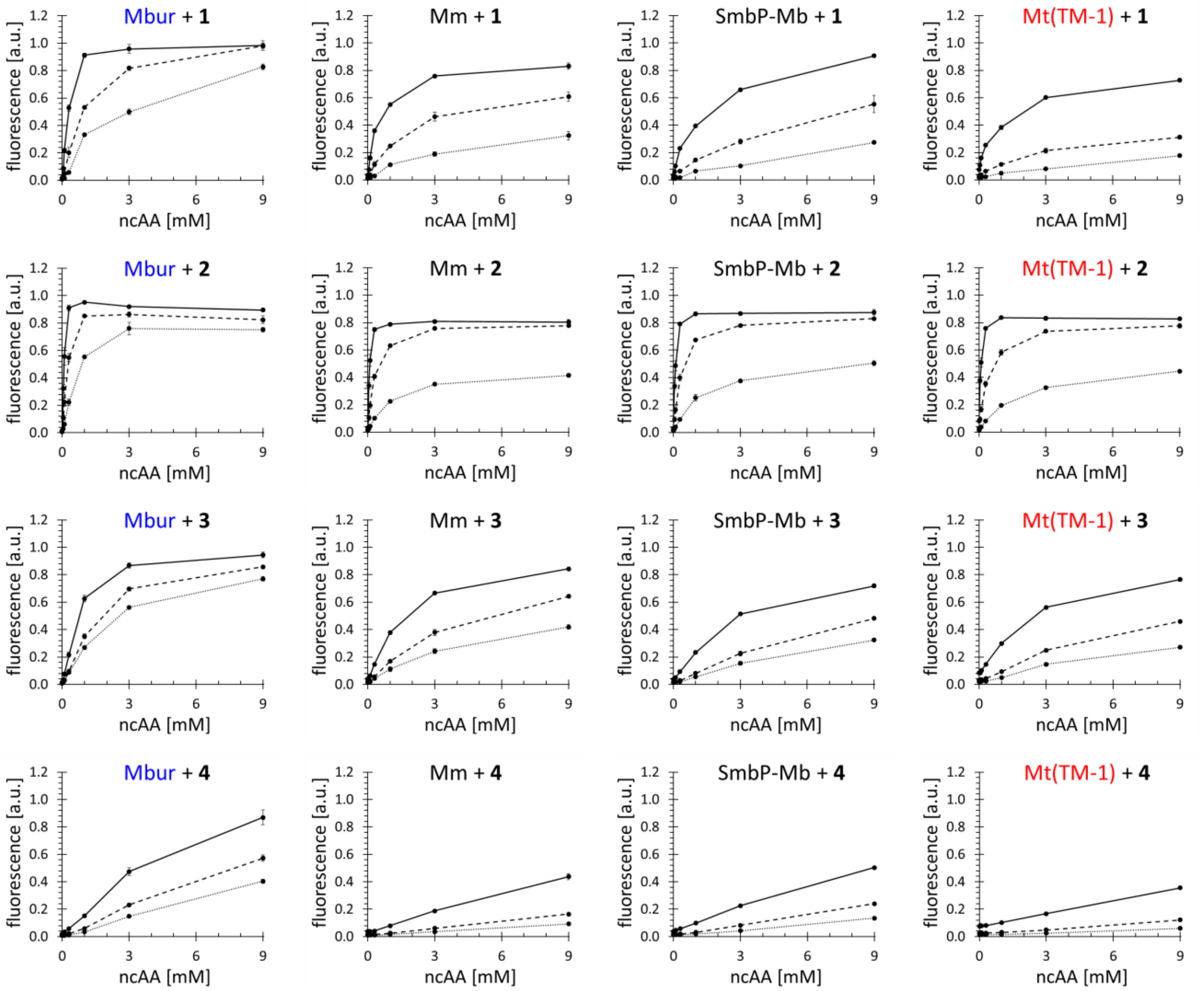

**Figure S8:** Concentration-dependent unnatural protein production using the four best +N PylRS from **Figure 2**. The host for protein production was RF1 deficient *Escherichia coli* B-95.ΔA. Endpoint measurements for ncAA concentrations of 0.05, 0.1, 0.3, 1, 3 and 9 mM. Fluorescence values were normalized their corresponding wild-type sfGFP reporter constructs (without in-frame stop codon), error bars represent the standard deviation of three replicates. Substrates and PylRS constructs are indicated in the headline.

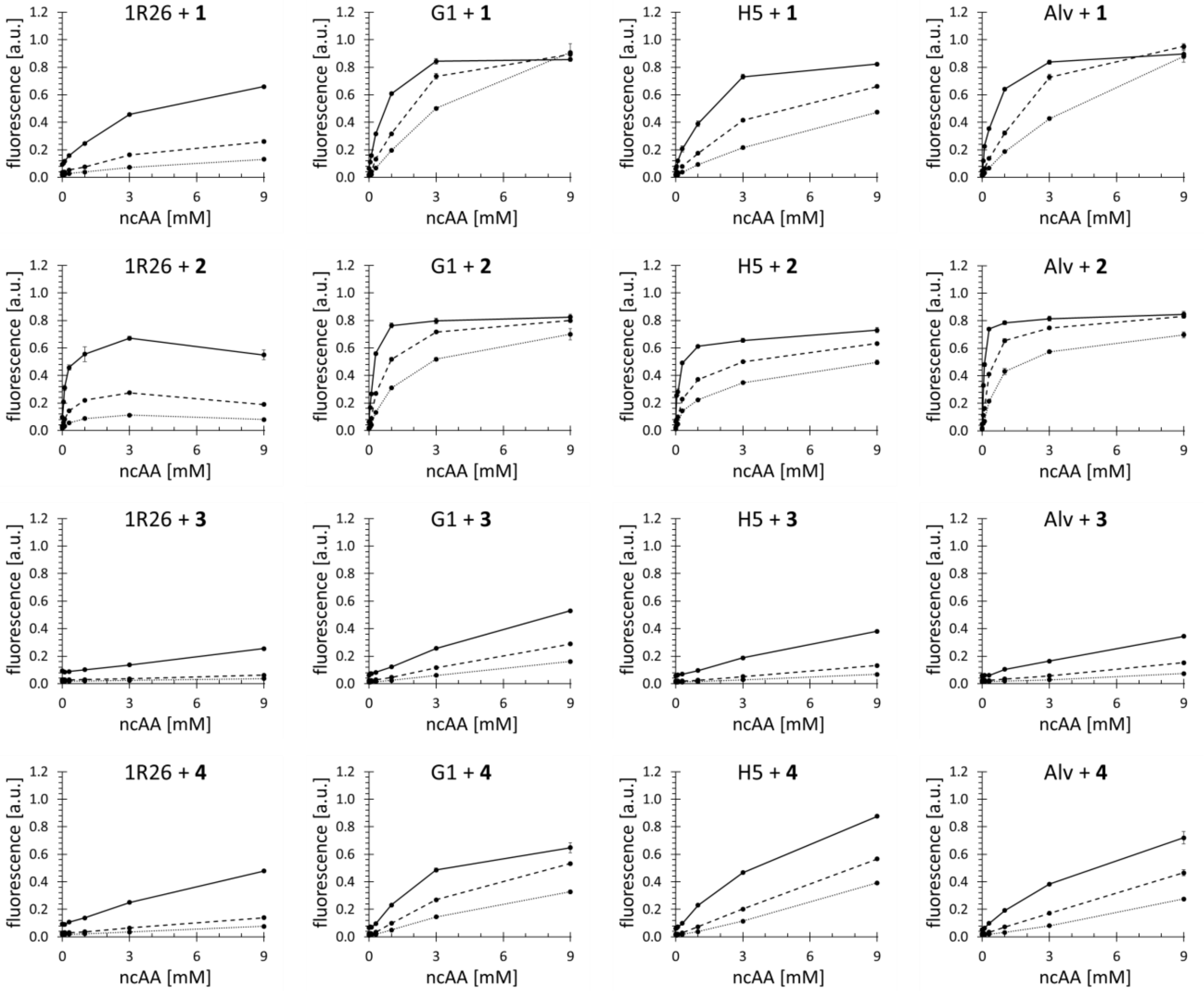

**Figure S9:** Concentration-dependent unnatural protein production using the four best ∆N PylRS from **Figure 2**. The host for protein production was RF1 deficient *Escherichia coli* B-95.ΔA. Endpoint measurements for ncAA concentrations of 0.05, 0.1, 0.3, 1, 3 and 9 mM. Fluorescence values were normalized their corresponding wild-type sfGFP reporter constructs (without in-frame stop codon), error bars represent the standard deviation of three replicates. Substrates and PylRS constructs are indicated in the headline.

#### 1.8 **Wild-type sfGFP and no ncAA protein production at 18°C and 37°C**

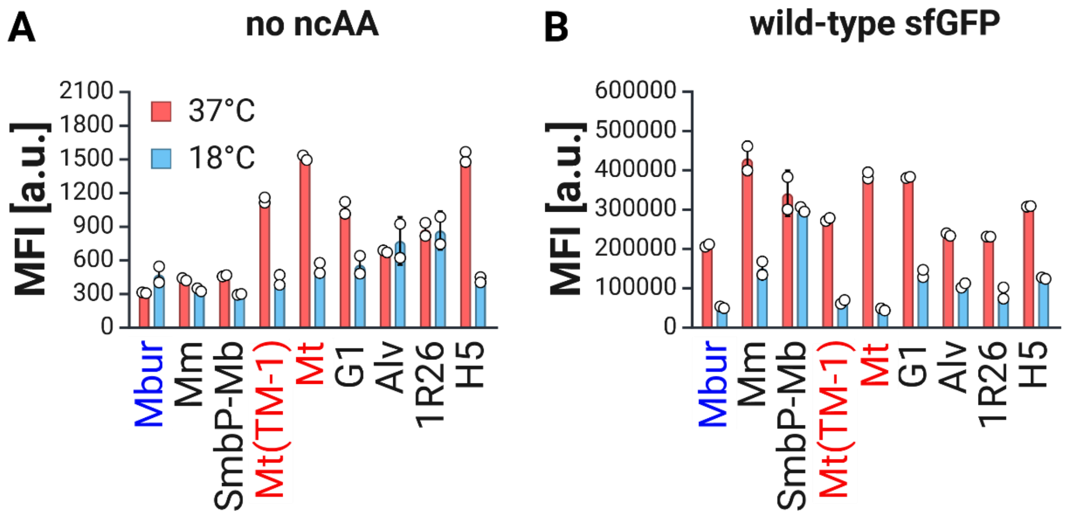

**Figure S10:** Temperature dependent protein production of **A**) sfGFP(5x amber) without supplied ncAA and **B**) wild-type sfGFP in the B-95.ΔA strain at 37°C(red columns) and 18°C(blue columns). The constructs were measured by flow cytometry in biological duplicates (n=2) and median fluorescence (MFI) of the target population is depicted.

#### 1.9 **Full Heatmaps for double Ala and Gly PylRS constructs**

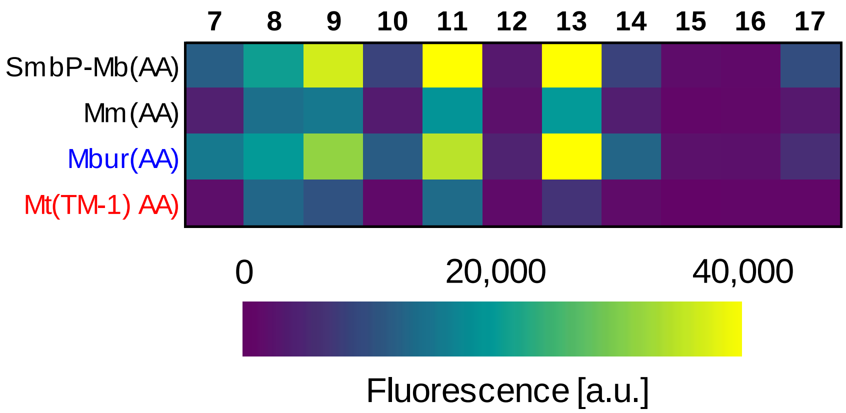

**Figure S11:** Heatmaps based on fluorescence of the sfGFP(1x amber) expression with *E. coli* BL21(DE3) and selected PylRS mutants with 10 mM supplied ncAA. Substrate numbering can be found in **Figure 1**. Background suppressions were construct dependent and ranged from 1,100 to 2,500 [a.u.].

#### 1.10 Additional data for promiscuity investigation

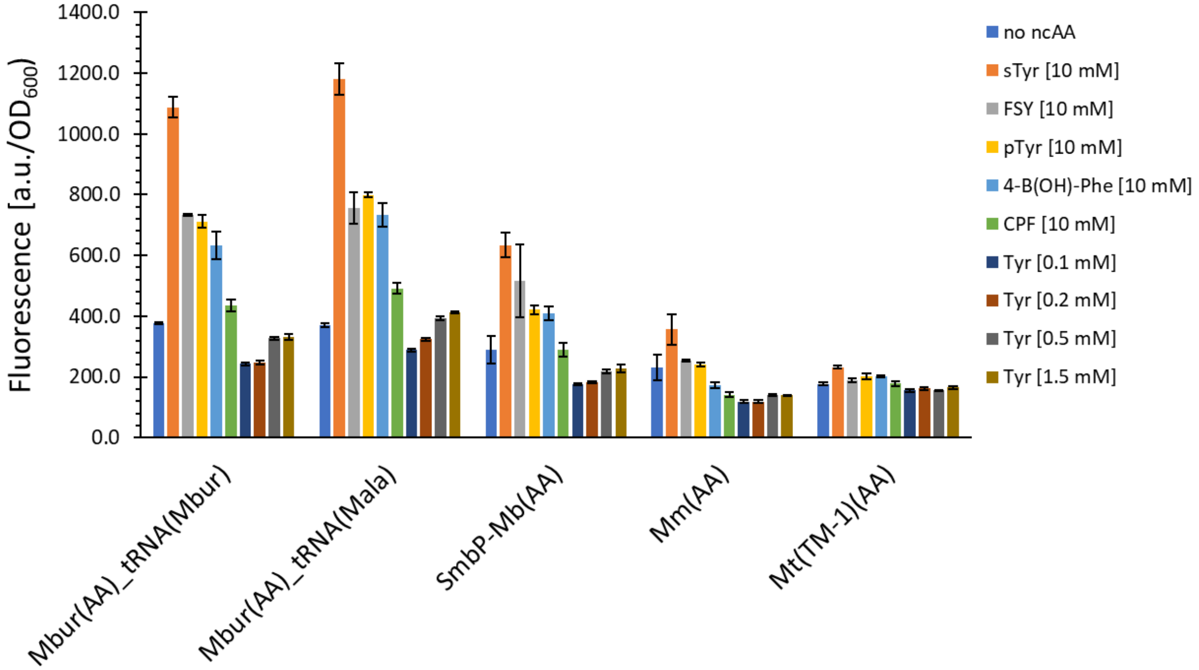

**Figure S12:** Comparison of charged and polar ncAA incorporation for selected PylRS variants. The fluorescence was measured for intact *E. coli* BL21(DE3) cells producing the sfGFP(1x amber) reporter. The data (incl. standard deviation) represent the mean of three biological replicates (n=3). ncAA abbreviations can be found in section 3.1, Table S4.

#### 1.11 Testing PylRS constructs for SproC (40) and Sac (43) incorporation

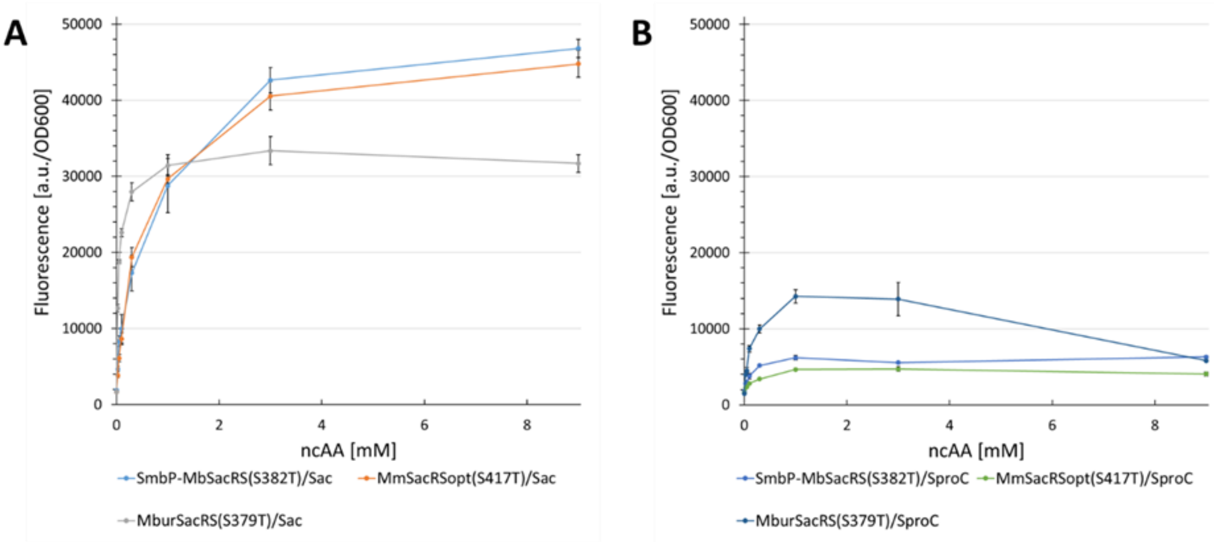

**Figure S13.** Concentration-dependent unnatural protein production for SacRS mutants supplemented with **A**) Sac (**38**). **B**) SproC (**35**). Using BL21(DE3) cells and sfGFP(1x amber) as reporter. Endpoint measurements for ncAA concentrations of 0.025, 0.05, 0.1, 0.3, 1, 3 and 9 mM.

#### 1.12 Multi-Site ncAAs Incorporation in BL21(DE3)

To compare the efficiency of Mbur with the best known PylRS OTS Mm) two substrates (**2** and **4**) were tested in conjunction with sfGFP reporter constructs containing between one and five stop-codons.

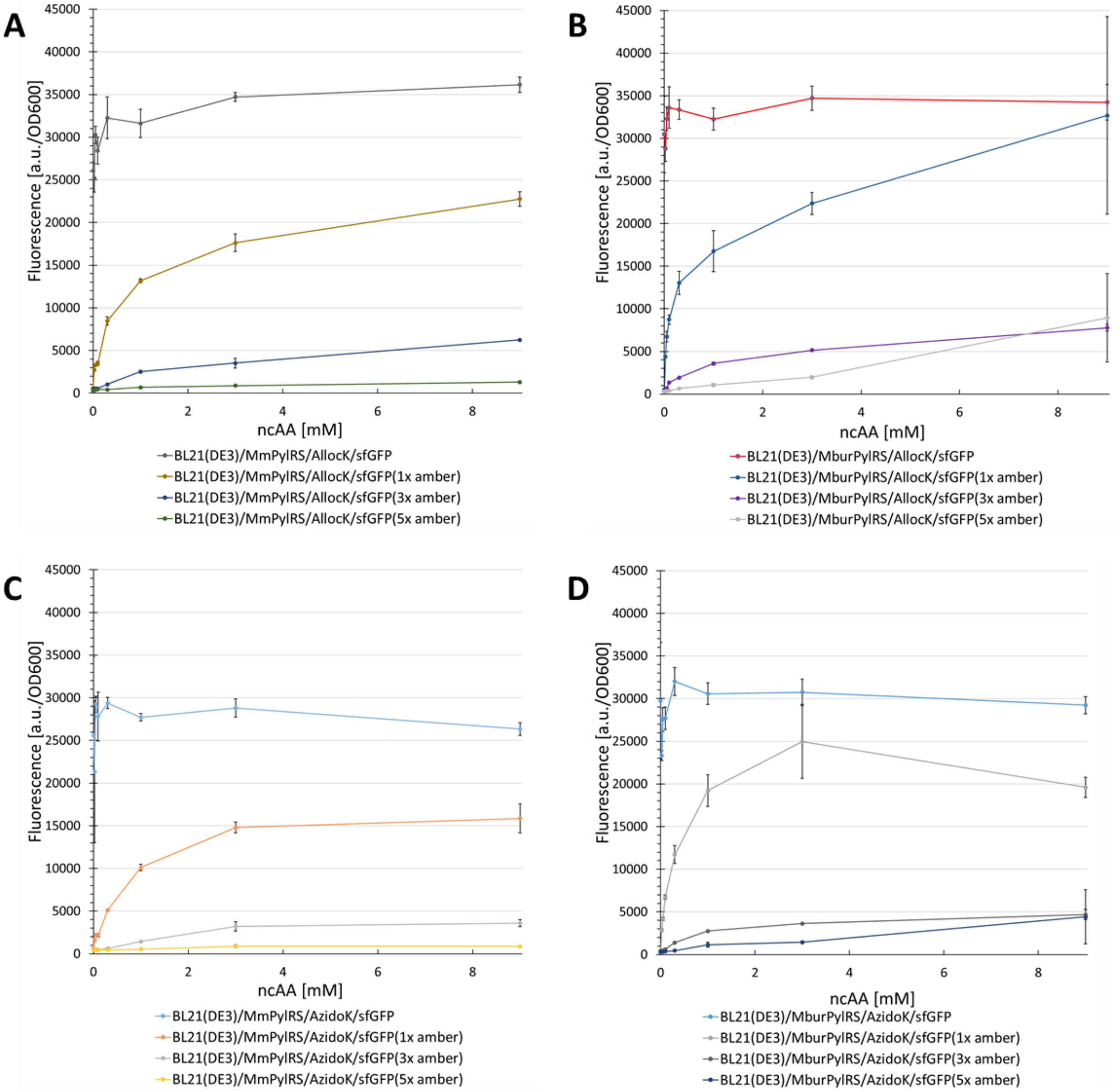

**Figure S14.** Concentration-dependent unnatural protein production with *Mm*PylRS (**A**, **C**) and *Mbur*PylRS (**B**, **D**) for different ncAA/reporter construct combinations.The host for protein production was BL21(DE3). Endpoint measurements for ncAA concentrations of 0.025, 0.05, 0.1, 0.3, 1, 3 and 9 mM.**Figure S8** shows that for BL21(DE3) expression of sfGFP(1x amber), the performances of both PylRS are comparable when supplied with AllocK(**2**). In the same setup with AzidoK(**4**), the *Mbur*PylRS outperforms *Mm*PylRS, and is twice as efficient. With the suppression of more than one stop-codon in the BL21(DE3) strain, the performance decreases significantly, but the *Mbur*PylRS shows at least twice the efficiency than the *Mm*PylRS in the range between 1 and 3 mM ncAA, albeit at a low level. For protein production in the B95.ΔA strain, the suppression of one stop-codon is similar to the BL21(DE3) experiment, but with generally higher suppression efficiencies (**Figure S9**). Encouragingly, it is possible with *Mbur*PylRS to achieve wild-type level protein production for both ncAAs when fed 1 mM. A comparison of the OTS performances with *Mbur*PylRS/AllocK (**2**) shows that the decrease in suppression efficiency of one to five stop-codons, when supplied with 1 mM, is 49%. With *Mm*PylRS, the decrease is 77%.

The performance does not decrease as much when more than one stop codon is suppressed compared to BL21(DE3), proving the advantage of employing *Mbur*PylRS. At an AzidoK(**4**) concentration of 1 mM and suppression of three and five stop-codons, the performance is three times better than that of *Mm*PylRS. This result suggests that *Mbur*PylRS is not only very efficient at low ncAA concentrations, but also more suitable for incorporation of ncAAs at multiple sites. This advantage might be even more pronounced in the absence of RF-1 competition, for example, in organisms with liberated codons or with sense codon suppression.

#### 1.13 Multi-Site Incorporation of Sac (43) and SproC (40)

Based on the high efficiency of the SacRS variants with the S→T mutation[14], the *M. barkeri* and *M. burtonii* variants were investigated for the multi-site incorporation of ncAAs with two different *E. coli* strains. The *M. barkeri* variant instead of the *M. mazei* variant was chosen, because of slightly better performance with SproC (**35**) (and equal Sac (**38**) activity). For reporter protein production in BL21(DE3) with Sac (**38**), the relative performance of *Mbur*SacRS(S379T) compared to SmbP-*Mb*SacRS(S382T) increases with the number of in-frame stop-codons (**Figure S10**). With one stop codon, the performance is 230% higher (at 0.3 mM), with three stop codons it is 490% higher (also at 0.3 mM), and five stop codons result in no incorporation with SmbP-*Mb*SacRS(S382T).

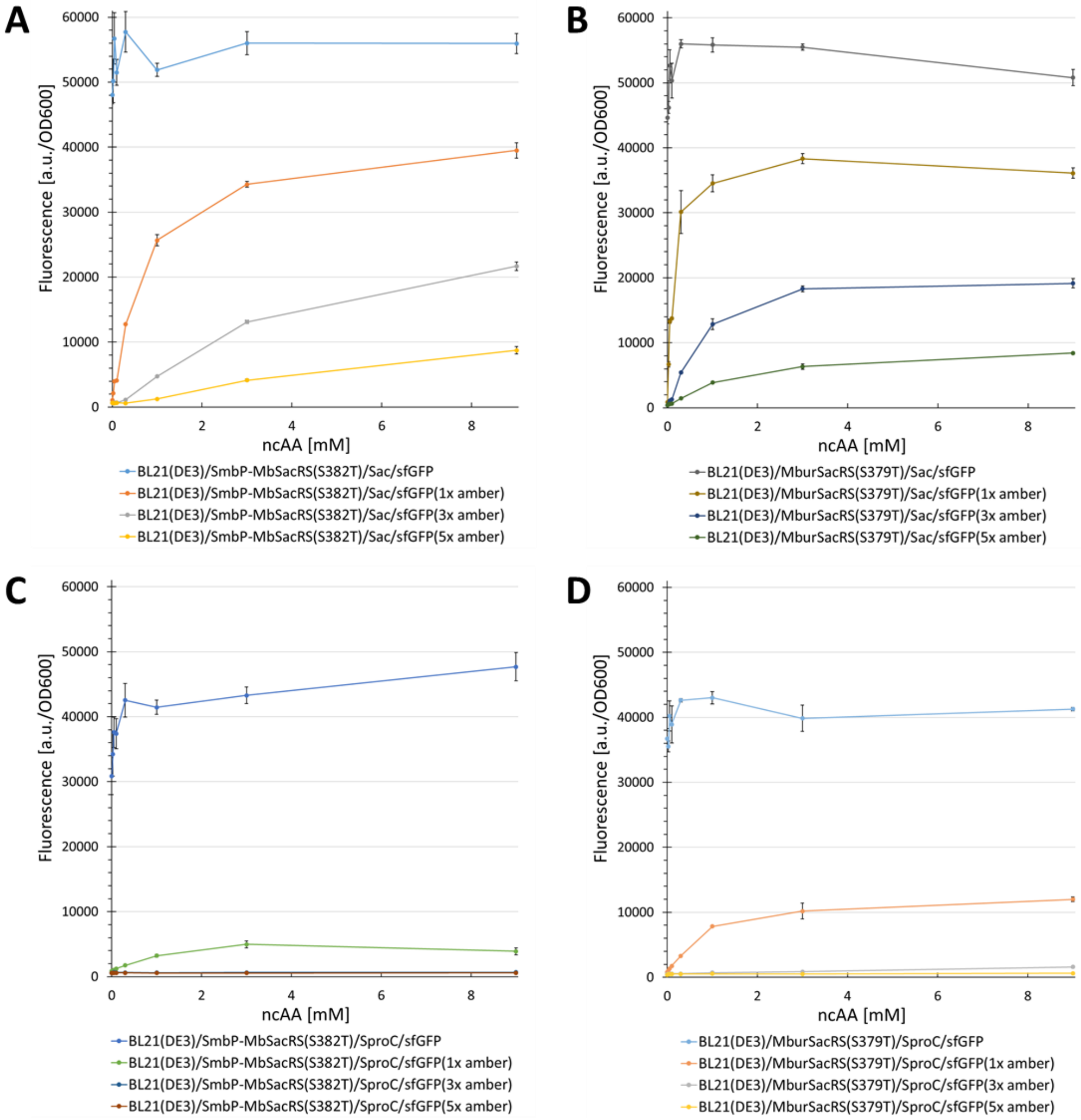

**Figure S15.** Concentration-dependent unnatural protein production with SmbP-*Mb*PylRS (**A**, **C**) and *Mbur*PylRS (**B**, **D**) for different ncAA/reporter construct combinations.The host for protein production was BL21(DE3). Endpoint measurements for ncAA concentrations of 0.025, 0.05, 0.1, 0.3, 1, 3 and 9 mM.The same trends are observed in the B-95.ΔA strain, with the difference that both OTS show significantly higher incorporation efficiencies when more than one in-frame stop codon is suppressed (**Figure S11**). But as with other *Mbur*PylRS variants, the efficiency is much higher at low ncAA concentrations and surprisingly, the decrease in incorporation efficiency with increasing number of in-frame stop codons is extremely low (for Sac (**1**)). A comparison of the OTSs performances with Sac (**1**) shows that the decrease for the *Mbur*SacRS(S379T) construct (fed with 1 mM) is 25% from the suppression of one to five in-frame stop codons. For SmbP-*Mb*SacRS(S382T) the decrease is

72%. As mentioned above, the SCS efficiency of *Mbur*SacRS(S379T) is at the same level as the wild-type enzyme when suppressing one stop codon, but when incorporating ncAAs at multi-sites, this mutant even surpasses the wild-type performance (compare **Figure 9** for the good substrate AllocK (**2**)). A PylRS mutant with better catalytic efficiency than the wild-type has never been reported before.

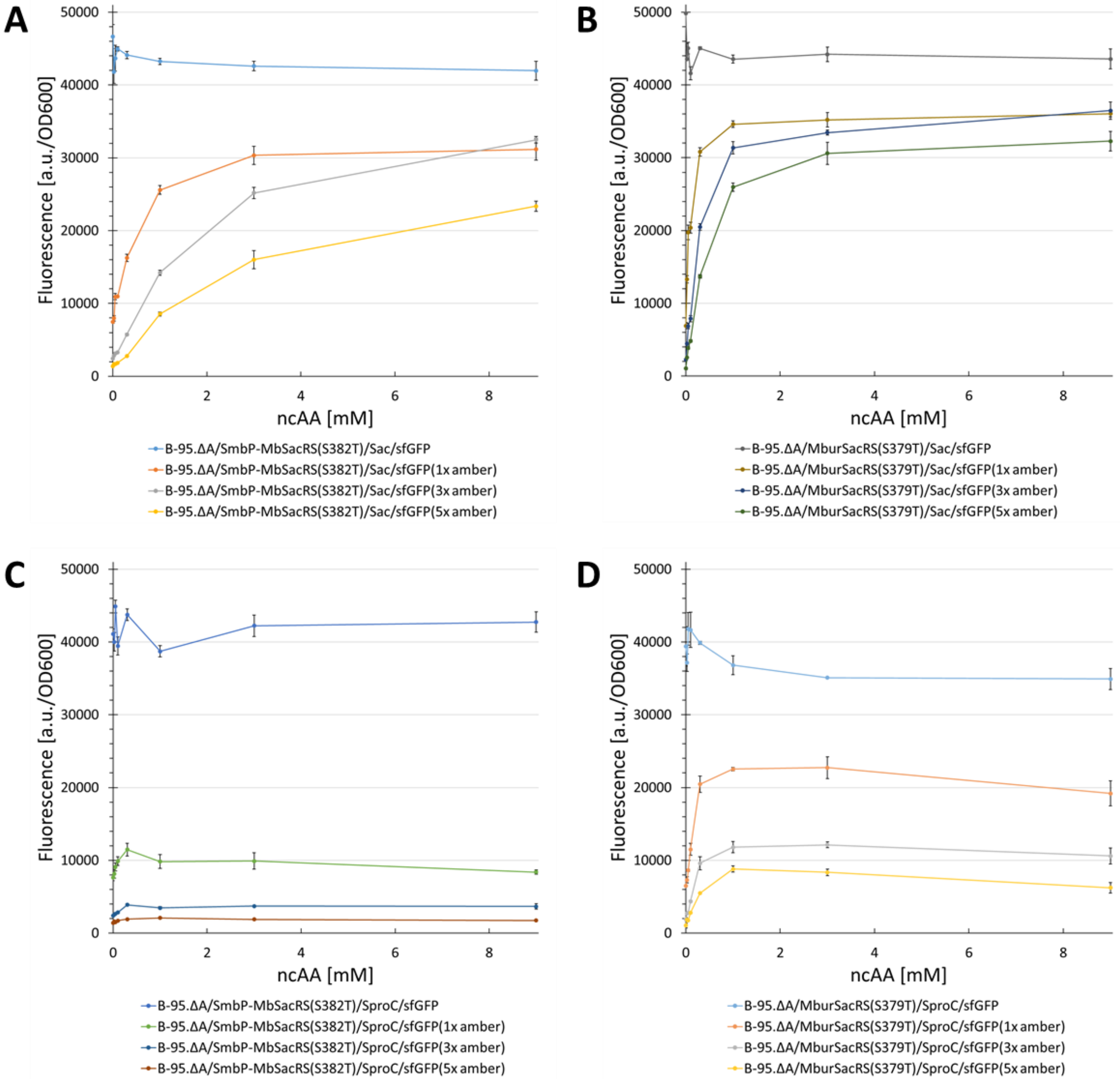

**Figure S16.** Concentration-dependent unnatural protein production with SmbP-*Mb*PylRS (**A**, **C**) and *Mbur*PylRS (**B**, **D**) for different ncAA/reporter construct combinations.The protein production host was B-95.ΔA. Endpoint measurements for ncAA concentrations of 0.025, 0.05, 0.1, 0.3, 1, 3 and 9 mM.

#### 1.14 Comparison of *Mj*TyrRS and *Mbur*PylRS performance

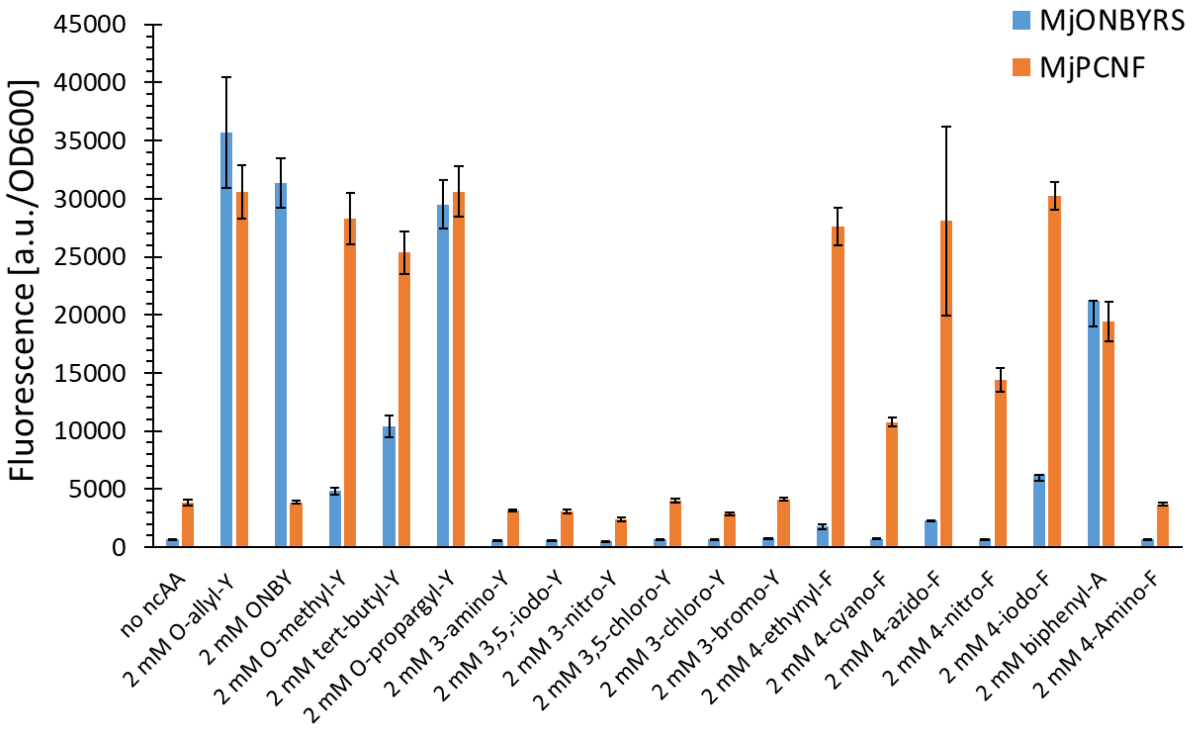

**Figure S17.** Prescreening of several ncAAs with MjONBYRS and MjPCNFRS. Fluorescence intensity of intact E. coli BL21(DE3) cells expressing the SUMO-sfGFP(1x amber) reporter. Endpoint measurements after 24 h with 2 mM ncAAs supplied. The data including the standard deviation represent the mean of three biological replicates.

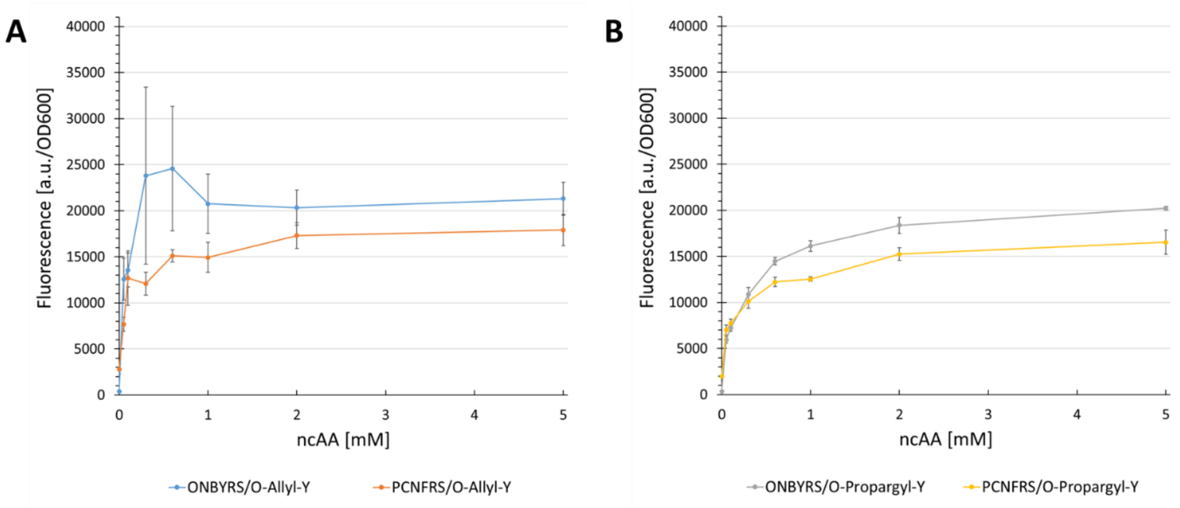

**Figure S18.** Concentration-dependent protein production for two different MjTyrRS/ncAA combinations. Fluorescence intensity of intact E. coli BL21(DE3) cells expressing the SUMO-sfGFP(1x amber) reporter. Endpoint measurements after 24 h with different ncAA concentrations (0.025, 0.05, 0.1, 0.3, 1, 2, and 5 mM). The data including the standard deviation represent the mean of three biological replicates.

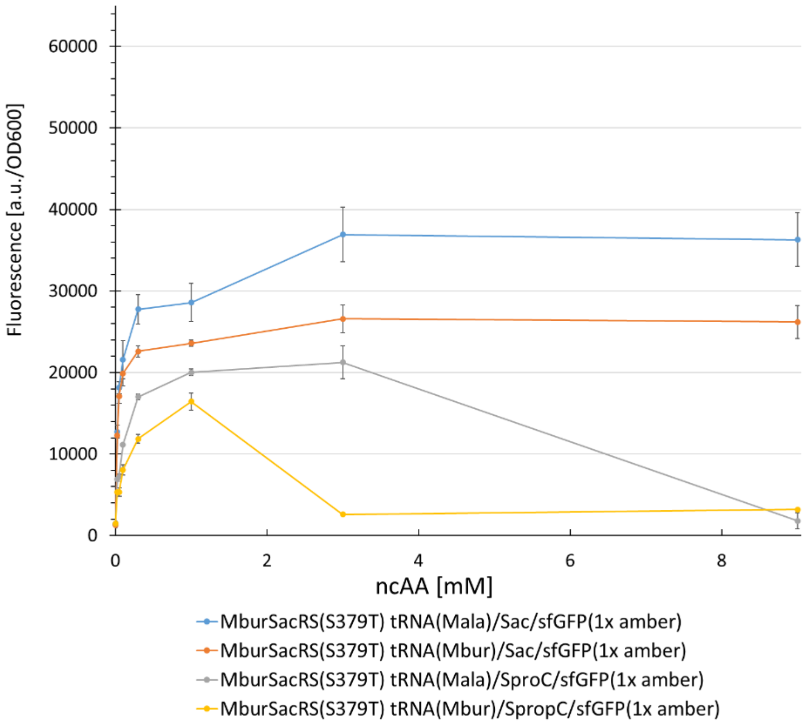

**Figure S19.** Concentration-dependent unnatural protein production with *Mbur*PylRS and different tRNA(organism abbreviation)/ncAA combinations. Using BL21(DE3) cells and sfGFP(1x amber) as reporter. Endpoint measurements for ncAA concentrations of 0.025, 0.05, 0.1, 0.3, 1, 3 and 9 mM.

### 2. Supplementary Data, DNA sequences and mass-profiles of intact ncAA-containing proteins

#### 2.1 Fluorescence Assays on the influence of ncAA concentration on PylRS-based OTS

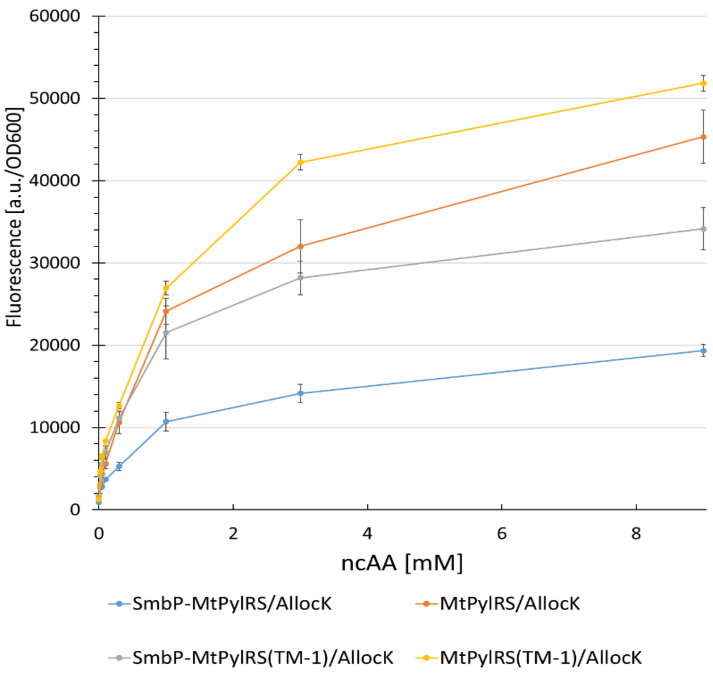

**Figure S20.** Concentration-dependent protein production for four different PylRS variants. Fluorescence intensity of intact *E. coli* BL21(DE3) cells expressing the SUMO-sfGFP(R2 amber) reporter, endpoint measurements after 24 h with different ncAA concentrations (0.025, 0.05, 0.1, 0.3, 1, 3, and 9 mM). Data including standard deviation represents the mean of three biological replicates.

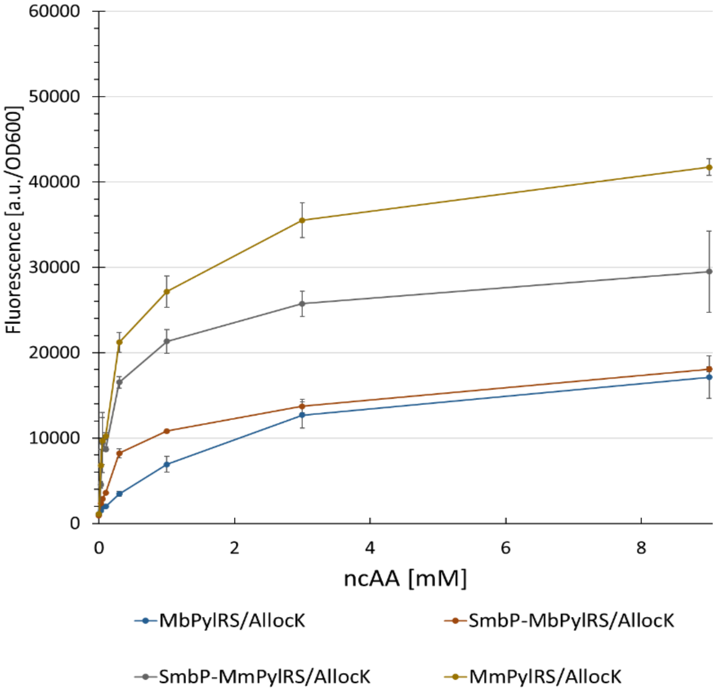

**Figure S21.** Concentration-dependent protein production for four different PylRS variants.Fluorescence intensity of intact *E. coli* BL21(DE3) cells expressing the SUMO-sfGFP(R2 amber) reporter, endpoint measurements after 24 h with different ncAA concentrations (0.025, 0.05, 0.1, 0.3, 1, 3, and 9 mM). Data including standard deviation represents the mean of three biological replicates.

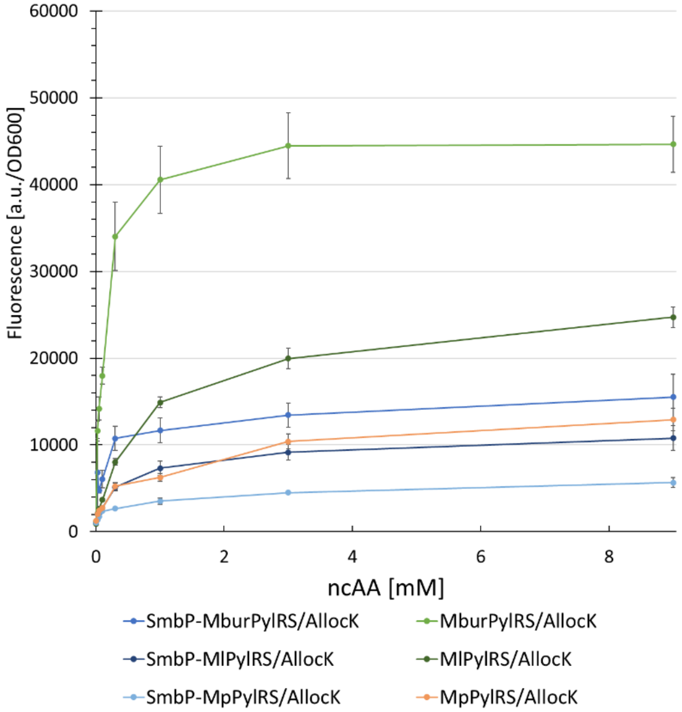

**Figure S22.** Concentration-dependent protein production for six different PylRS variants.Fluorescence intensity of intact *E. coli* BL21(DE3) cells expressing the SUMO-sfGFP(R2 amber) reporter, endpoint measurements after 24 h

with different ncAA concentrations (0.025, 0.05, 0.1, 0.3, 1, 3, and 9 mM). The data including the standard deviation

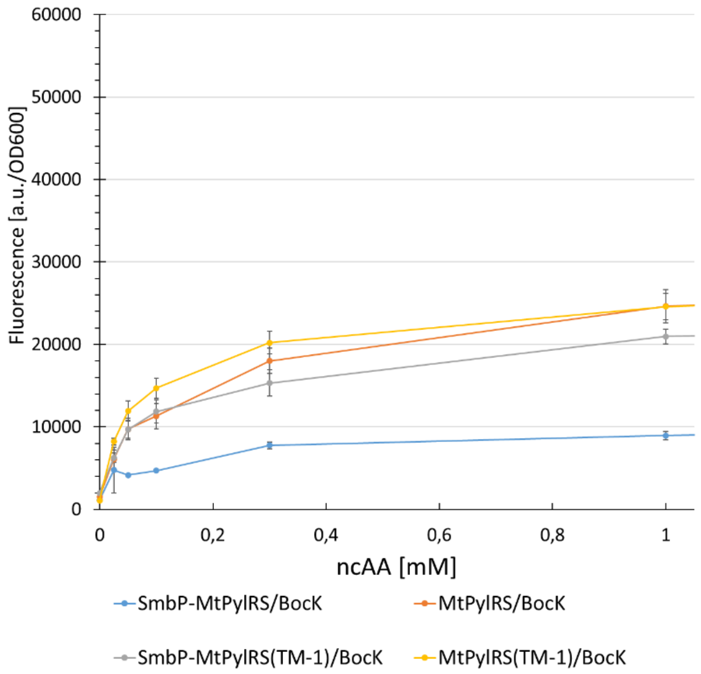

represent the mean of three biological replicates.

**Figure S23.** Concentration-dependent protein production for four different PylRS variants.Fluorescence intensity of intact *E. coli* BL21(DE3) cells expressing the SUMO-sfGFP(R2 amber) reporter, endpoint measurements after 24 h with different ncAA concentrations (0.025, 0.05, 0.1, 0.3, 1, 3, and 9 mM). Data including standard deviation represents

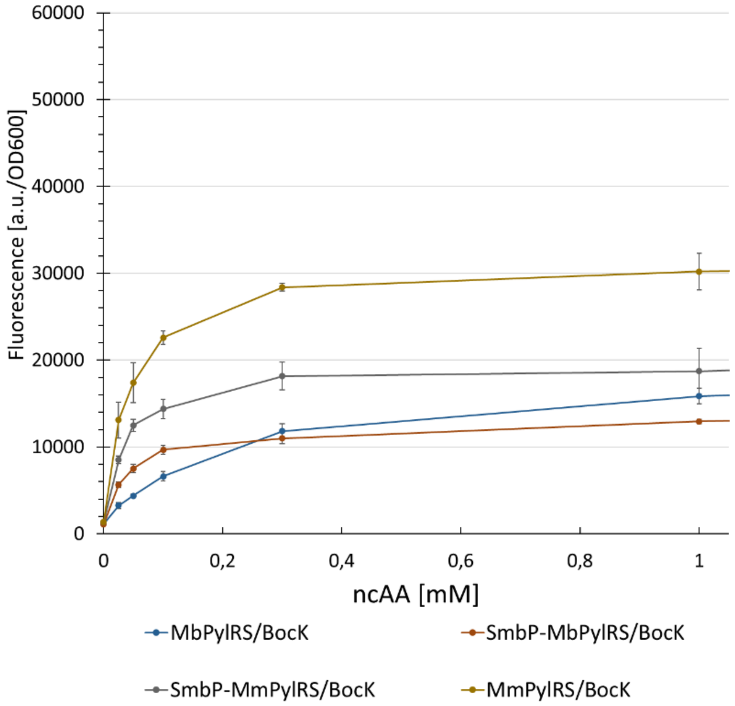

the mean of three biological replicates.

**Figure S24** Concentration-dependent protein production for four different PylRS variants.Fluorescence intensity of intact *E. coli* BL21(DE3) cells expressing the SUMO-sfGFP(R2 amber) reporter, endpoint measurements after 24 h with different ncAA concentrations (0.025, 0.05, 0.1, 0.3, 1, 3, and 9 mM). The data including the standard deviation represent the mean of three biological replicates.

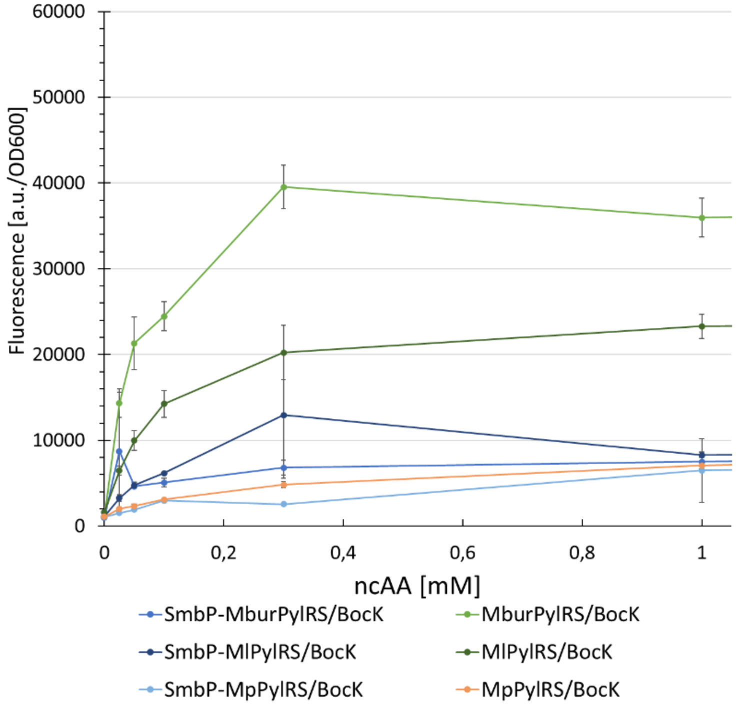

**Figure S25.** Concentration-dependent protein production for six different PylRS variants.Fluorescence intensity of intact *E. coli* BL21(DE3) cells expressing the SUMO-sfGFP(R2 amber) reporter, endpoint measurements after 24 h with different ncAA concentrations (0.025, 0.05, 0.1, 0.3, 1, 3, and 9 mM). Data including standard deviation represents the mean of three biological replicates.

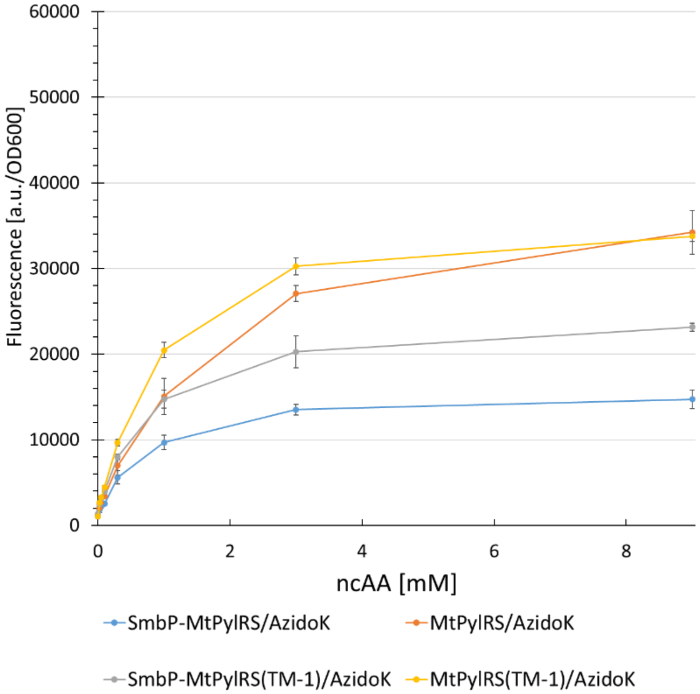

**Figure S26.** Concentration-dependent protein production for four different PylRS variants.Fluorescence intensity of intact *E. coli* BL21(DE3) cells expressing the SUMO-sfGFP(R2 amber) reporter, endpoint measurements after 24 h

with different ncAA concentrations (0.025, 0.05, 0.1, 0.3, 1, 3, and 9 mM). Data including standard deviation represents the mean of three biological replicates.

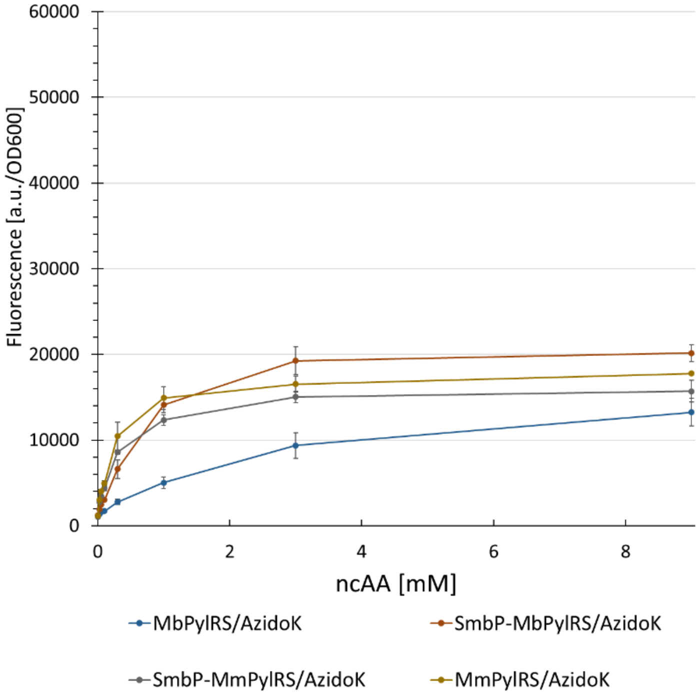

**Figure S27.** Concentration-dependent protein production for four different PylRS variants.Fluorescence intensity of intact *E. coli* BL21(DE3) cells expressing the SUMO-sfGFP(R2 amber) reporter, endpoint measurements after 24 h with different ncAA concentrations (0.025, 0.05, 0.1, 0.3, 1, 3, and 9 mM). Data including standard deviation represents the mean of three biological replicates.

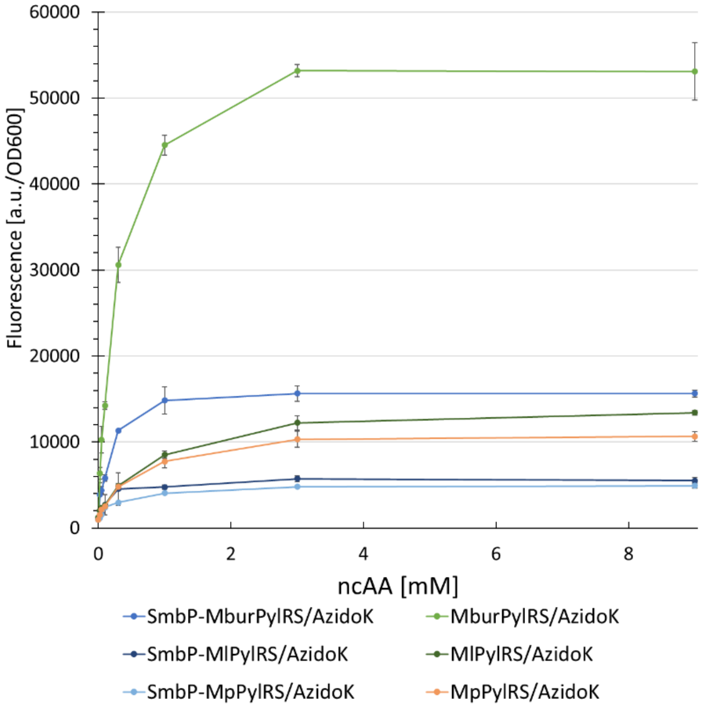

**Figure S28.** Concentration-dependent protein production for six different PylRS variants.Fluorescence intensity of intact *E. coli* BL21(DE3) cells expressing the SUMO-sfGFP(R2 amber) reporter, endpoint measurements after 24 h

with different ncAA concentrations (0.025, 0.05, 0.1, 0.3, 1, 3, and 9 mM). Data including standard deviation represents

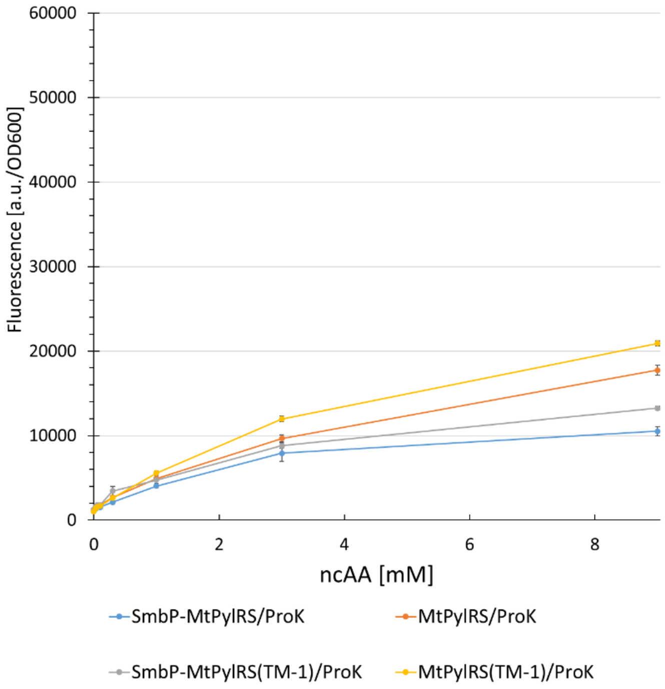

the mean of three biological replicates.

**Figure S29.** Concentration-dependent protein production for four different PylRS variants.Fluorescence intensity of intact *E. coli* BL21(DE3) cells expressing the SUMO-sfGFP(R2 amber) reporter, endpoint measurements after 24 h with different ncAA concentrations (0.025, 0.05, 0.1, 0.3, 1, 3, and 9 mM). Data including standard deviation represents

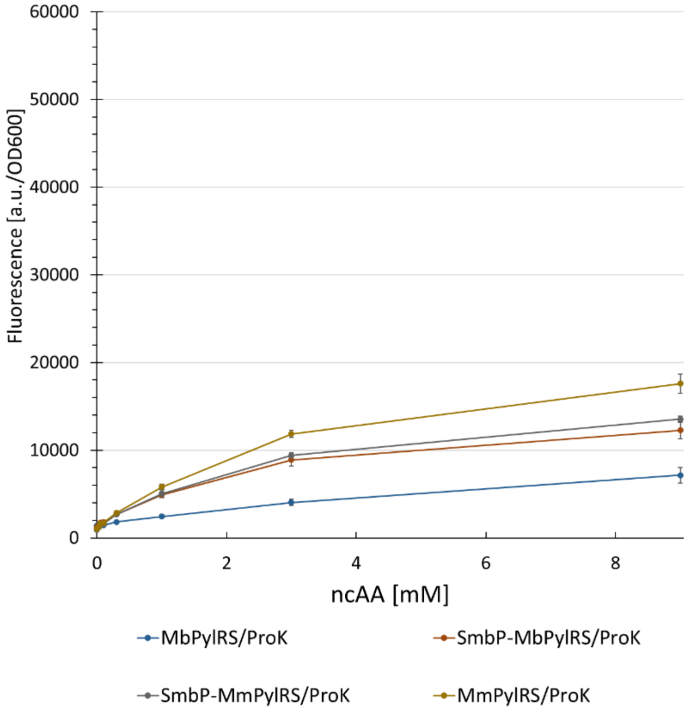

the mean of three biological replicates.

**Figure S30.** Concentration-dependent protein production for four different PylRS variants.Fluorescence intensity of intact *E. coli* BL21(DE3) cells expressing the SUMO-sfGFP(R2 amber) reporter, endpoint measurements after 24 h

with different ncAA concentrations (0.025, 0.05, 0.1, 0.3, 1, 3, and 9 mM). Data including standard deviation represents

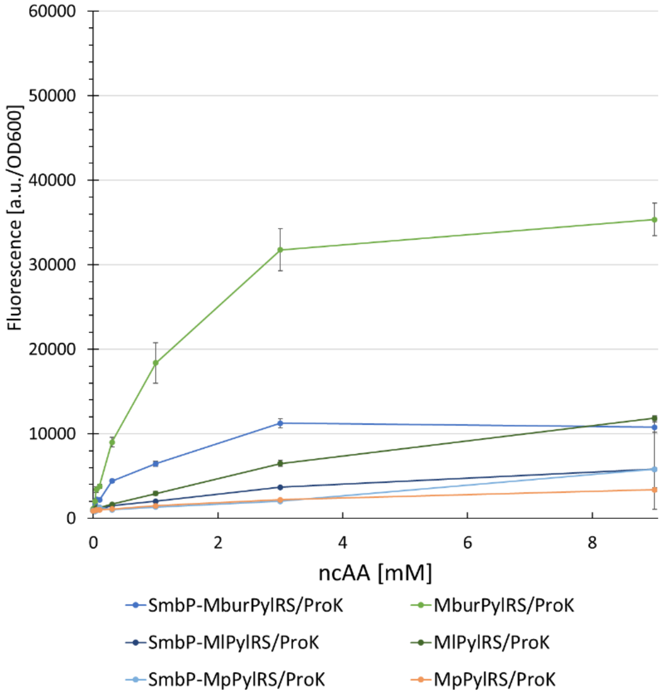

the mean of three biological replicates.

**Figure S31.** Concentration-dependent protein production for six different PylRS variants.Fluorescence intensity of intact *E. coli* BL21(DE3) cells expressing the SUMO-sfGFP(R2 amber) reporter, endpoint measurements after 24 h with different ncAA concentrations (0.025, 0.05, 0.1, 0.3, 1, 3, and 9 mM). Data including standard deviation represents

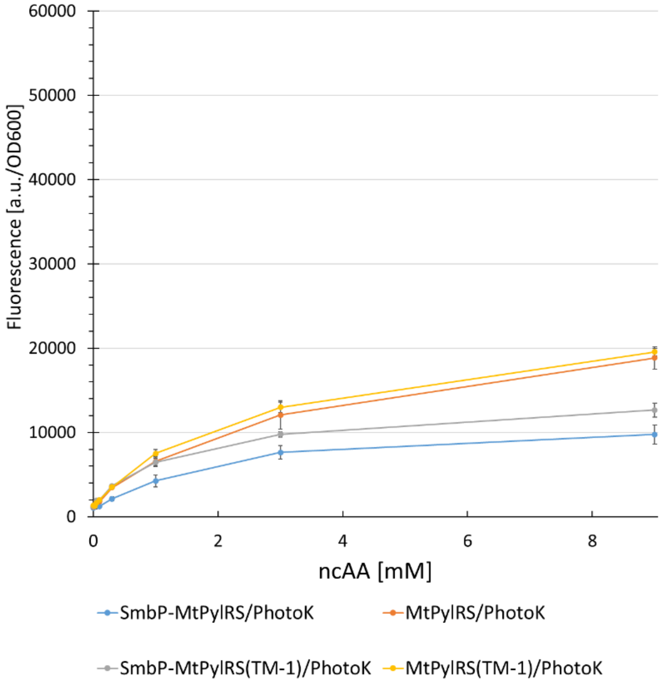

the mean of three biological replicates.

**Figure S32.** Concentration-dependent protein production for four different PylRS variants.Fluorescence intensity of intact *E. coli* BL21(DE3) cells expressing the SUMO-sfGFP(R2 amber) reporter, endpoint measurements after 24 h

with different ncAA concentrations (0.025, 0.05, 0.1, 0.3, 1, 3, and 9 mM). TData including standard deviation represents

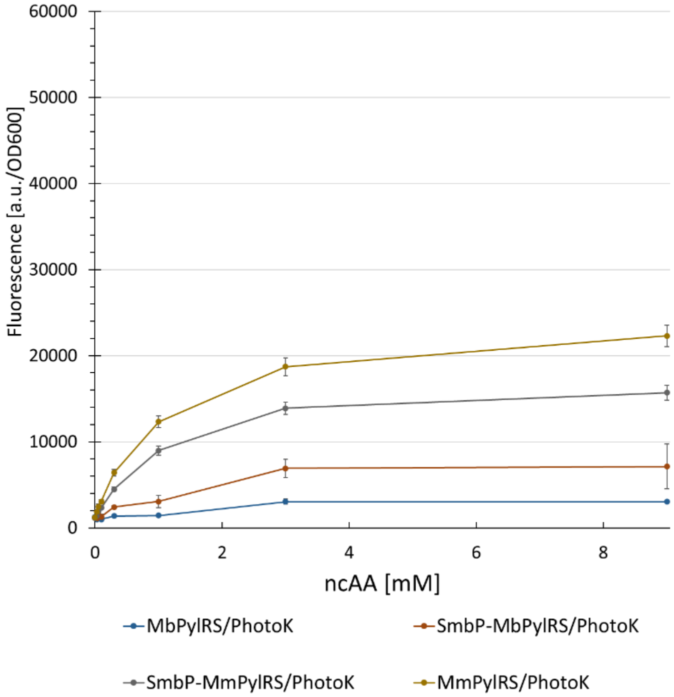

the mean of three biological replicates.

**Figure S33.** Concentration-dependent protein production for four different PylRS variants.Fluorescence intensity of intact *E. coli* BL21(DE3) cells expressing the SUMO-sfGFP(R2 amber) reporter, endpoint measurements after 24 h with different ncAA concentrations (0.025, 0.05, 0.1, 0.3, 1, 3, and 9 mM). Data including standard deviation represents

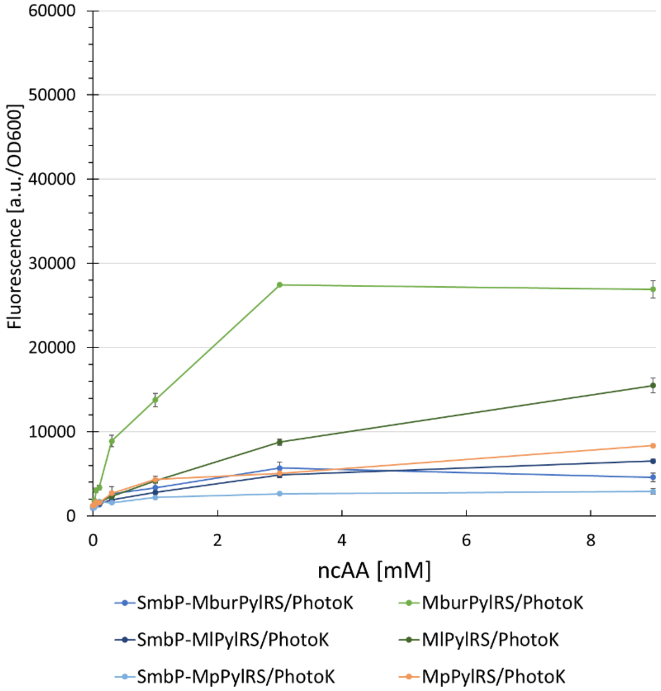

the mean of three biological replicates.

**Figure S34.** Concentration-dependent protein production for six different PylRS variants.Fluorescence intensity of intact *E. coli* BL21(DE3) cells expressing the SUMO-sfGFP(R2 amber) reporter, endpoint measurements after 24 h

with different ncAA concentrations (0.025, 0.05, 0.1, 0.3, 1, 3, and 9 mM). Data including standard deviation represents

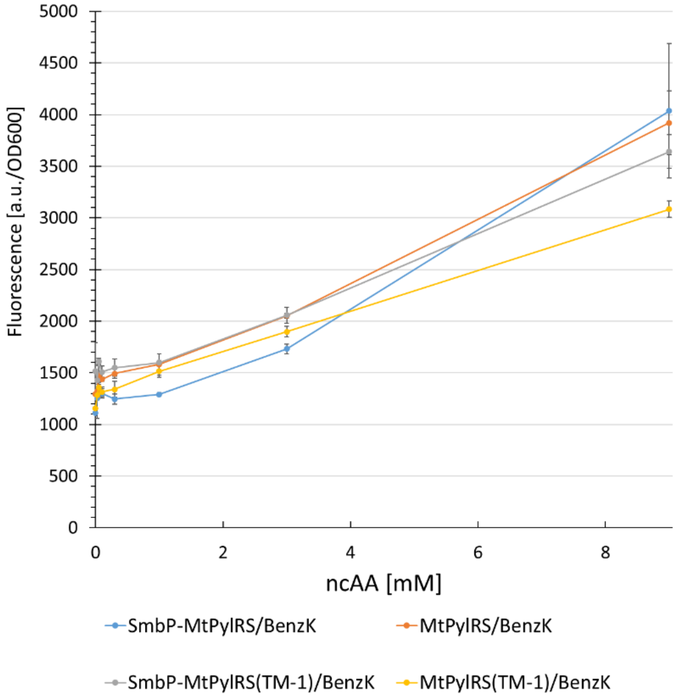

the mean of three biological replicates.

**Figure S35.** Concentration-dependent protein production for four different PylRS variants.Fluorescence intensity of intact *E. coli* BL21(DE3) cells expressing the SUMO-sfGFP(R2 amber) reporter, endpoint measurements after 24 h with different ncAA concentrations (0.025, 0.05, 0.1, 0.3, 1, 3, and 9 mM). Data including standard deviation represents

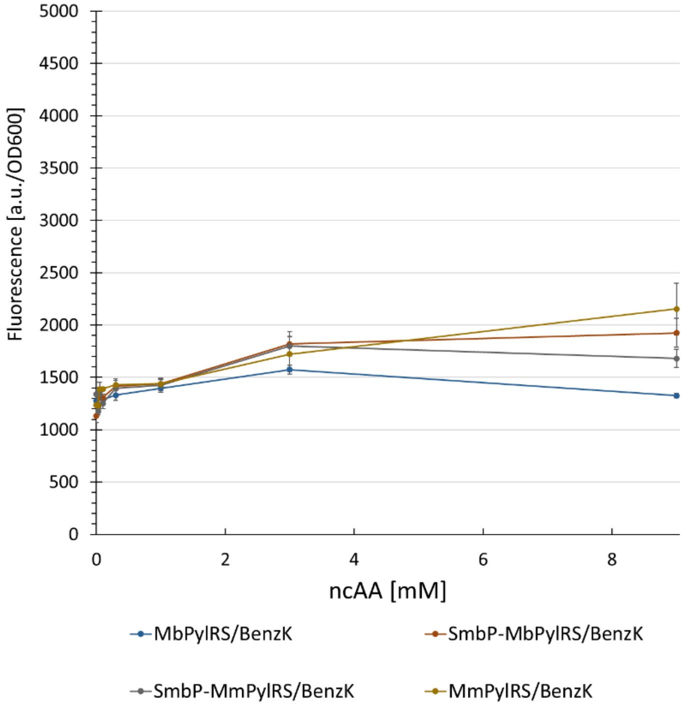

the mean of three biological replicates.

**Figure S36.** Concentration-dependent protein production for four different PylRS variants.Fluorescence intensity of intact *E. coli* BL21(DE3) cells expressing the SUMO-sfGFP(R2 amber) reporter, endpoint measurements after 24 h

with different ncAA concentrations (0.025, 0.05, 0.1, 0.3, 1, 3, and 9 mM). Data including standard deviation represents

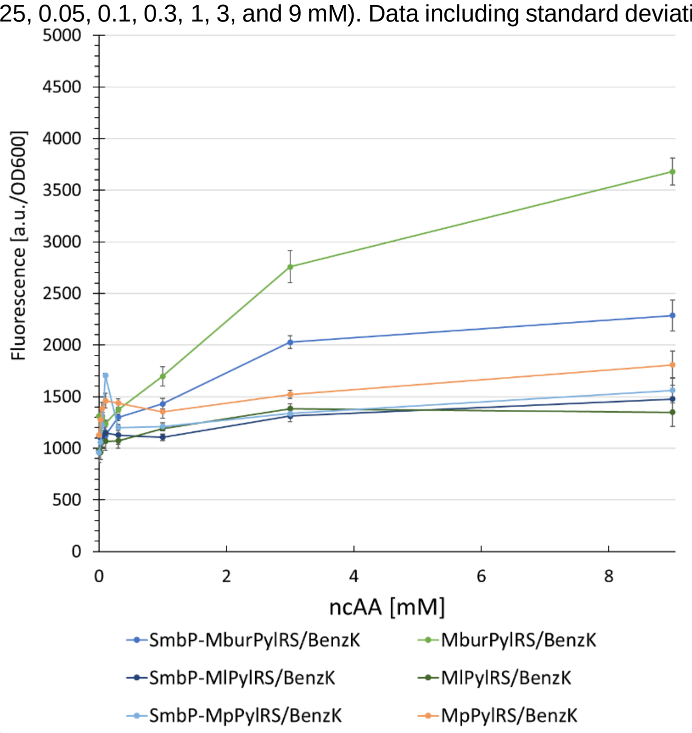

the mean of three biological replicates.

**Figure S37.** Concentration-dependent protein production for six different PylRS variants.Fluorescence intensity of intact *E. coli* BL21(DE3) cells expressing the SUMO-sfGFP(R2 amber) reporter, endpoint measurements after 24 h with different ncAA concentrations (0.025, 0.05, 0.1, 0.3, 1, 3, and 9 mM). Data including standard deviation represents the mean of three biological replicates.

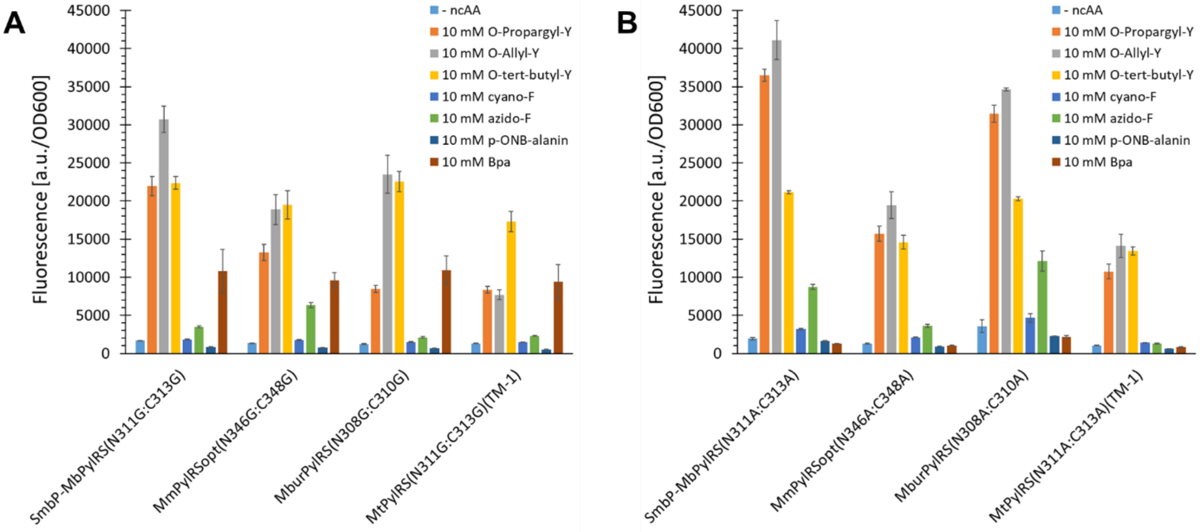

**Figure S38.** Comparison of ncAA (**39**, **40**, **44**, **45**, **46**, **49**, **50**) incorporation efficiency for PylRS double Ala and double Gly mutants.Fluorescence measurement of intact *E. coli* BL21(DE3) cells producing the SUMO-sfGFP(R2amber) reporter protein. Endpoint measurements after 24 h of incubation. Data including standard deviation represents the mean of three biological replicates. 10 mM ncAAs supplied

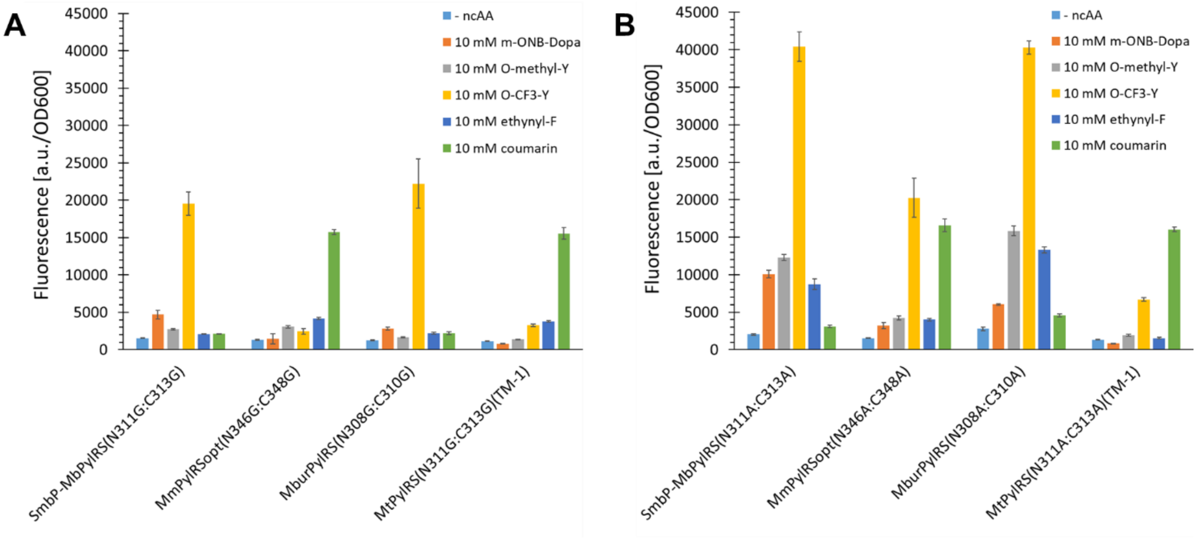

**Figure S39.** Comparison of ncAA (**38**, **47**, **48**, **51**, **52**) incorporation efficiency for PylRS double Ala and double Gly mutants.Fluorescence measurement of intact *E. coli* BL21(DE3) cells producing the SUMO-sfGFP(R2amber) reporter protein. Endpoint measurements after 24 h of incubation. Data including standard deviation represents the mean of three biological replicates. 10 mM ncAAs supplied

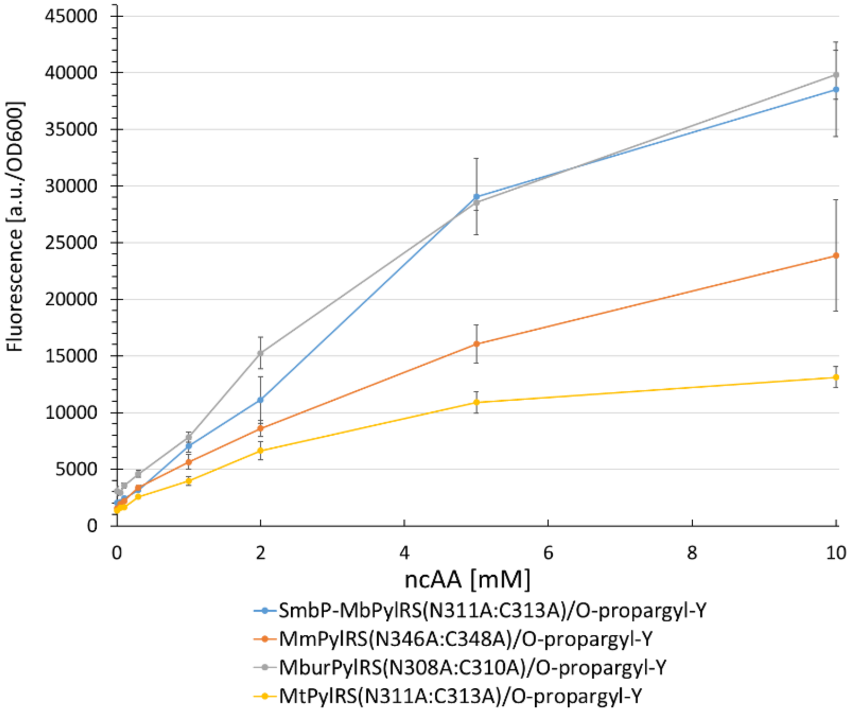

**Figure S40.** Concentration-dependent protein production for four different PylRS double Ala mutants.Fluorescence intensity of intact *E. coli* BL21(DE3) cells expressing the SUMO-sfGFP(R2 amber) reporter. Endpoint measurements after 24 h with different ncAA concentrations (0.5, 0.1, 0.3, 1, 2, 3, and 10 mM). Data including standard deviation

represents the mean of three biological replicates.

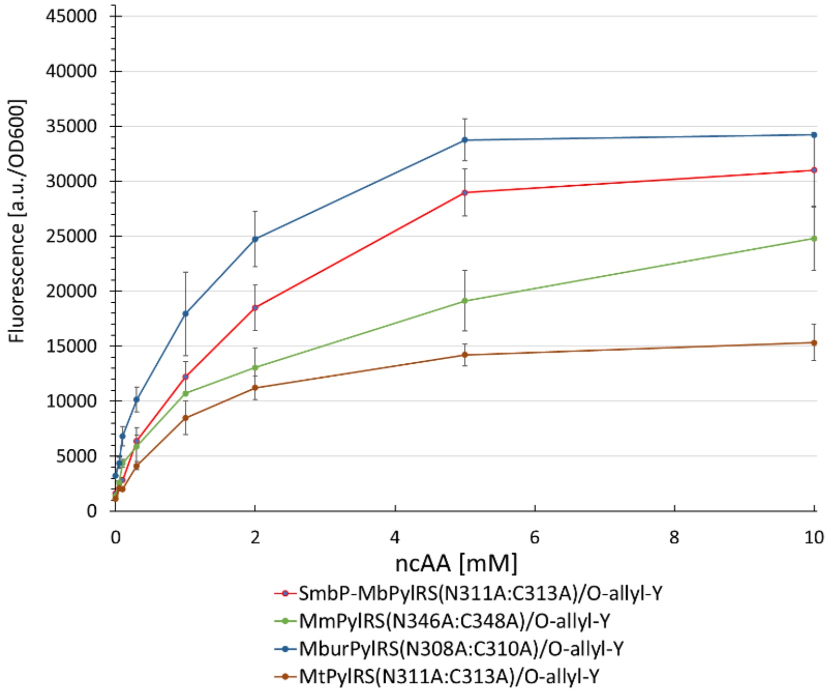

**Figure S41.** Concentration-dependent protein production for four different PylRS double Ala mutants. Fluorescence intensity of intact *E. coli* BL21(DE3) cells expressing the SUMO-sfGFP(R2 amber) reporter. Endpoint measurements after 24 h with different ncAA concentrations (0.5, 0.1, 0.3, 1, 2, 3, and 10 mM). Data including standard deviation represents the mean of three biological replicates.

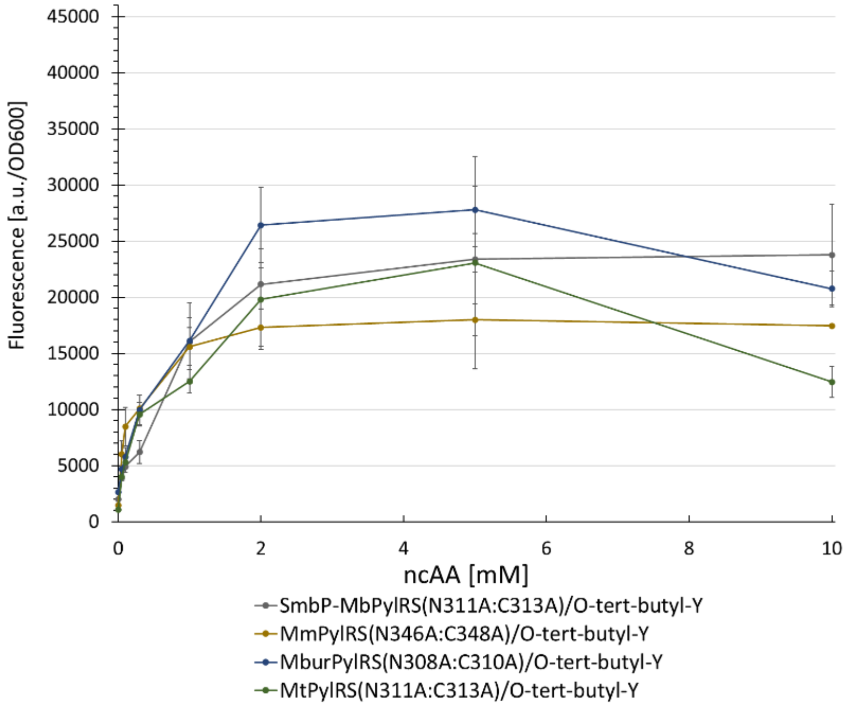

**Figure S42.** Concentration-dependent protein production for four different PylRS double Ala mutants. Fluorescence intensity of intact *E. coli* BL21(DE3) cells expressing the SUMO-sfGFP(R2 amber) reporter. Endpoint measurements after 24 h with different ncAA concentrations (0.5, 0.1, 0.3, 1, 2, 3, and 10 mM). Data including standard deviation

represents the mean of three biological replicates.

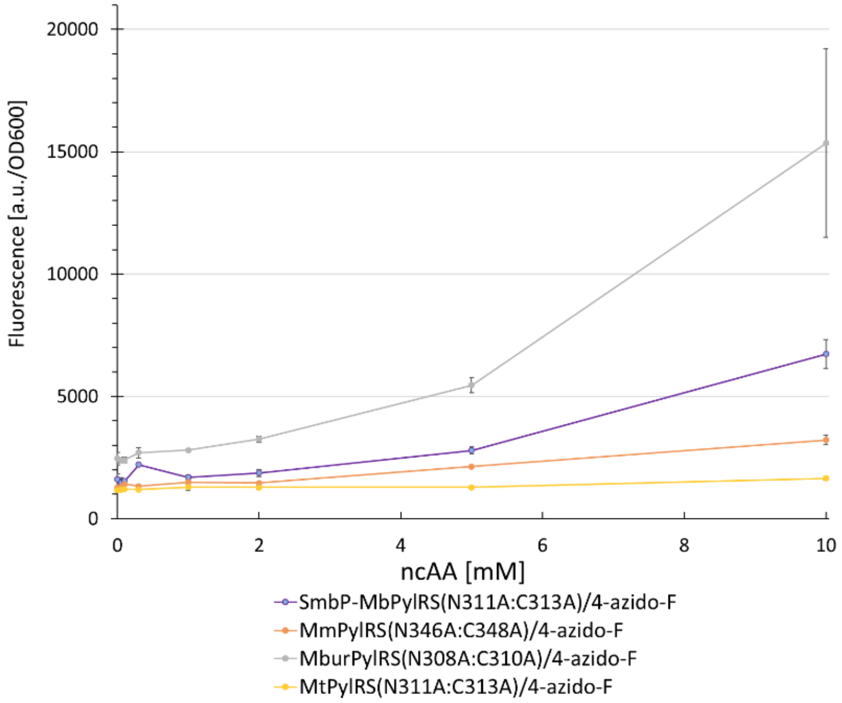

**Figure S43.** Concentration-dependent protein production for four different PylRS double Ala mutants. Fluorescence intensity of intact *E. coli* BL21(DE3) cells expressing the SUMO-sfGFP(R2 amber) reporter. Endpoint measurements after 24 h with different ncAA concentrations (0.5, 0.1, 0.3, 1, 2, 3, and 10 mM). Data including standard deviation represents the mean of three biological replicates.

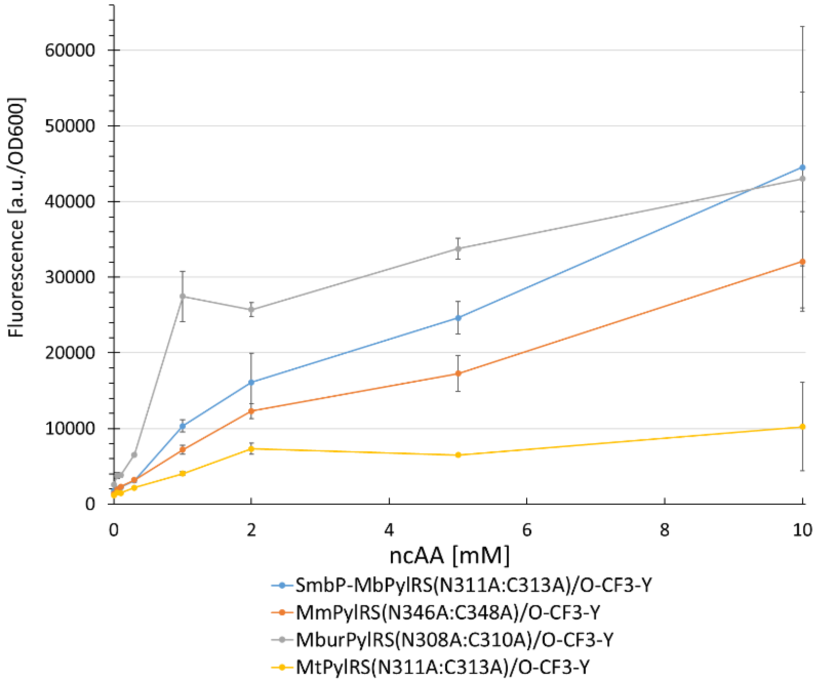

**Figure S44.** Concentration-dependent protein production for four different PylRS double Ala mutants. Fluorescence intensity of intact *E. coli* BL21(DE3) cells expressing the SUMO-sfGFP(R2 amber) reporter. Endpoint measurements after 24 h with different ncAA concentrations (0.5, 0.1, 0.3, 1, 2, 3, and 10 mM). Data including standard deviation

represents the mean of three biological replicates.

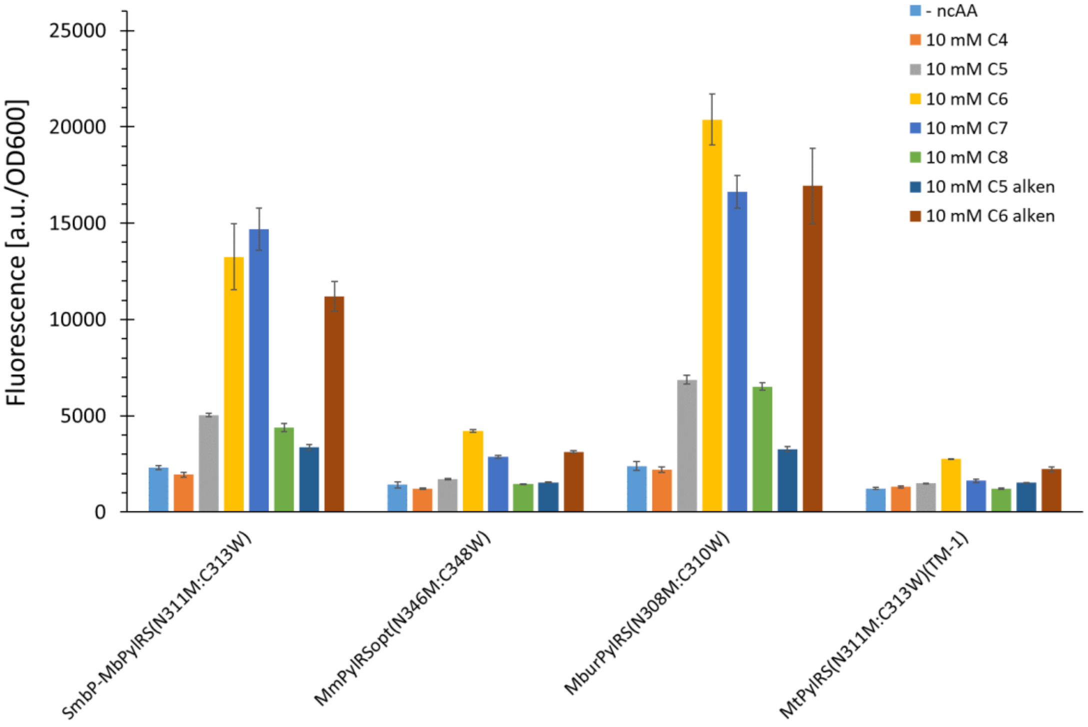

**Figure S45.** Comparison of ncAA (**10**, **7**, **5**, **2**, **3**, **8**, **6**) incorporation efficiency for different PylRS constructs. Fluorescence measurement of intact *E. coli* BL21(DE3) cells producing the SUMO-sfGFP(R2amber) reporter protein. Endpoint measurements after 24 h of incubation. Data including standard deviation represents the mean of three biological replicates. 10 mM ncAAs supplied

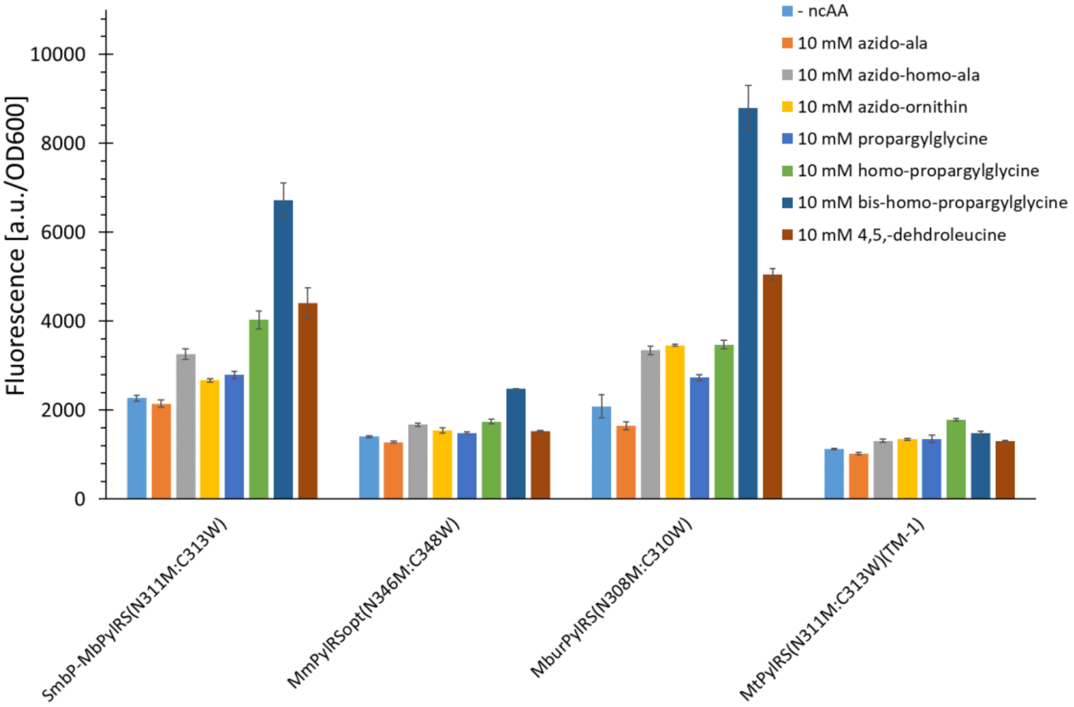

**Figure S46.** Comparison of ncAA (**14**, **15**, **16**, **13**, **12**, **11**, **21**) incorporation efficiency for different PylRS constructs. Fluorescence measurement of intact *E. coli* BL21(DE3) cells producing the SUMO-sfGFP(R2amber) reporter protein. Endpoint measurements after 24 h of incubation. Data including standard deviation represents the mean of three

biological replicates. 10 mM ncAAs supplied

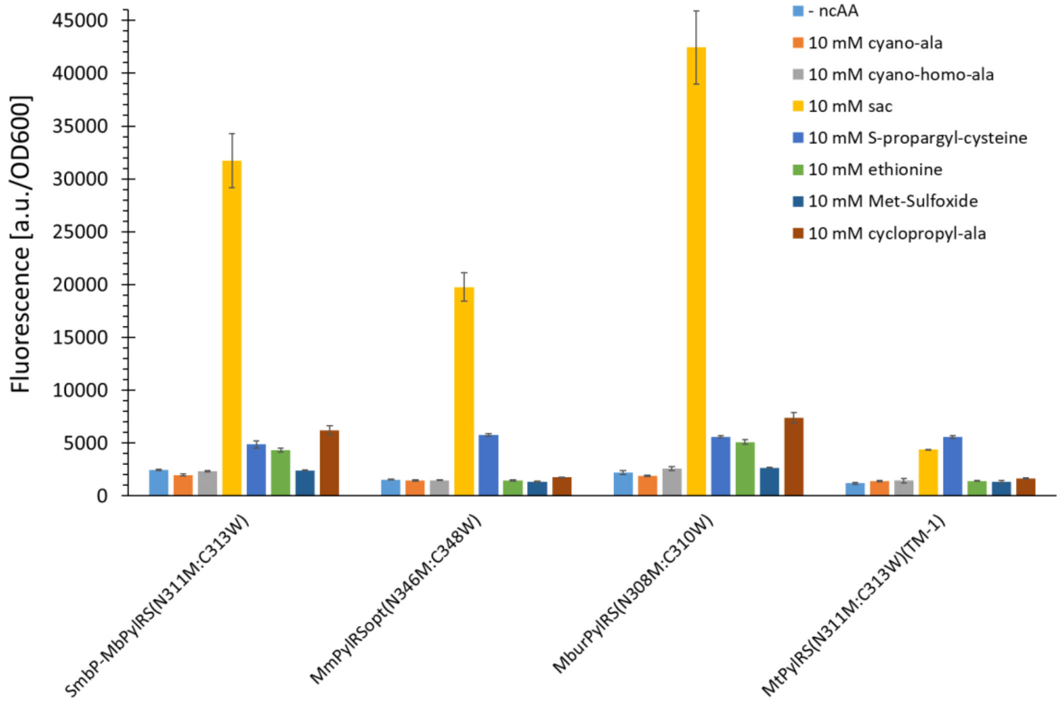

**Figure S47.** Comparison of ncAA (**18**, **19**, **1**, **27**, **25**, **23**, **9**) incorporation efficiency for different PylRS constructs. Fluorescence measurement of intact *E. coli* BL21(DE3) cells producing the SUMO-sfGFP(R2amber) reporter protein. Endpoint measurements after 24 h of incubation. Data including standard deviation represents the mean of three biological replicates. 10 mM ncAAs supplied

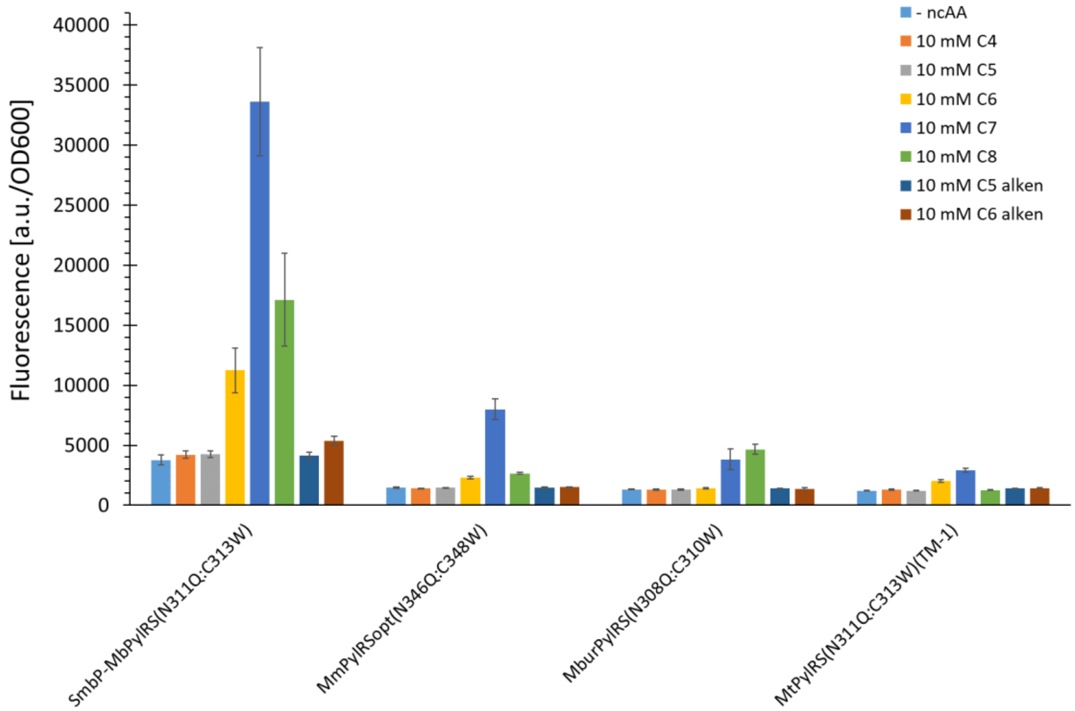

**Figure S48.** Comparison of ncAA (**10**, **7**, **5**, **2**, **3**, **8**, **6**) incorporation efficiency for different PylRS constructs. Fluorescence measurement of intact *E. coli* BL21(DE3) cells producing the SUMO-sfGFP(R2amber) reporter protein. Endpoint measurements after 24 h of incubation. Data including standard deviation represents the mean of three

biological replicates. 10 mM ncAAs supplied

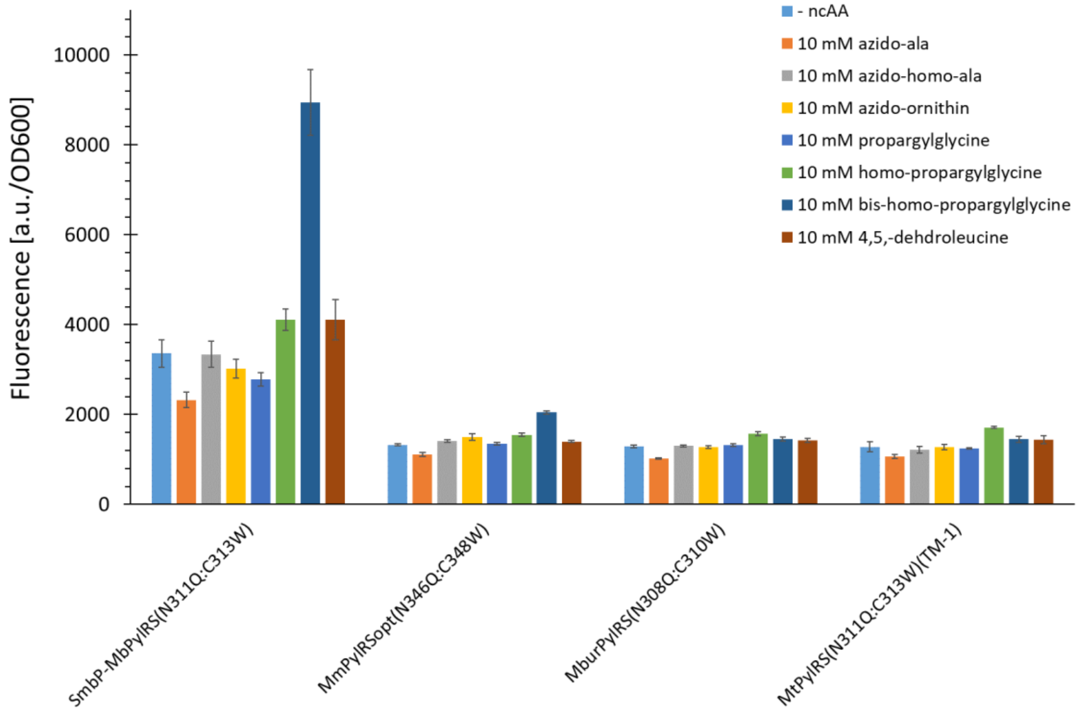

**Figure S49.** Comparison of ncAA (**14**, **15**, **16**, **13**, **12**, **11**, **21**) incorporation efficiency for different PylRS constructs. Fluorescence measurement of intact *E. coli* BL21(DE3) cells producing the SUMO-sfGFP(R2amber) reporter protein. Endpoint measurements after 24 h of incubation. Data including standard deviation represents the mean of three biological replicates. 10 mM ncAAs supplied

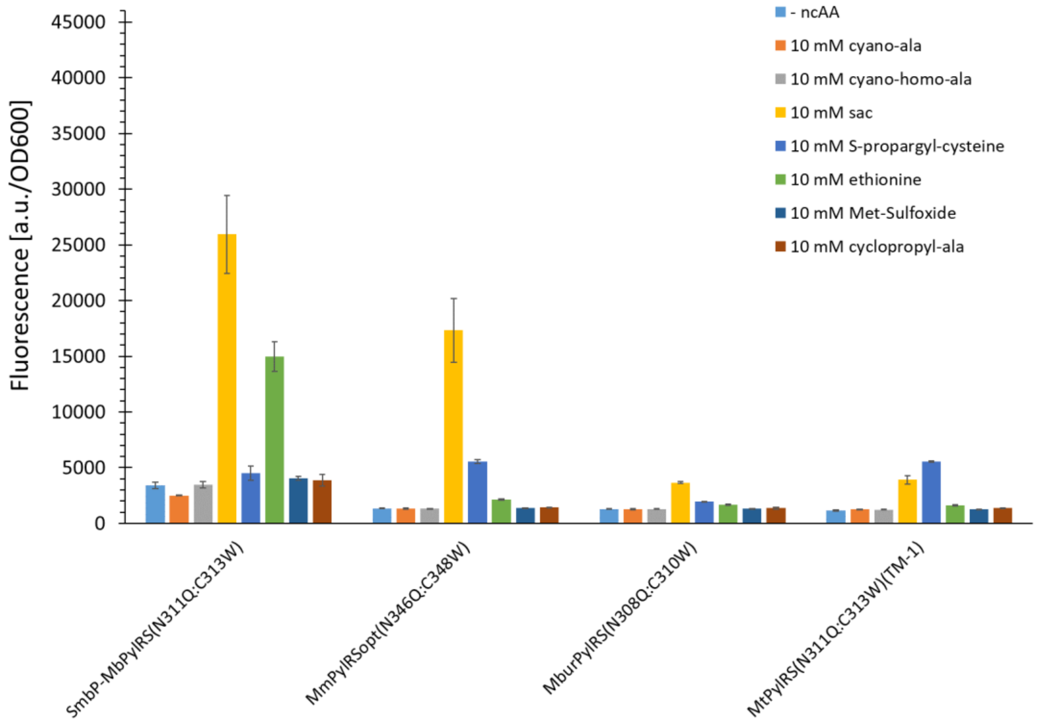

**Figure S50.** Comparison of ncAA (**18**, **19**, **1**, **27**, **25**, **23**, **9**) incorporation efficiency for different PylRS constructs. Fluorescence measurement of intact *E. coli* BL21(DE3) cells producing the SUMO-sfGFP(R2amber) reporter protein. Endpoint measurements after 24 h of incubation. Data including standard deviation represents the mean of three

biological replicates. 10 mM ncAAs supplied

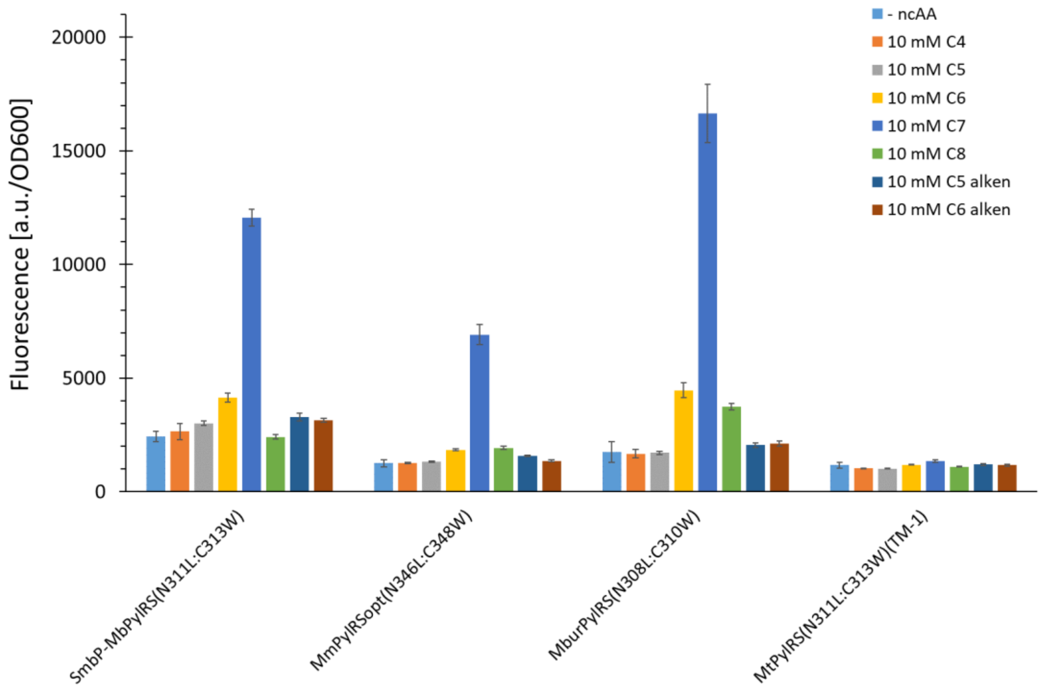

**Figure S51.** Comparison of ncAA (**10**, **7**, **5**, **2**, **3**, **8**, **6**) incorporation efficiency for different PylRS constructs. Fluorescence measurement of intact *E. coli* BL21(DE3) cells producing the SUMO-sfGFP(R2amber) reporter protein. Endpoint measurements after 24 h of incubation. Data including standard deviation represents the mean of three biological replicates. 10 mM ncAAs supplied

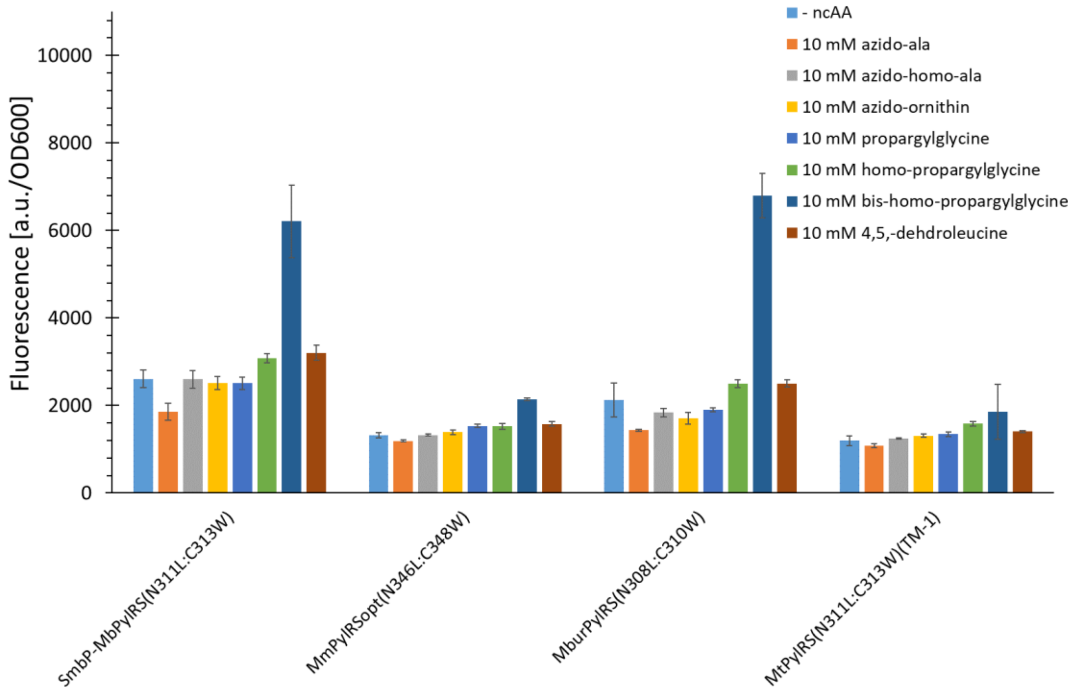

**Figure S52.** Comparison of ncAA (**14**, **15**, **16**, **13**, **12**, **11**, **21**) incorporation efficiency for different PylRS constructs. Fluorescence measurement of intact *E. coli* BL21(DE3) cells producing the SUMO-sfGFP(R2amber) reporter protein. Endpoint measurements after 24 h of incubation. Data including standard deviation represents the mean of three

biological replicates. 10 mM ncAAs supplied

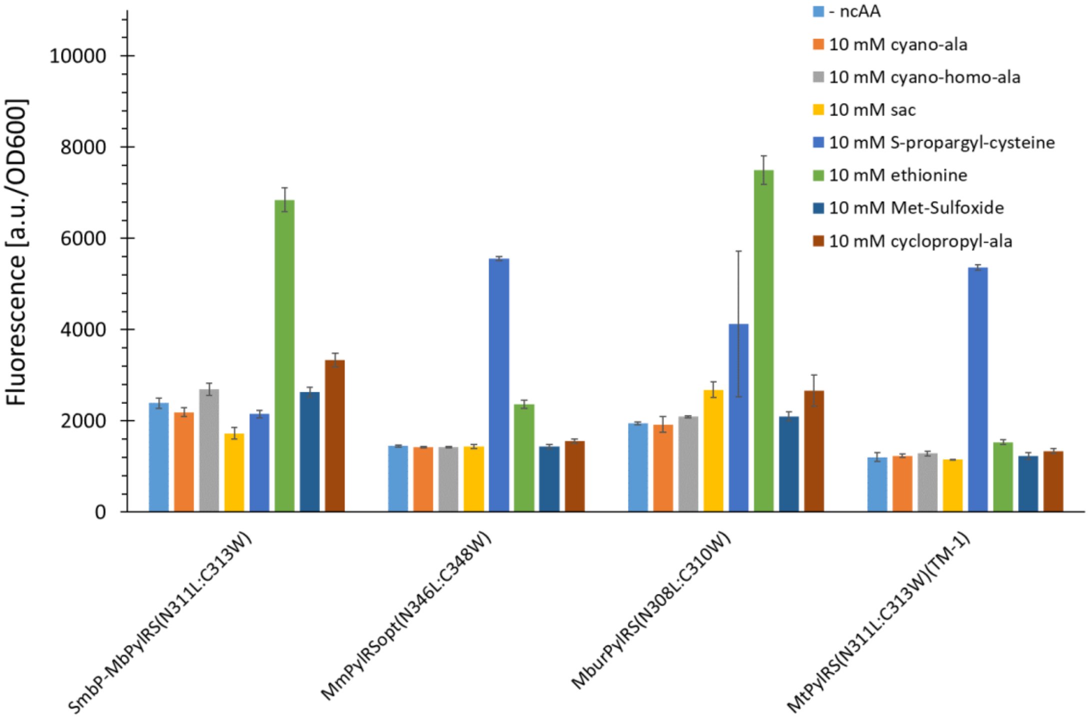

**Figure S53.** Comparison of ncAA (**18**, **19**, **1**, **27**, **25**, **23**, **9**) incorporation efficiency for different PylRS constructs. Fluorescence measurement of intact *E. coli* BL21(DE3) cells producing the SUMO-sfGFP(R2amber) reporter protein. Endpoint measurements after 24 h of incubation. Data including standard deviation represents the mean of three

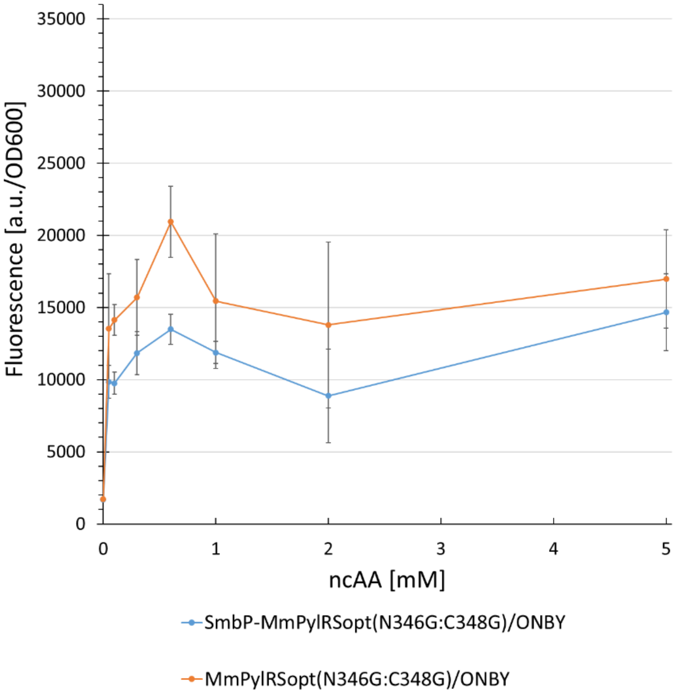

biological replicates. 10 mM ncAAs supplied

**Figure S54.** Concentration-dependent protein production for two different PylRS double Gly mutants. Fluorescence intensity of intact *E. coli* BL21(DE3) cells expressing the SUMO-sfGFP(R2 amber) reporter. Endpoint measurements

after 24 h with different ncAA concentrations (0.05, 0.1, 0.3, 0.6, 1, 2, and 5 mM). Data including standard deviation represents the mean of three biological replicates.

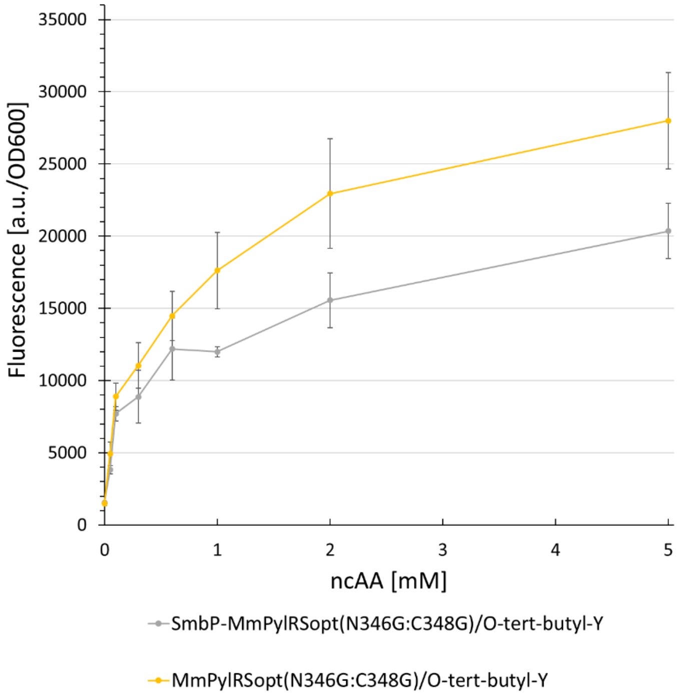

**Figure S55.** Concentration-dependent protein production for two different PylRS double Gly mutants. Fluorescence intensity of intact *E. coli* BL21(DE3) cells expressing the SUMO-sfGFP(R2 amber) reporter. Endpoint measurements after 24 h with different ncAA concentrations (0.05, 0.1, 0.3, 0.6, 1, 2, and 5 mM). Data including standard deviation represents the mean of three biological replicates.

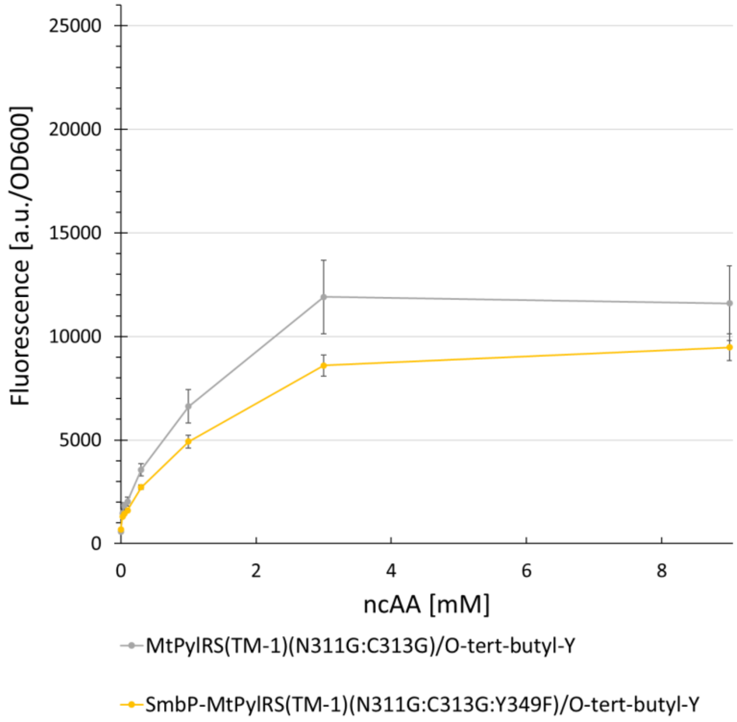

**Figure S56.** Concentration-dependent protein production for two different PylRS double Gly mutants. Fluorescence intensity of intact *E. coli* BL21(DE3) cells expressing the SUMO-sfGFP(R2 amber) reporter. Endpoint measurements

after 24 h with different ncAA concentrations (0.05, 0.1, 0.3, 0.6, 1, 2, and 5 mM). Data including standard deviation represents the mean of three biological replicates.

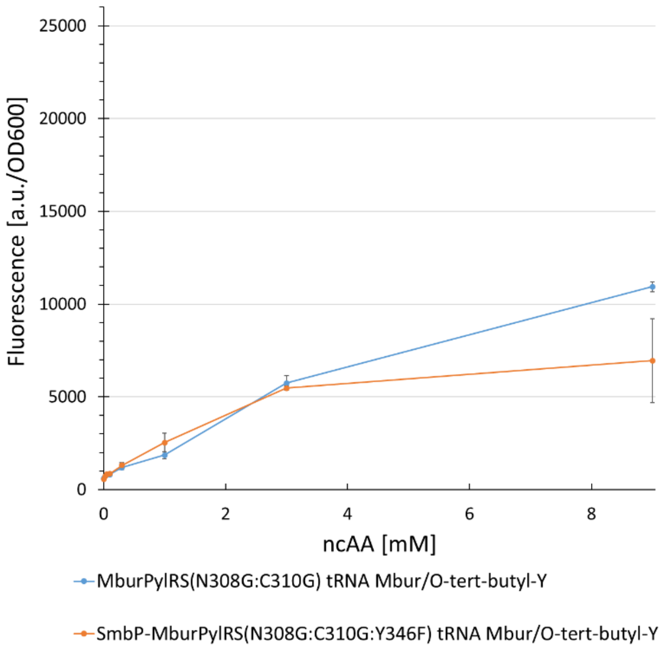

**Figure S57.** Concentration-dependent protein production for two different PylRS double Gly mutants. Fluorescence intensity of intact *E. coli* BL21(DE3) cells expressing the SUMO-sfGFP(R2 amber) reporter. Endpoint measurements after 24 h with different ncAA concentrations (0.05, 0.1, 0.3, 0.6, 1, 2, and 5 mM). Data including standard deviation represents the mean of three biological replicates.

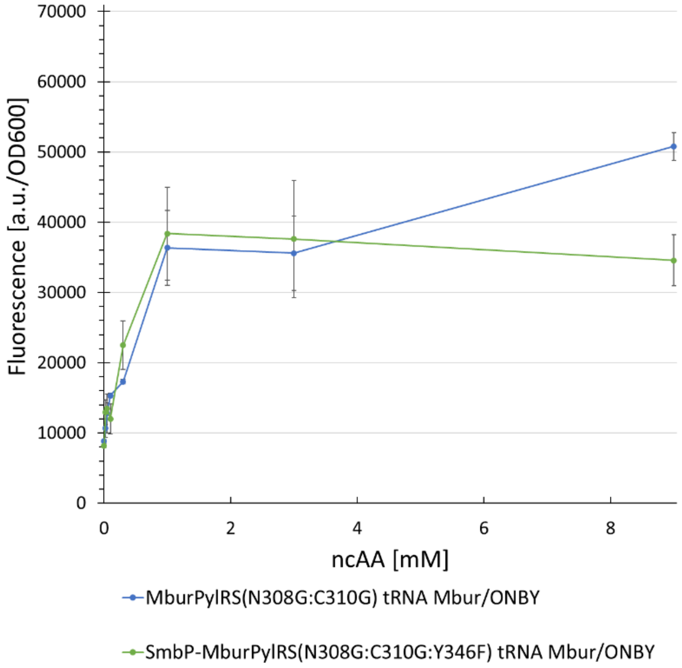

**Figure S58.** Concentration-dependent protein production for two different PylRS double Gly mutants. Fluorescence intensity of intact *E. coli* BL21(DE3) cells expressing the SUMO-sfGFP(R2 amber) reporter. Endpoint measurements

after 24 h with different ncAA concentrations (0.05, 0.1, 0.3, 0.6, 1, 2, and 5 mM). Data including standard deviation

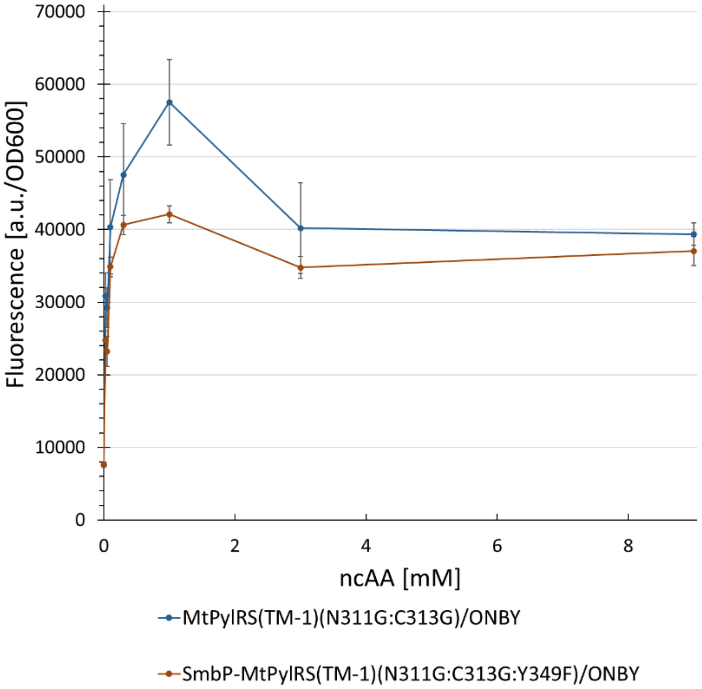

represents the mean of three biological replicates.

**Figure S59.** Concentration-dependent protein production for two different PylRS double Gly mutants. Fluorescence intensity of intact *E. coli* BL21(DE3) cells expressing the SUMO-sfGFP(R2 amber) reporter. Endpoint measurements after 24 h with different ncAA concentrations (0.05, 0.1, 0.3, 0.6, 1, 2, and 5 mM). Data including standard deviation

represents the mean of three biological replicates.

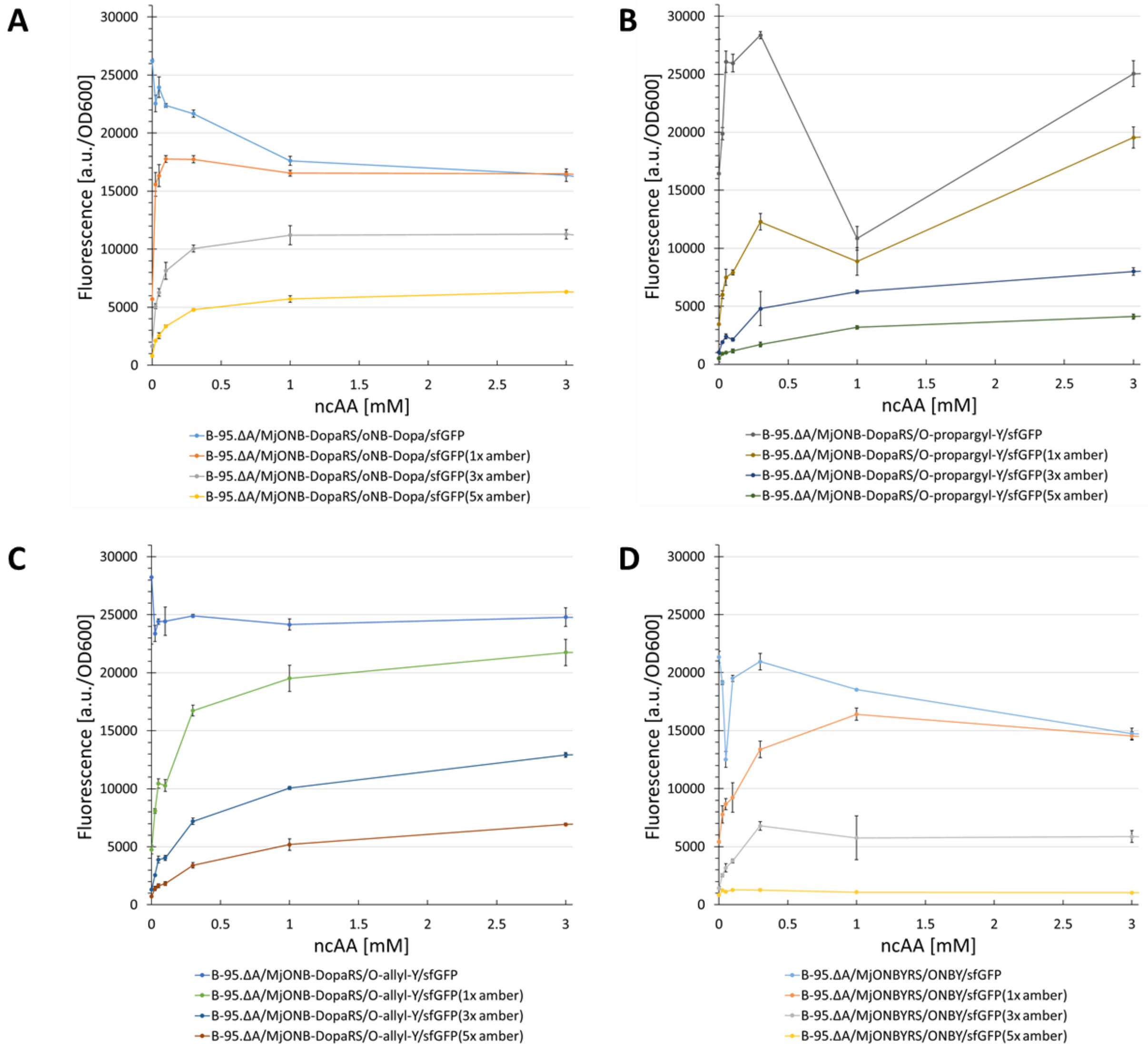

**Figure S60.** Concentration-dependent protein production for four different *Mj*TyrRS/ncAA combinations. Fluorescence intensity of intact B-95.Δ cells expressing the SUMO-sfGFP reporter containing indicated number of stop codons. Endpoint measurements after 24 h with different ncAA concentrations (0.025, 0.05, 0.1, 0.3, 1, 3, and 9 (not shown)

mM). Data including standard deviation represents the mean of three biological replicates.

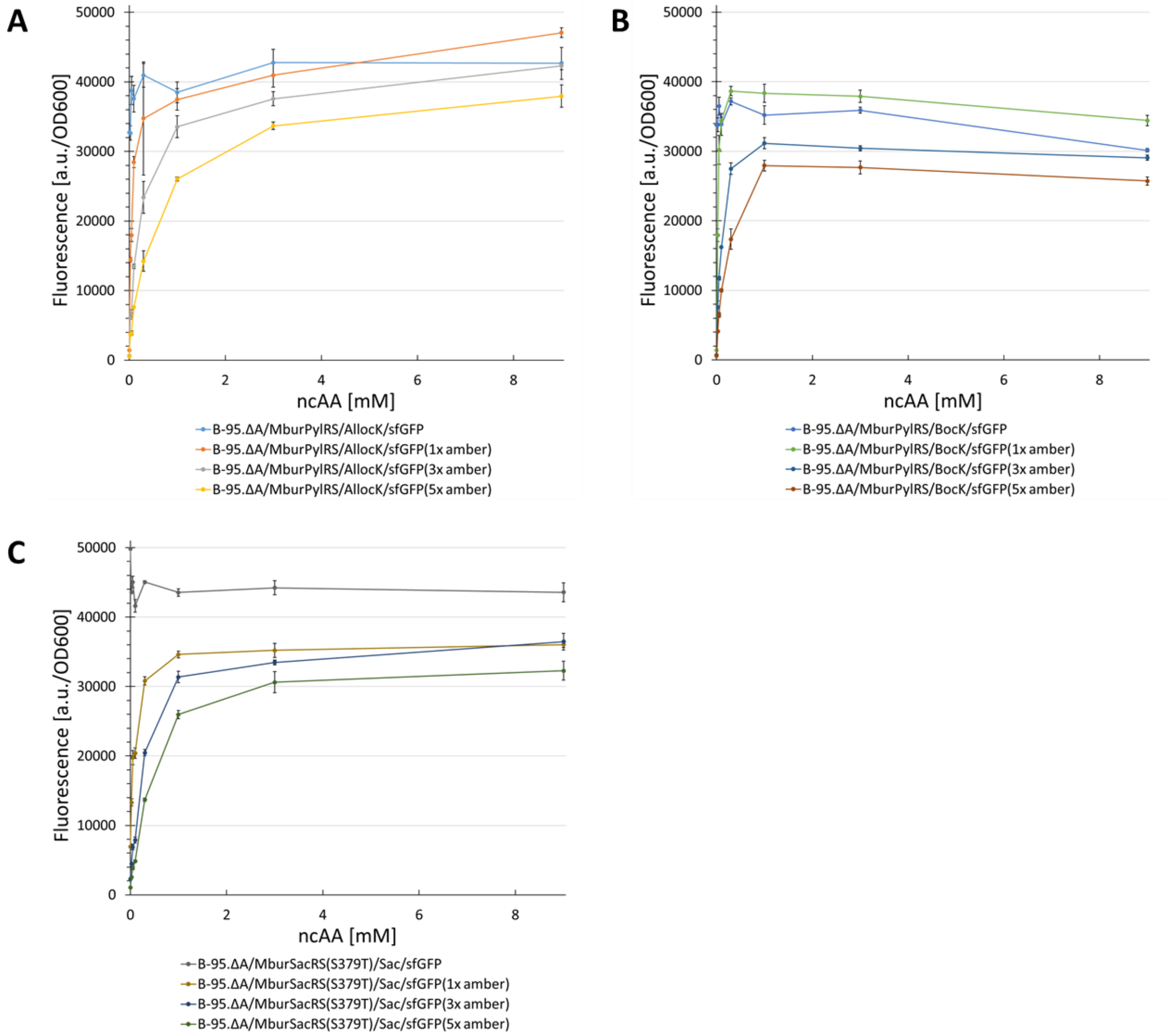

**Figure S61.** Concentration-dependent protein production for three different MburPylRSRS/ncAA combinations. Fluorescence intensity of intact B-95.Δ cells expressing the SUMO-sfGFP reporter containing indicated number of stop codons. Endpoint measurements after 24 h with different ncAA concentrations (0.025, 0.05, 0.1, 0.3, 1, 3, and 9 mM). Data including standard deviation represents the mean of three biological replicates.

#### 2.2 **DNA/RNA Sequences**

##### 2.2.1 Used PylRS

The PylRS amino acid and $tRNA^{Pyl}$ sequences used in this study are found in the supplemental excel sheet. All PylRS sequences were codon optimized for *E. coli*. Sequences for the ΔN PylRS class were taken from the publication of Dunkelmann et al.[15]

##### 2.2.2 Reporter Constructs

1) SUMO-sfGFP(1x amber)-$His_6$ reporter construct with amber codons at position R2

ATGGGCAGCAGCGACTCCGAAGTCAATCAAGAAGCTAAGCCAGAGGTCAAGCCAGAAGTC
AAGCCTGAGACTCACATCAATTTAAAGGTGTCCGATGGATCTTCAGAGATCTTCTTCAAGAT
CAAAAAGACCACTCCTCTGCGTCGTCTGATGGAAGCGTTCGCTAAAAGACAGGGTAAGGA
AATGGACTCCTTAAGATTCTTGTACGACGGTATTAGAATCCAAGCTGATCAGACCCCTGAA
GATTTGGACATGGAGGATAACGATATTATTGAGGCTCATCGCGAACAGATTGGTGGCATGT
AGAAAGGCGAAGAGCTGTTCACTGGTGTCGTCCCTATTCTGGTGGAACTGGATGGTGATG
TCAACGGTCATAAGTTTTCCGTGCGTGGCGAGGGTGAAGGTGACGCAACTAATGGTAAAC
TGACGCTGAAGTTCATCTGTACTACTGGTAAACTGCCGGTACCTTGGCCGACTCTGGTAAC
GACGCTGACTTATGGTGTTCAGTGCTTTGCTCGTTATCCGGACCATATGAAGCAGCATGAC
TTCTTCAAGTCCGCCATGCCGGAAGGCTATGTGCAGGAACGCACGATTTCCTTTAAGGATG
ACGGCACGTACAAAACGCGTGCGGAAGTGAAATTTGAAGGCGATACCCTGGTAAACCGCA
TTGAGCTGAAAGGCATTGACTTTAAAGAAGACGGCAATATCCTGGGCCATAAGCTGGAATA
CAATTTTAACAGCCACAATGTTTACATCACCGCCGATAAACAAAAAAATGGCATTAAAGCGA
ATTTTAAAATTCGCCACAACGTGGAGGATGGCAGCGTGCAGCTGGCTGATCACTACCAGC
AAAACACTCCAATCGGTGATGGTCCTGTTCTGCTGCCAGACAATCACTATCTGAGCACGCA
AAGCGTTCTGTCTAAAGATCCGAACGAGAAACGCGATCATATGGTTCTGCTGGAGTTCGTA
ACCGCAGCGGGCATCACGCATGGTATGGATGAACTGTACAAAAGCGCTCATCATCATCATC
ATCACTAA

2) SUMO-sfGFP(3x amber)-$His_6$ reporter construct with amber codons at positions R2, N39 and K101

ATGGGCAGCAGCGACTCCGAAGTCAATCAAGAAGCTAAGCCAGAGGTCAAGCCAGAAGTC
AAGCCTGAGACTCACATCAATTTAAAGGTGTCCGATGGATCTTCAGAGATCTTCTTCAAGAT
CAAAAAGACCACTCCTCTGCGTCGTCTGATGGAAGCGTTCGCTAAAAGACAGGGTAAGGA
AATGGACTCCTTAAGATTCTTGTACGACGGTATTAGAATCCAAGCTGATCAGACCCCTGAA
GATTTGGACATGGAGGATAACGATATTATTGAGGCTCATCGCGAACAGATTGGTGGCATGT

AGAAAGGCGAAGAGCTGTTCACTGGTGTCGTCCCTATTCTGGTGGAACTGGATGGTGATG
TCAACGGTCATAAGTTTTCCGTGCGTGGCGAGGGTGAAGGTGACGCAACTTAGGGTAAAC
TGACGCTGAAGTTCATCTGTACTACTGGTAAACTGCCGGTACCTTGGCCGACTCTGGTAAC
GACGCTGACTTATGGTGTTCAGTGCTTTGCTCGTTATCCGGACCATATGAAGCAGCATGAC
TTCTTCAAGTCCGCCATGCCGGAAGGCTATGTGCAGGAACGCACGATTTCCTTTTAGGATG
ACGGCACGTACAAAACGCGTGCGGAAGTGAAATTTGAAGGCGATACCCTGGTAAACCGCA
TTGAGCTGAAAGGCATTGACTTTAAAGAAGACGGCAATATCCTGGGCCATAAGCTGGAATA
CAATTTTAACAGCCACAATGTTTACATCACCGCCGATAAACAAAAAAATGGCATTAAAGCGA
ATTTTAAAATTCGCCACAACGTGGAGGATGGCAGCGTGCAGCTGGCTGATCACTACCAGC
AAAACACTCCAATCGGTGATGGTCCTGTTCTGCTGCCAGACAATCACTATCTGAGCACGCA
AAGCGTTCTGTCTAAAGATCCGAACGAGAAACGCGATCATATGGTTCTGCTGGAGTTCGTA
ACCGCAGCGGGCATCACGCATGGTATGGATGAACTGTACAAAAGCGCTCATCATCATCATC
ATCACTAA

3) SUMO-sfGFP(5x amber)-$His_6$ reporter construct with amber codons at positions R2, N39, K101, E132 and D190

ATGGGCAGCAGCGACTCCGAAGTCAATCAAGAAGCTAAGCCAGAGGTCAAGCCAGAAGTC
AAGCCTGAGACTCACATCAATTTAAAGGTGTCCGATGGATCTTCAGAGATCTTCTTCAAGAT
CAAAAAGACCACTCCTCTGCGTCGTCTGATGGAAGCGTTCGCTAAAAGACAGGGTAAGGA
AATGGACTCCTTAAGATTCTTGTACGACGGTATTAGAATCCAAGCTGATCAGACCCCTGAA
GATTTGGACATGGAGGATAACGATATTATTGAGGCTCATCGCGAACAGATTGGTGGCATGT
AGAAAGGCGAAGAGCTGTTCACTGGTGTCGTCCCTATTCTGGTGGAACTGGATGGTGATG
TCAACGGTCATAAGTTTTCCGTGCGTGGCGAGGGTGAAGGTGACGCAACTTAGGGTAAAC
TGACGCTGAAGTTCATCTGTACTACTGGTAAACTGCCGGTACCTTGGCCGACTCTGGTAAC
GACGCTGACTTATGGTGTTCAGTGCTTTGCTCGTTATCCGGACCATATGAAGCAGCATGAC
TTCTTCAAGTCCGCCATGCCGGAAGGCTATGTGCAGGAACGCACGATTTCCTTTTAGGATG
ACGGCACGTACAAAACGCGTGCGGAAGTGAAATTTGAAGGCGATACCCTGGTAAACCGCA
TTGAGCTGAAAGGCATTGACTTTAAATAGGACGGCAATATCCTGGGCCATAAGCTGGAATA
CAATTTTAACAGCCACAATGTTTACATCACCGCCGATAAACAAAAAAATGGCATTAAAGCGA
ATTTTAAAATTCGCCACAACGTGGAGGATGGCAGCGTGCAGCTGGCTGATCACTACCAGC
AAAACACTCCAATCGGTTAGGGTCCTGTTCTGCTGCCAGACAATCACTATCTGAGCACGCA
AAGCGTTCTGTCTAAAGATCCGAACGAGAAACGCGATCATATGGTTCTGCTGGAGTTCGTA
ACCGCAGCGGGCATCACGCATGGTATGGATGAACTGTACAAAAGCGCTCATCATCATCATC
ATCACTAA

#### 2.3 tRNA Sequences

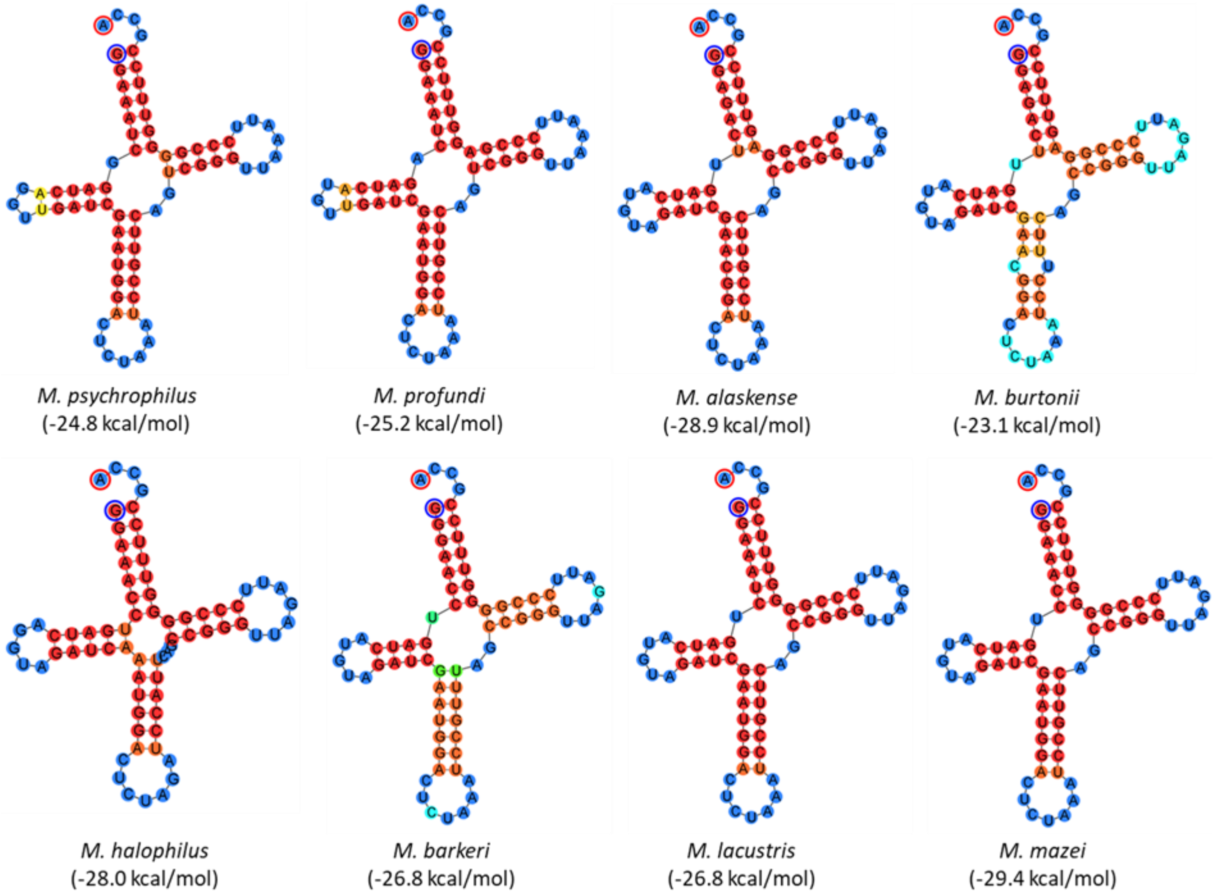

**Figure S62.** Selection of $tRNA^{Pyl}$. ΔG values in brackets, calculated at 37 °C. The nucleotide binding probability is indicated by color; red = high, green = middle and blue = low. The fold and free energy prediction was performed with the Geneious software (version 7.1.9), which uses the ViennaRNA Package.[16] The nucleotide with blue a circle is the 5' end and with a red circle the 3' end.

#### 2.4 Deconvoluted ESI-MS Spectra of intact ncAA-containing protein variants

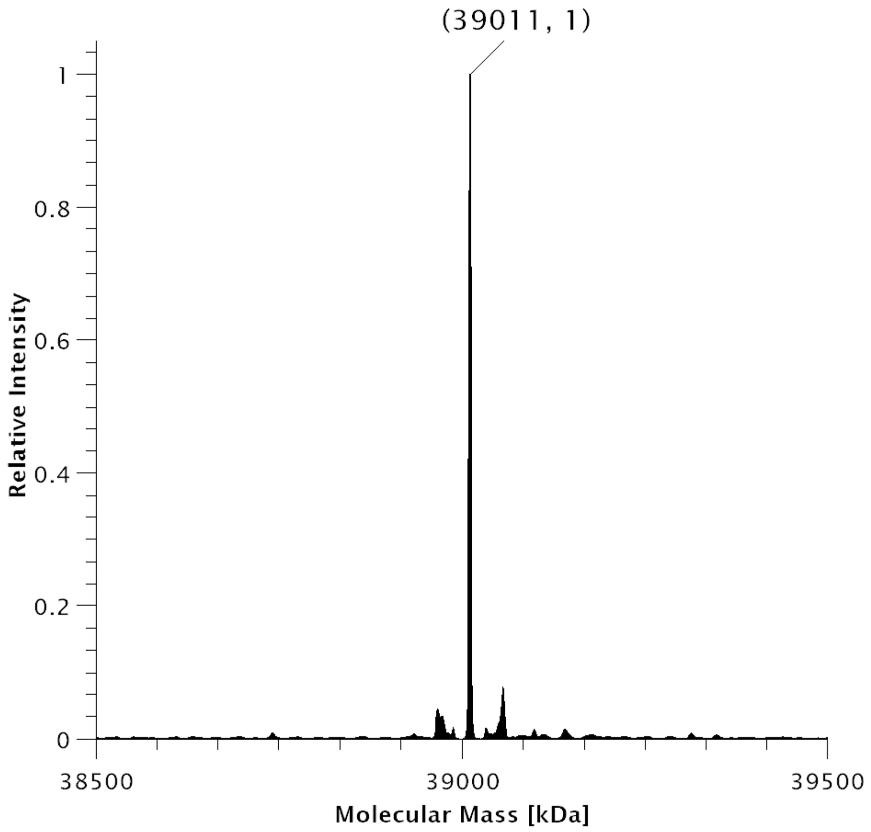

**Figure S63**. Deconvoluted ESI-MS spectrum of SUMO-sfGFP(1x(**38**))-His$_6$ production in *E. coli* BL21(DE3) with co-expression of *Mbur*SacRS(S379T).Expected protein mass: 39011 Da. Observed mass: 39011 Da.

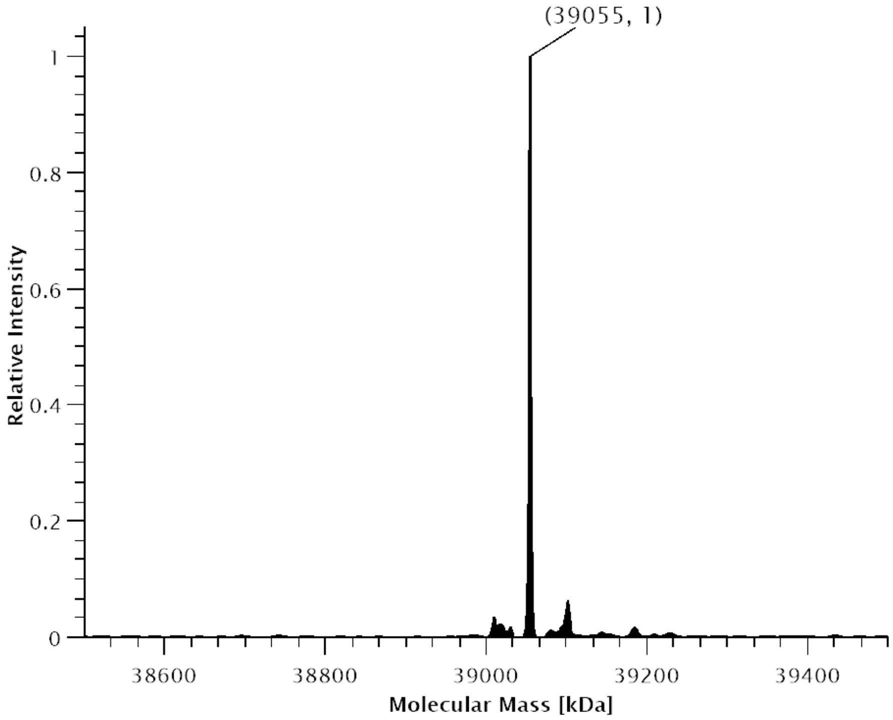

**Figure S64.** Deconvoluted ESI-MS spectrum of SUMO-sfGFP(3x **38**)-His$_6$ production in *E. coli* BL21(DE3) with co-expression of *Mbur*SacRS(S379T).Expected protein mass: 39055 Da. Observed mass: 39055 Da.

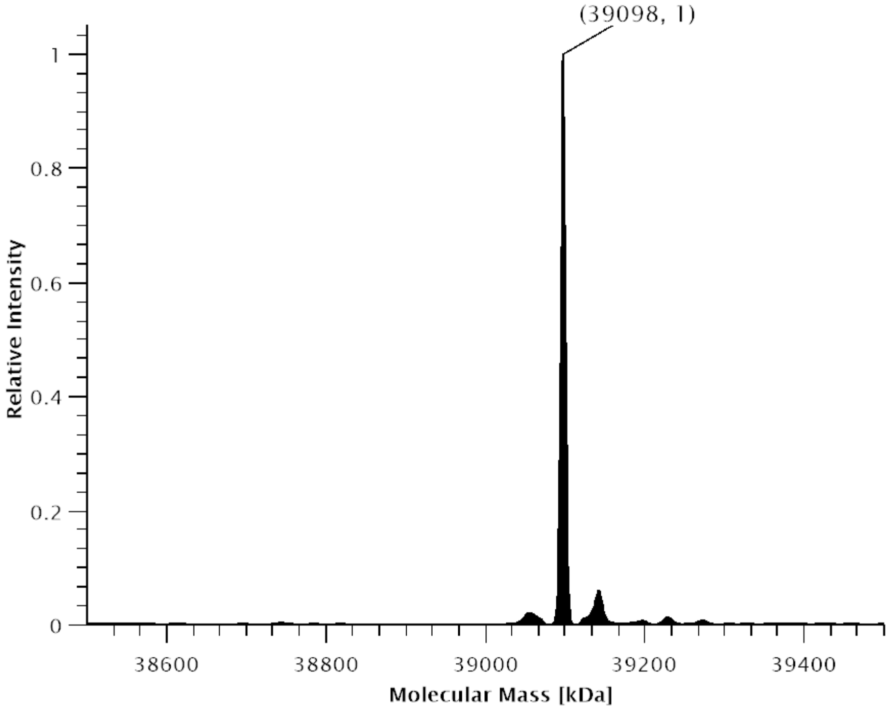

**Figure S65.** Deconvoluted ESI-MS spectrum of SUMO-sfGFP(5x(**38**)-His$_6$ production in *E. coli* BL21(DE3) with co-expression of *Mbur*SacRS(S379T).Expected protein mass: 39098 Da. Observed mass: 39098 Da.

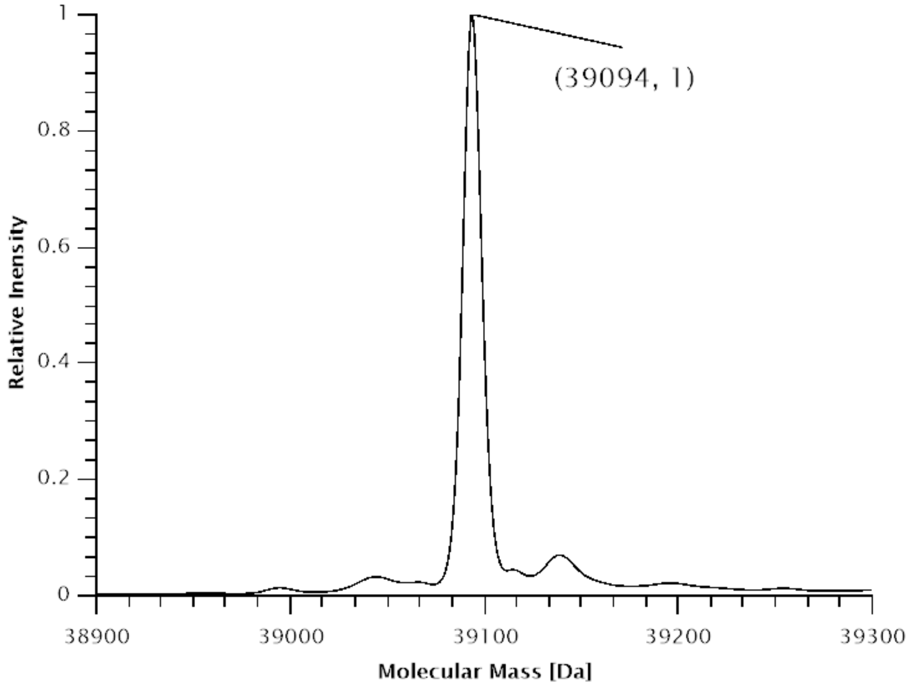

**Figure S66.** Deconvoluted ESI-MS spectrum of SUMO-sfGFP(1x(**1**))-His$_6$ production in *E. coli* BL21(DE3) with co-expression of *Mm*PylRS.Expected protein mass: 39096 Da. Observed mass: 39094 Da.

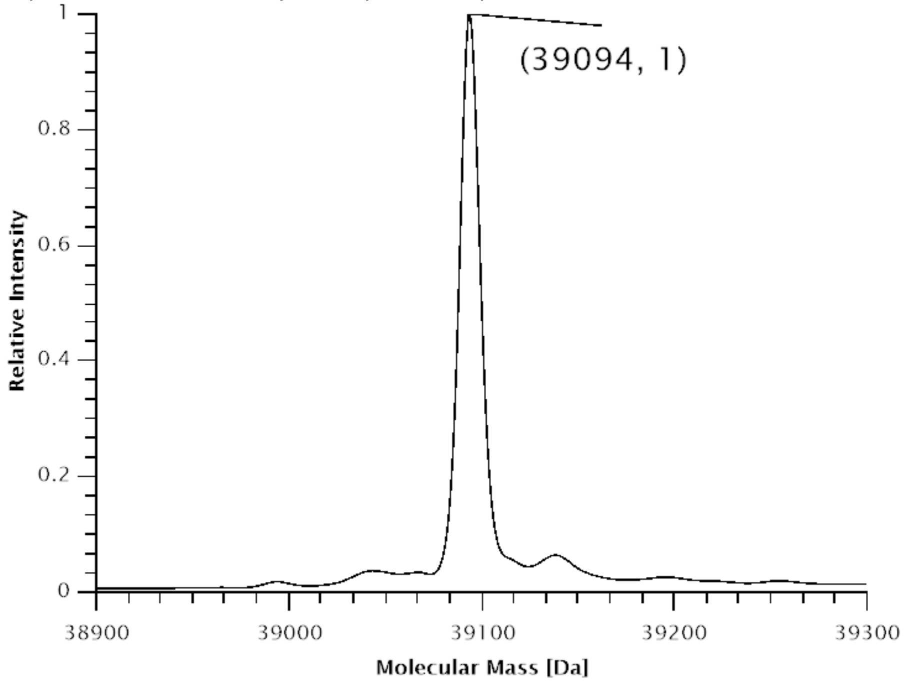

**Figure S67.** Deconvoluted ESI-MS spectrum of SUMO-sfGFP(1x(**1**))-$His_6$ production in *E. coli* BL21(DE3) with co-expression of *Mbur*PylRS.Expected protein mass: 39096 Da. Observed mass: 39094 Da.

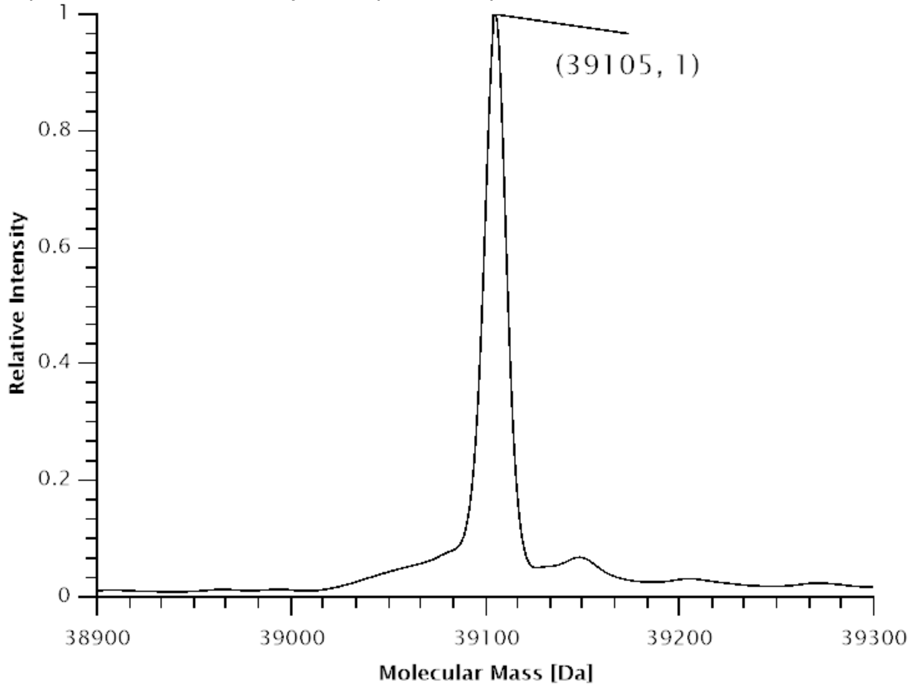

**Figure S68.** Deconvoluted ESI-MS spectrum of SUMO-sfGFP(1x(**4**))-$His_6$ production in *E. coli* BL21(DE3) with co-expression of *Mm*PylRS.Expected protein mass: 39099 Da. Observed mass: 39095 Da.

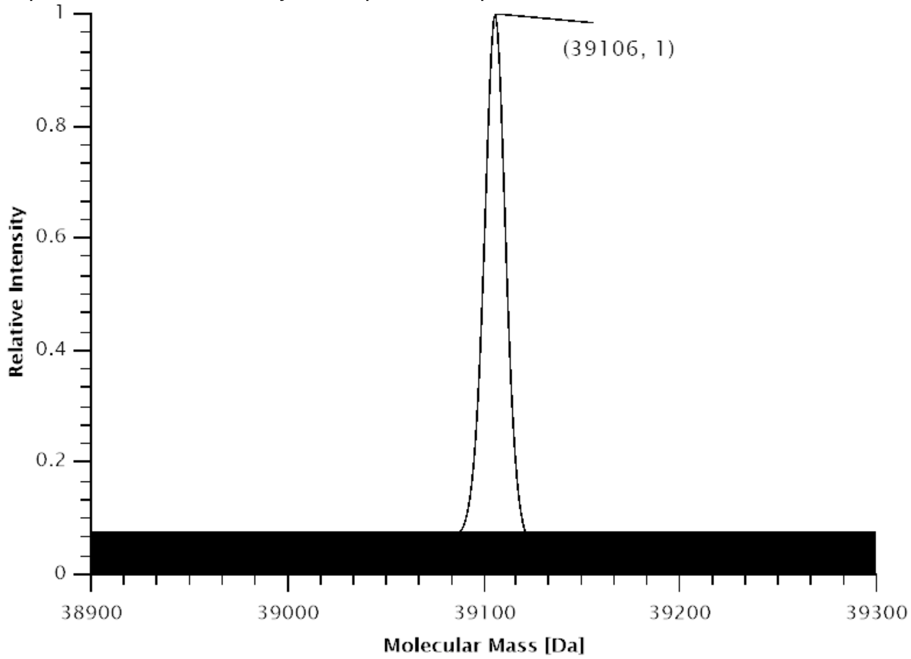

**Figure S69.** Deconvoluted ESI-MS spectrum of SUMO-sfGFP(1x(**4**))-His$_6$ production in *E. coli* BL21(DE3) with co-expression of *Mbur*PylRS.Expected protein mass: 39099 Da. Observed mass: 39096 Da.

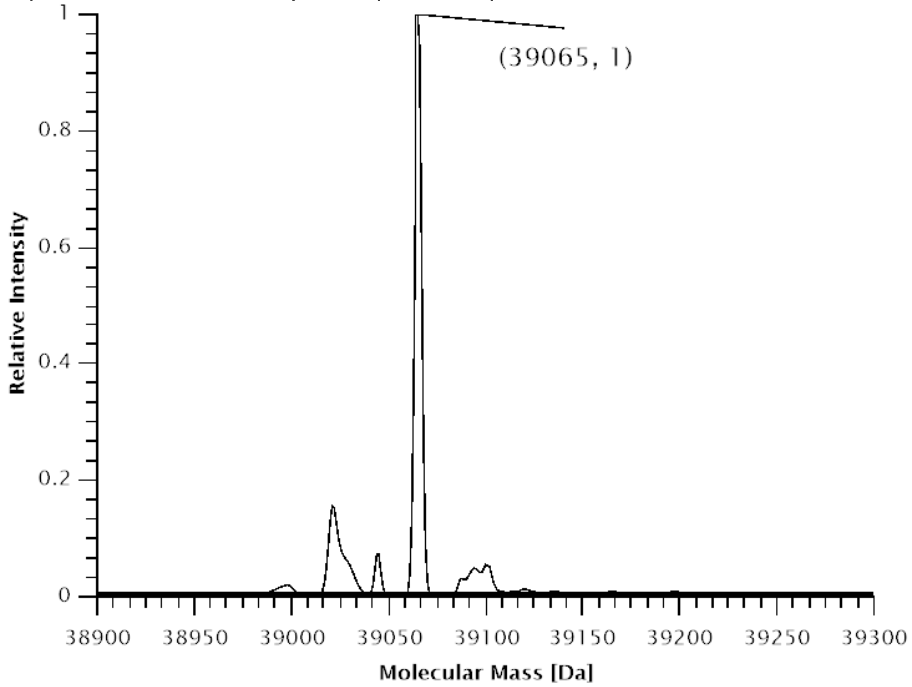

**Figure S70.** Deconvoluted ESI-MS spectrum of SUMO-sfGFP(1x(**9**))-His$_6$ production in *E. coli* BL21(DE3) with co-expression of *Mm*PylRS.Expected protein mass: 39069 Da. Observed mass: 39065 Da.

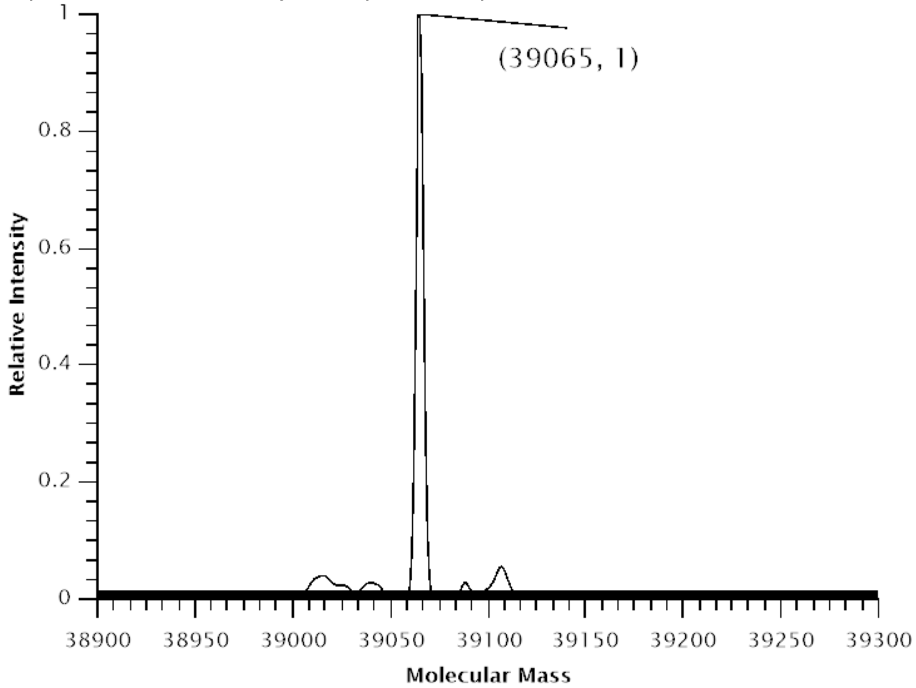

**Figure S71.** Deconvoluted ESI-MS spectrum of SUMO-sfGFP(1x(**9**))-$His_6$ production in *E. coli* BL21(DE3) with co-expression of *Mbur*PylRS.Expected protein mass: 39069 Da. Observed mass: 39065 Da.

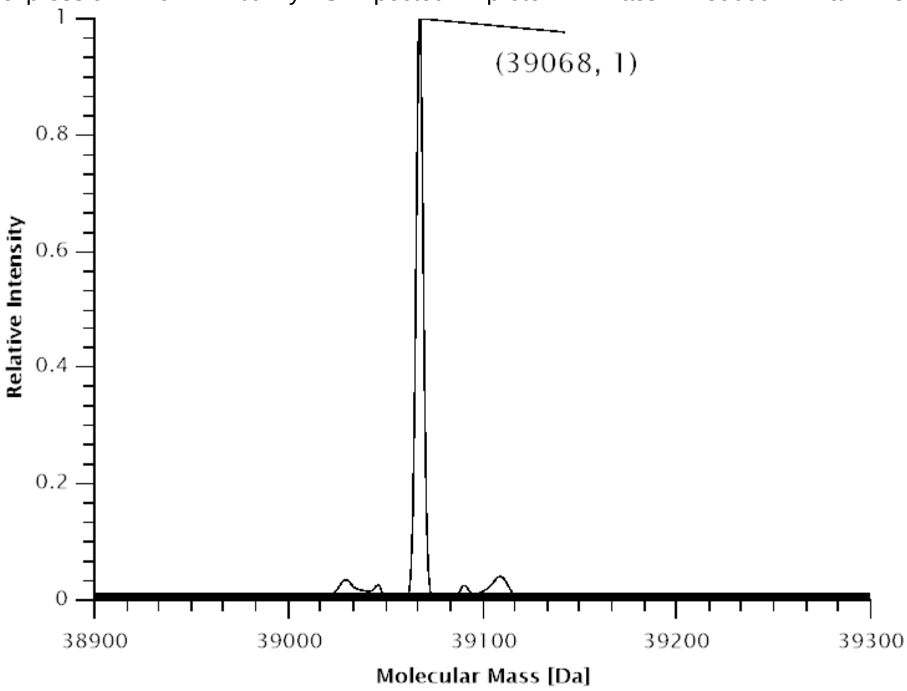

**Figure S72.** Deconvoluted ESI-MS spectrum of SUMO-sfGFP(1x(**11**))-$His_6$ production in *E. coli* BL21(DE3) with co-expression of *Mm*PylRS.Expected protein mass: 39071 Da. Observed mass: 39068 Da.

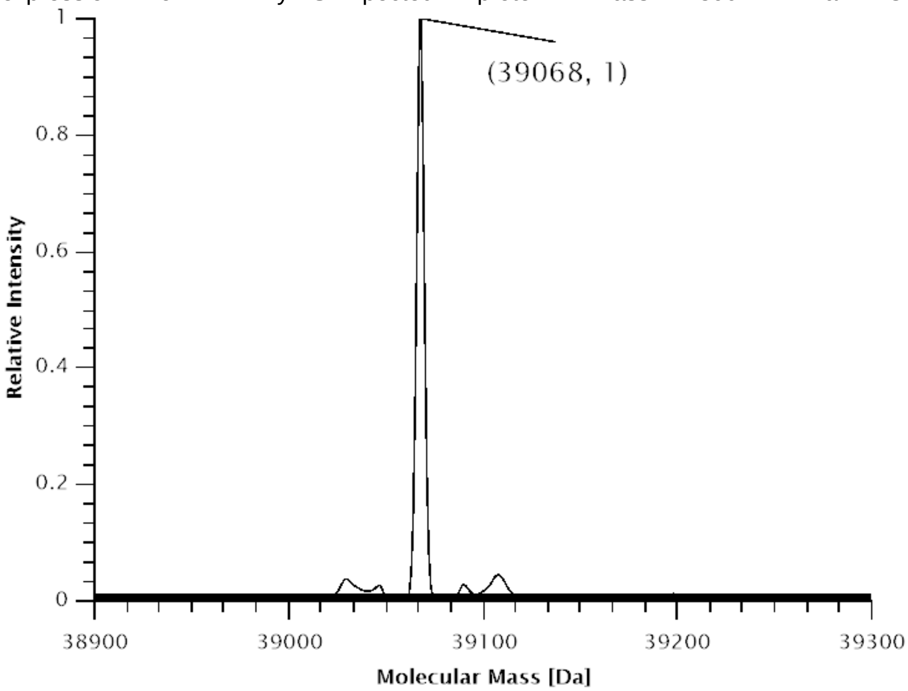

**Figure S73.** Deconvoluted ESI-MS spectrum of SUMO-sfGFP(1x(**11**))-His6 production in *E. coli* BL21(DE3) with co-expression of *Mbur*PylRS.Expected protein mass: 39071 Da. Observed mass: 39068 Da.

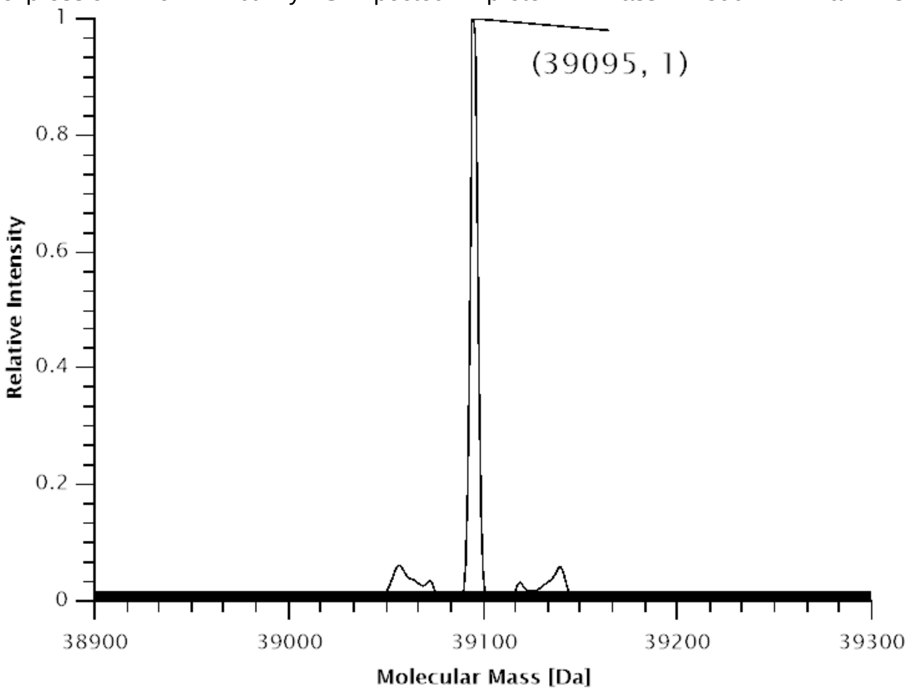

**Figure S74.** Deconvoluted ESI-MS spectrum of SUMO-sfGFP(1x(**13**))-His6 production in *E. coli* BL21(DE3) with co-expression of *Mm*PylRS.Expected protein mass: 39099 Da. Observed mass: 39095 Da.

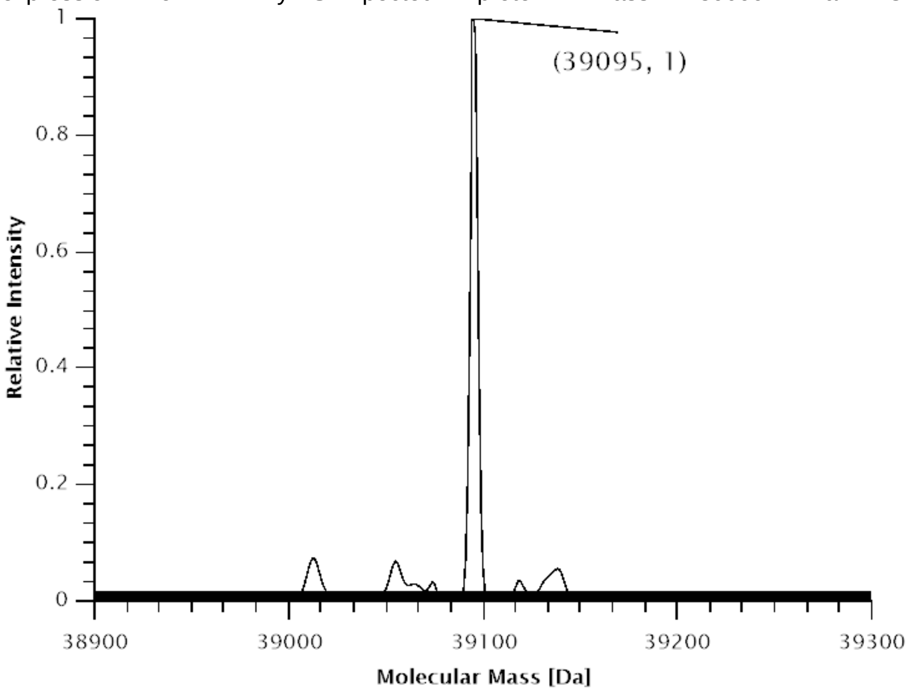

**Figure S75.** Deconvoluted ESI-MS spectrum of SUMO-sfGFP(1x(**13**))-$His_6$ production in *E. coli* BL21(DE3) with co-expression of *Mbur*PylRS.Expected protein mass: 39099 Da. Observed mass: 39095 Da.

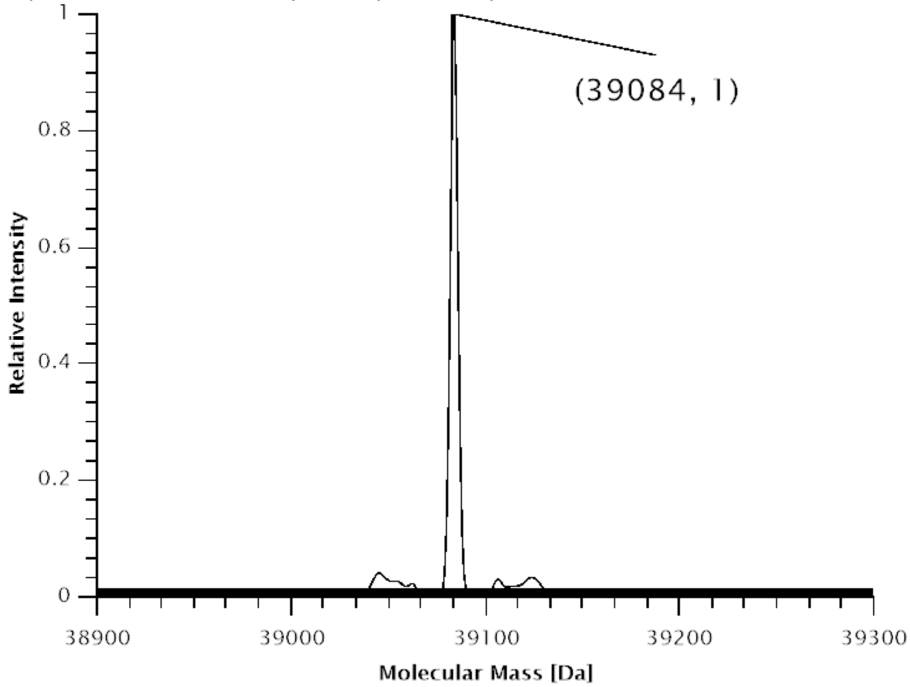

**Figure S76.** Deconvoluted ESI-MS spectrum of SUMO-sfGFP(1x(**8**))-$His_6$ production in *E. coli* BL21(DE3) with co-expression of *Mm*PylRS.Expected protein mass: 39087 Da. Observed mass: 39084 Da.

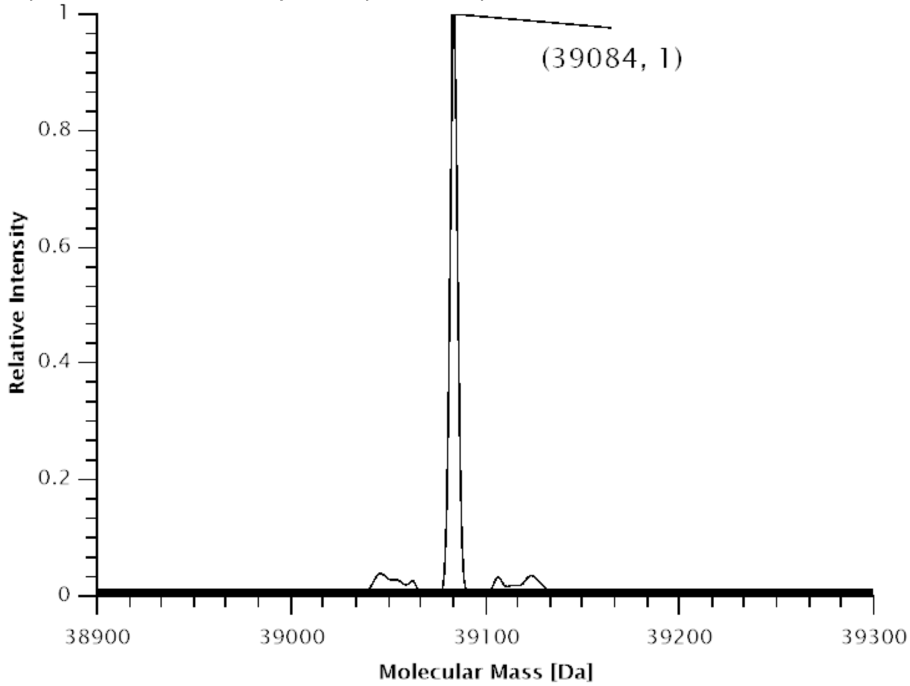

**Figure S77.** Deconvoluted ESI-MS spectrum of SUMO-sfGFP(1x(**8**))-His$_6$ production in *E. coli* BL21(DE3) with co-expression of *Mbur*PylRS.Expected protein mass: 39087 Da. Observed mass: 39084 Da.

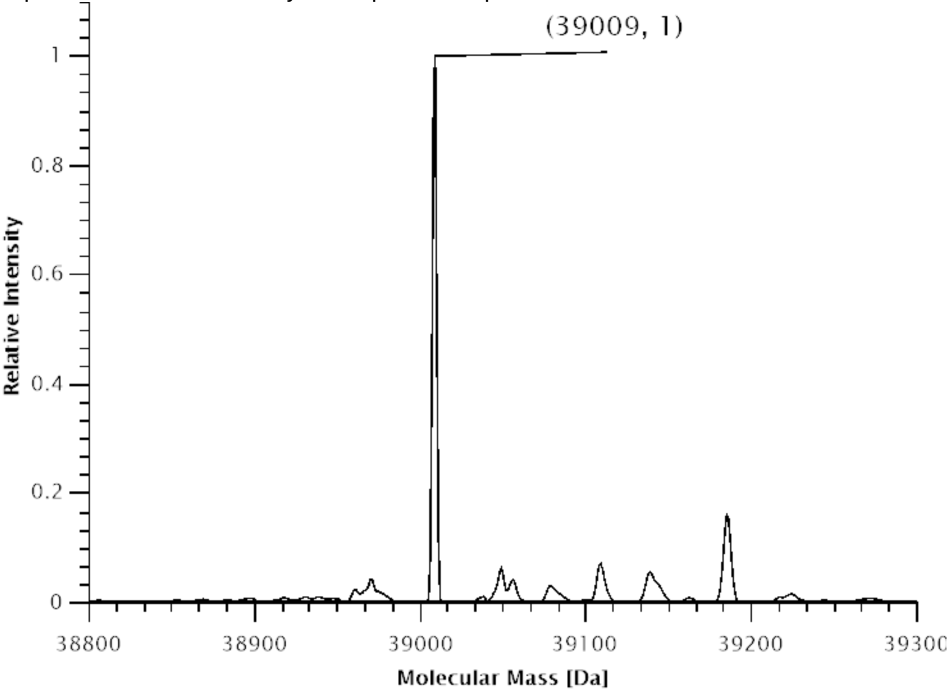

**Figure S78.** Deconvoluted ESI-MS spectrum of SUMO-sfGFP(1x(**35**))-His$_6$ production in *E. coli* B95.ΔA with co-expression of SmbP-*Mb*SacRS(S382T).Expected protein mass: 39008.9 Da. Observed mass: 39009 Da.

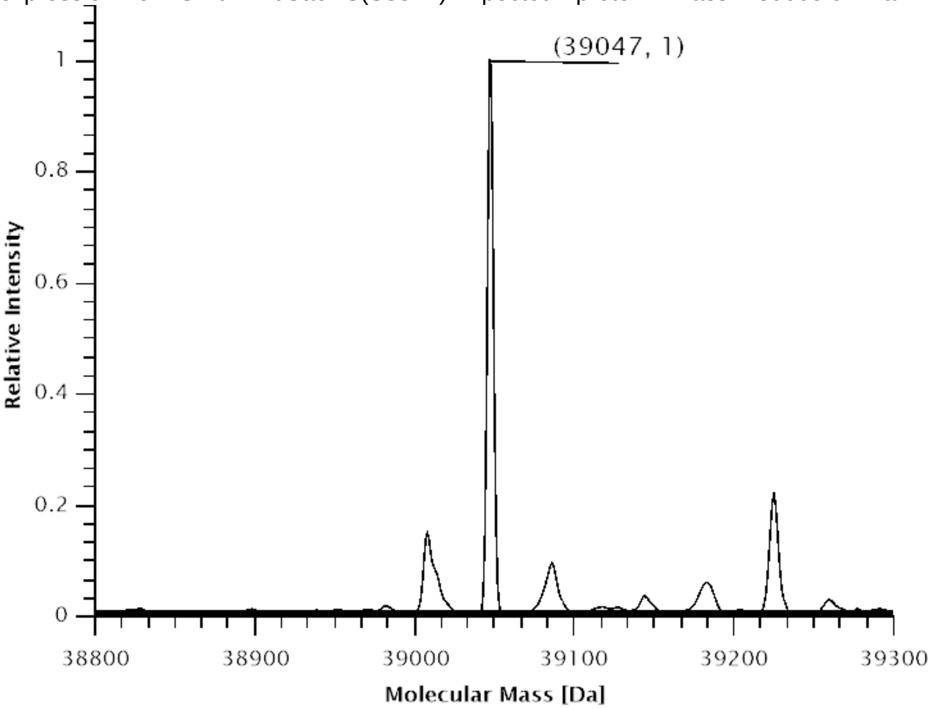

**Figure S79.** Deconvoluted ESI-MS spectrum of SUMO-sfGFP(3x(**35**))-His$_6$ production in *E. coli* B95.ΔA with co-expression of SmbP-*Mb*SacRS(S382T).Expected protein mass: 39049.0 Da. Observed mass: 39047 Da.

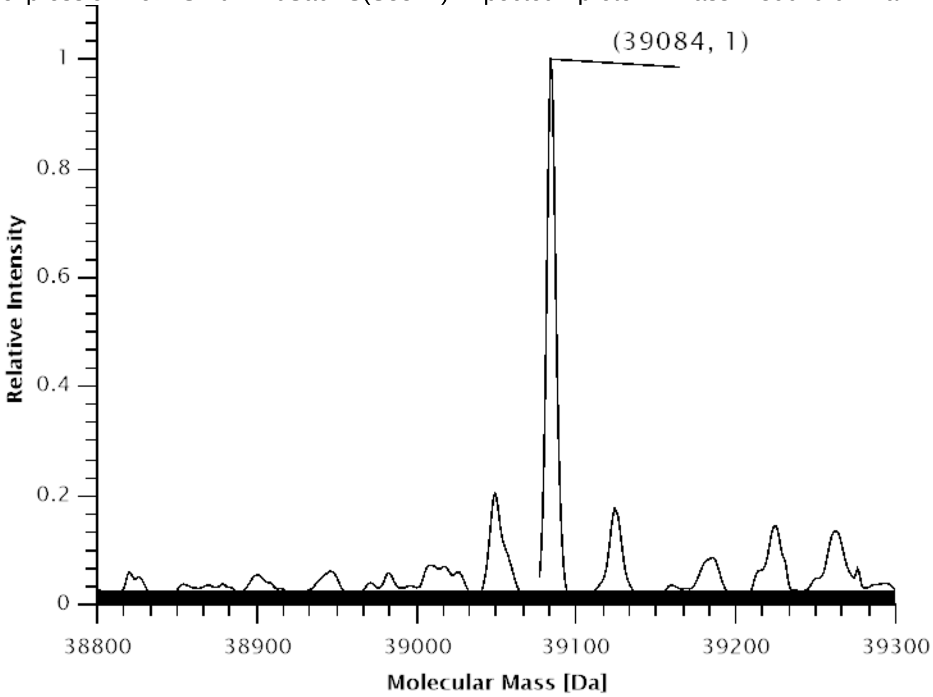

**Figure S80.** Deconvoluted ESI-MS spectrum of SUMO-sfGFP(5x(**35**))-His$_6$ production in *E. coli* B95.ΔA with co-expression of SmbP-*Mb*SacRS(S382T).Expected protein mass: 39087.2 Da. Observed mass: 39084 Da.

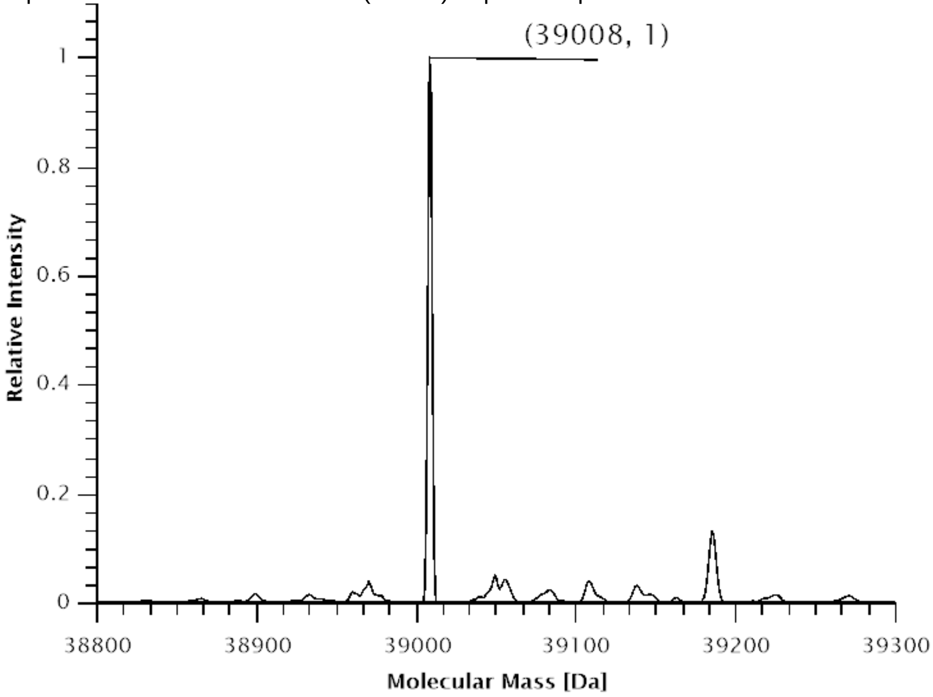

**Figure S81.** Deconvoluted ESI-MS spectrum of SUMO-sfGFP(1x(**35**))-$His_6$ production in *E. coli* B95.ΔA with co-expression of *Mbur*SacRS(S379T).Expected protein mass: 39008.9 Da. Observed mass: 39008 Da.

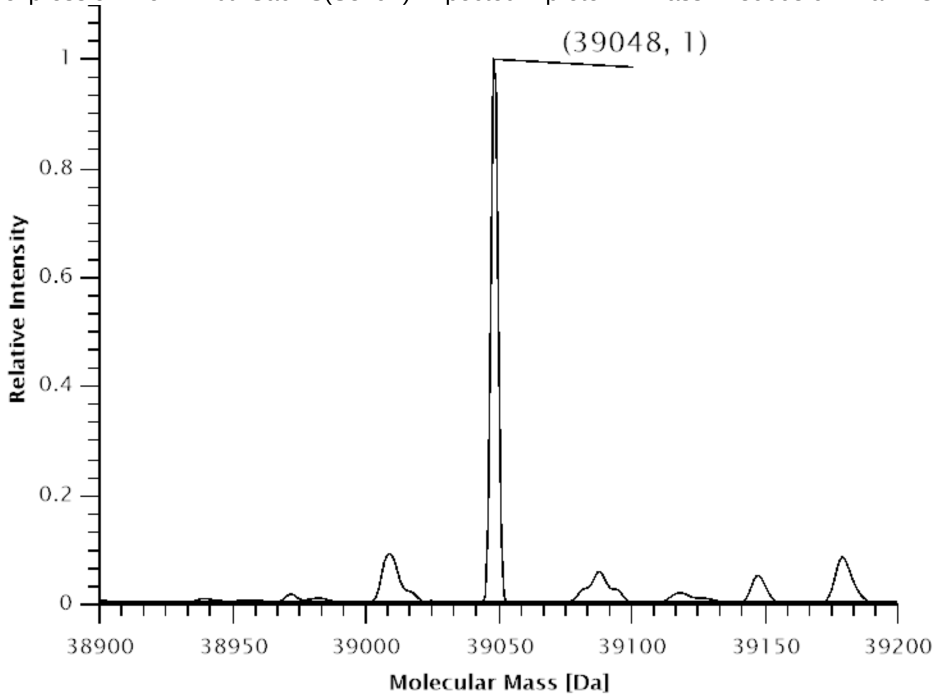

**Figure S82.** Deconvoluted ESI-MS spectrum of SUMO-sfGFP(3x(**35**))-$His_6$ production in *E. coli* B95.ΔA with co-expression of *Mbur*SacRS(S379T).Expected protein mass: 39049.0 Da. Observed mass: 39048 Da.

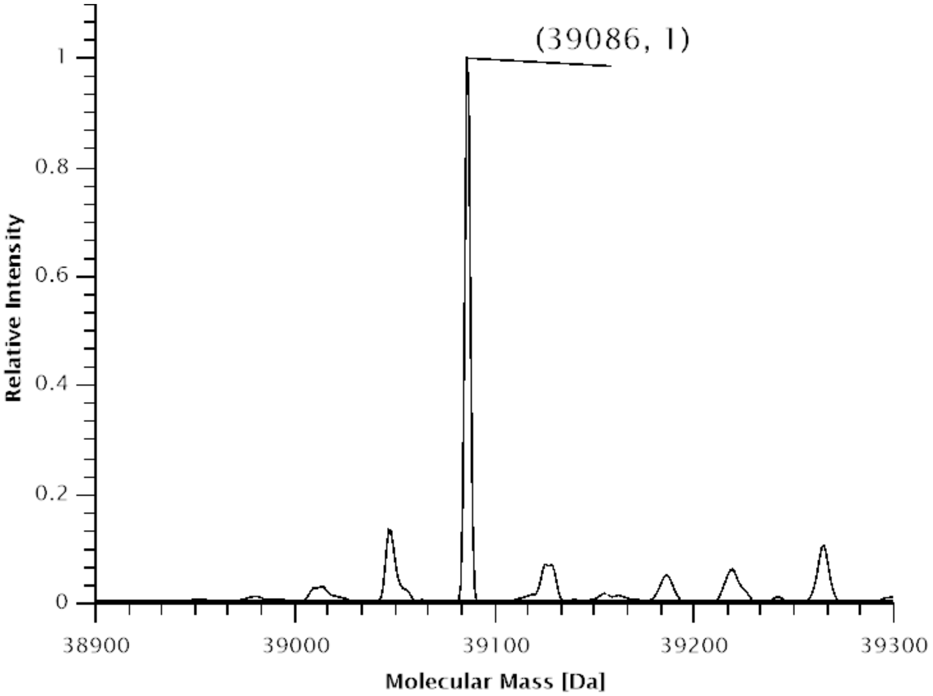

**Figure S83.** Deconvoluted ESI-MS spectrum of SUMO-sfGFP(5x(**35**))-$His_6$ production in *E. coli* B95.ΔA with co-expression of *Mbur*SacRS(S379T).Expected protein mass: 39087.2 Da. Observed mass: 39086 Da.

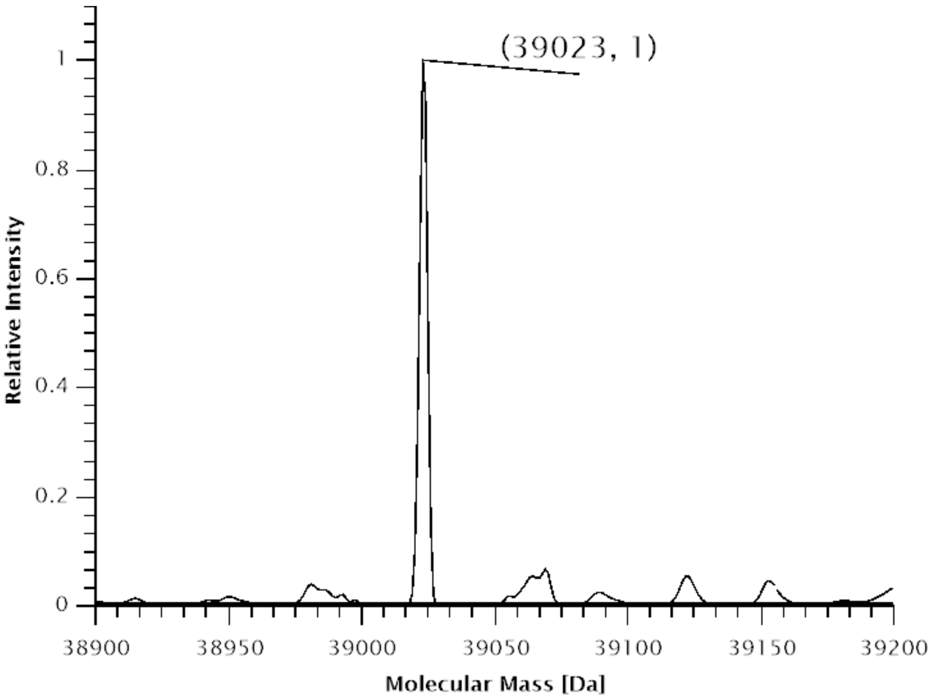

**Figure S84.** Deconvoluted ESI-MS spectrum of SUMO-sfGFP-His$_6$ production in *E. coli* B95.ΔA with co-expression of SmbP-*Mb*SacRS(S382T). Expected protein mass: 39023.9 Da. Observed mass: 39023 Da.

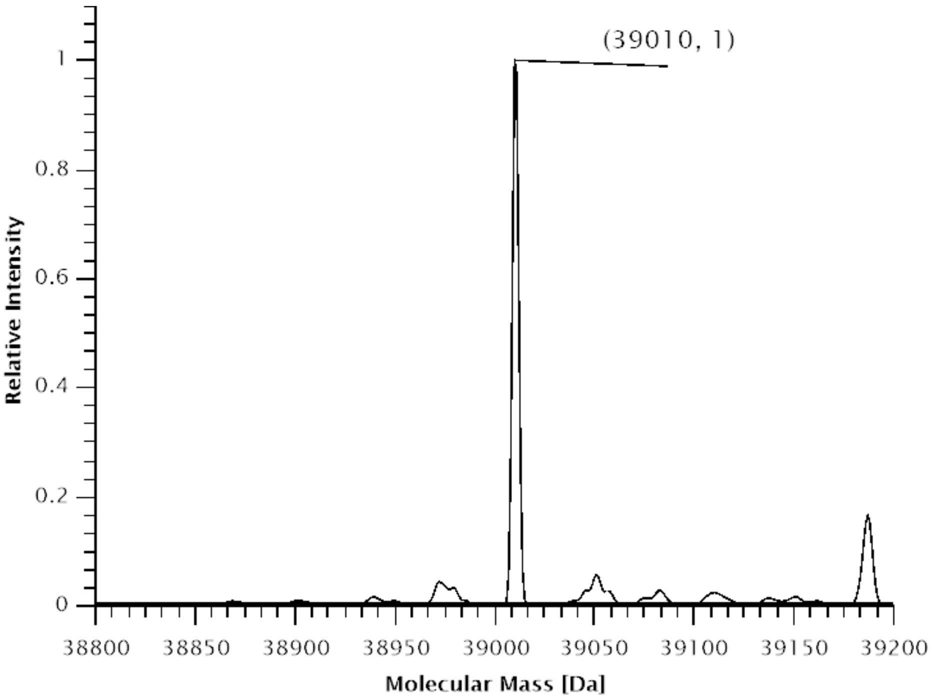

**Figure S85.** Deconvoluted ESI-MS spectrum of SUMO-sfGFP(1x(**38**))-His$_6$ production in *E. coli* B95.ΔA with co-expression of SmbP-*Mb*SacRS(S382T).Expected protein mass: 39010.9 Da. Observed mass: 39010 Da.

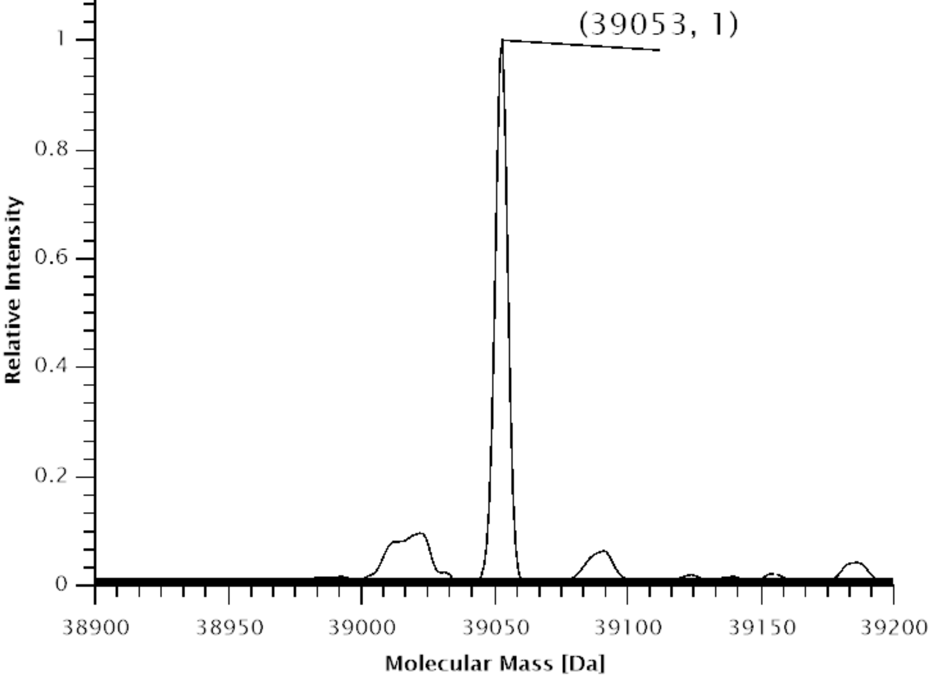

**Figure S86.** Deconvoluted ESI-MS spectrum of SUMO-sfGFP(3x(**38**))-His$_6$ production in *E. coli* B95.ΔA with co-expression of SmbP-*Mb*SacRS(S382T).Expected protein mass: 39055.2 Da. Observed mass: 39053 Da.

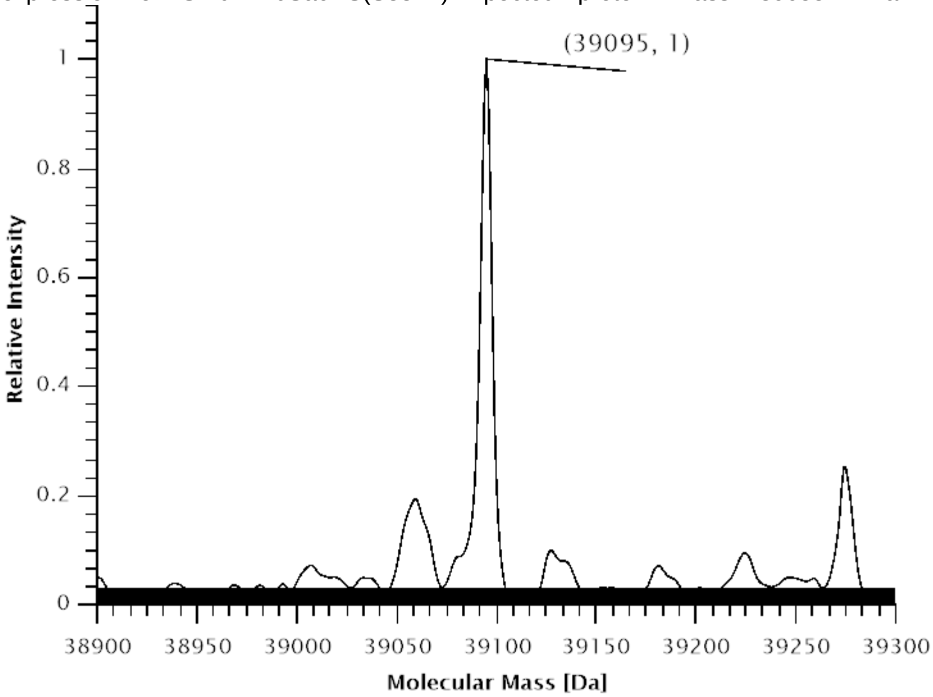

**Figure S87.** Deconvoluted ESI-MS spectrum of SUMO-sfGFP(5x(**38**))-His$_6$ production in *E. coli* B95.ΔA with co-expression of SmbP-*Mb*SacRS(S382T).Expected protein mass: 39097.5 Da. Observed mass: 39095 Da.

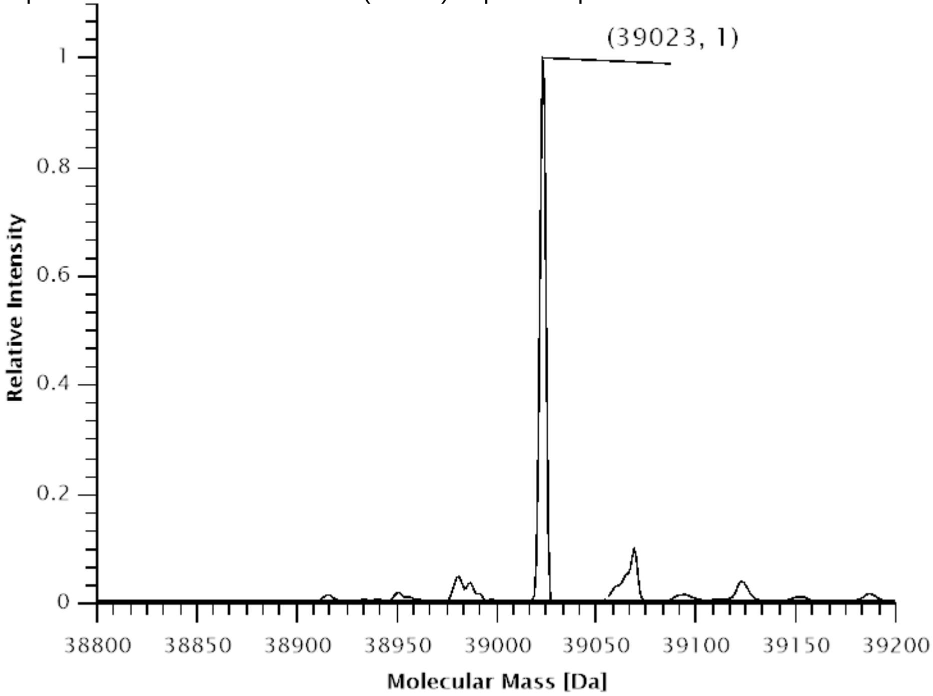

**Figure S88.** Deconvoluted ESI-MS spectrum of SUMO-sfGFP-His6 production in *E. coli* B95.ΔA with co-expression of *Mbur*SacRS(S379T).Expected protein mass: 39023.9 Da. Observed mass: 39023 Da.

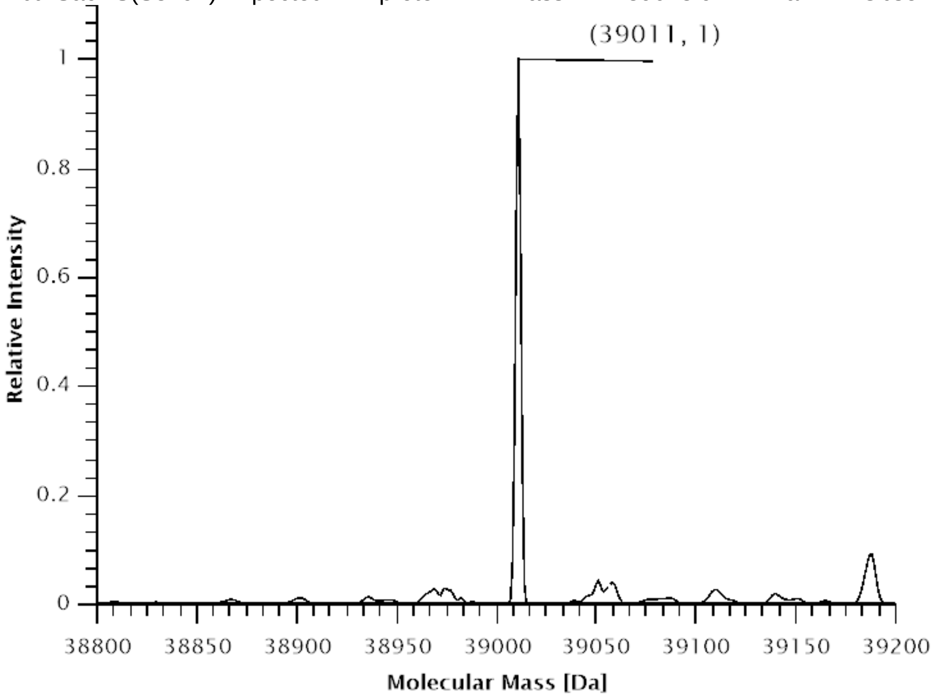

**Figure S89.** Deconvoluted ESI-MS spectrum of SUMO-sfGFP(1x(**38**))-His6 production in *E. coli* B95.ΔA with co-expression of *Mbur*SacRS(S379T).Expected protein mass: 39010.9 Da. Observed mass: 39011 Da.

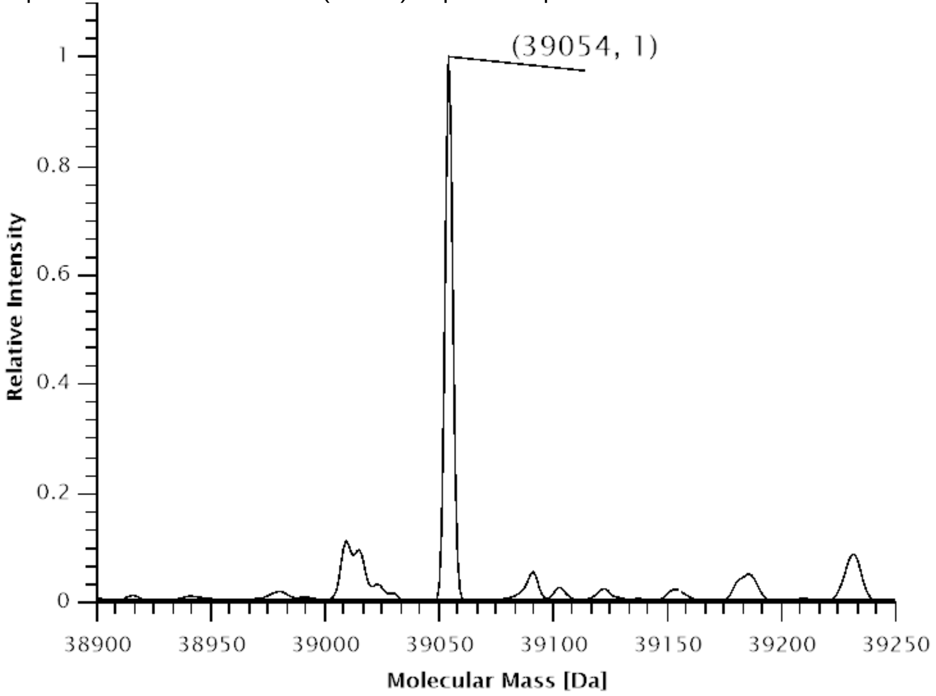

**Figure S90.** Deconvoluted ESI-MS spectrum of SUMO-sfGFP(3x(**38**))-His$_6$ production in *E. coli* B95.ΔA with co-expression of *Mbur*SacRS(S379T).Expected protein mass: 39055.2 Da. Observed mass: 39054 Da.

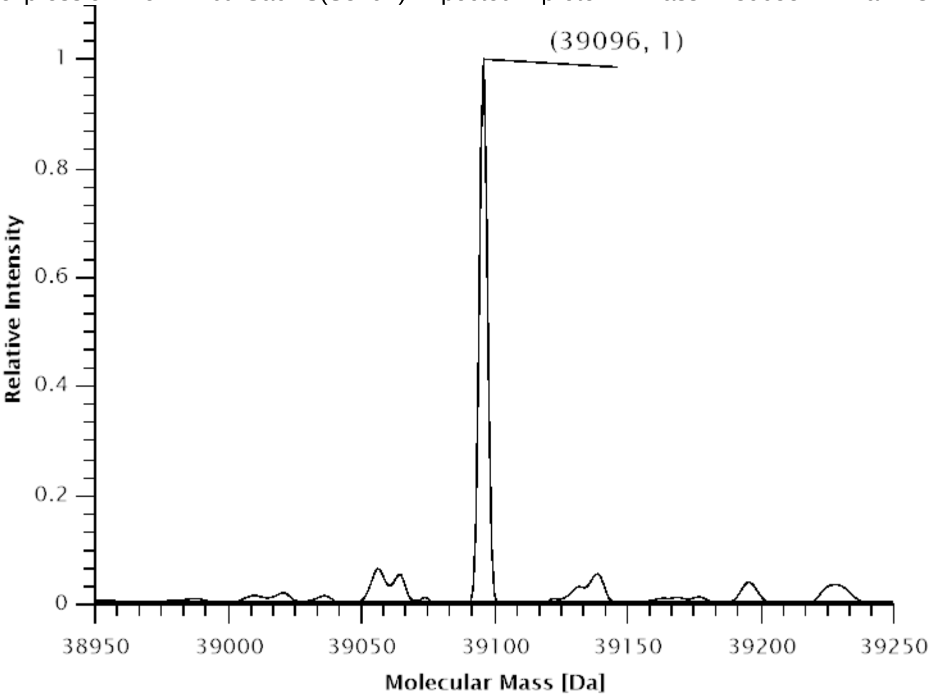

**Figure S91.** Deconvoluted ESI-MS spectrum of SUMO-sfGFP(5x(**38**))-His$_6$ production in *E. coli* B95.ΔA with co-expression of *Mbur*SacRS(S379T).Expected protein mass: 39097.5 Da. Observed mass: 39096 Da.

#### 2.5 ESI-MS data

**Table S1.** Setup of protein production platform with calculated and observed molecular weights of reporter proteinscorresponding to a) SUMO-sfGFP(1x amber), b) SUMO-sfGFP(3x amber), c) SUMO-sfGFP(5x amber), d) SUMO-sfGFP(wild-type). The masses were determined by ESI-MS of the intact proteins (**Figure S60-S88**).

| ncAA | [mM] | E. coli strain[a] | PylRS construct | re-porter | calcu-lated mass [Da] | found mass [Da]] | Δ mass [Da] |
|---|---|---|---|---|---|---|---|
| **2** | 1 | BL21 | Mm | a | 39096.0 | 39094 | 2 |
| **2** | 1 | BL21 | Mbur | a | 39096.0 | 39094 | 2 |
| **3** | 1 | BL21 | Mm | a | 39109.0 | 39105 | 4 |
| **3** | 1 | BL21 | Mbur | a | 39109.0 | 39106 | 3 |
| **9** | 3 | BL21 | Mm(N346A:C348A) | a | 39069.0 | 39065 | 4 |
| **9** | 3 | BL21 | Mbur(N308A:C310A) | a | 39069.0 | 39065 | 4 |
| **11** | 3 | BL21 | Mm(N346A:C348A) | a | 39071.0 | 39068 | 3 |
| **11** | 3 | BL21 | Mbur(N308A:C310A) | a | 39071.0 | 39068 | 3 |
| **13** | 1 | BL21 | Mm(N346A:C348A) | a | 39098.9 | 39095 | 3.9 |

| | | | | | | | |
|---|---|---|---|---|---|---|---|
| **13** | 1 | BL21 | Mbur(N308A:C310A) | a | 39098.9 | 39095 | 3.9 |
| **8** | 3 | BL21 | Mm(N346A:C348A) | a | 39087.0 | 39084 | 3 |
| **8** | 3 | BL21 | Mbur(N308A:C310A) | a | 39087.0 | 39084 | 3 |
| **38** | 0.3 | B-95.ΔA | SmbP-Mb(C313W:W382T) | d | 39023.9 | 39023 | 0.9 |
| **38** | 0.3 | B-95.ΔA | SmbP-Mb(C313W:W382T) | a | 39010.9 | 39010 | 0.9 |
| **38** | 0.3 | B-95.ΔA | SmbP-Mb(C313W:W382T) | b | 39055.2 | 39053 | 2.2 |
| **38** | 0.3 | B-95.ΔA | SmbP-Mb(C313W:W382T) | c | 39097.5 | 39095 | 2.5 |
| **38** | 0.3 | B-95.ΔA | Mbur(C310W:W379T) | d | 39023.9 | 39023 | 0.9 |
| **38** | 0.3 | B-95.ΔA | Mbur(C310W:W379T) | a | 39010.9 | 39011 | 0.1 |
| **38** | 0.3 | B-95.ΔA | Mbur(C310W:W379T) | b | 39055.2 | 39054 | 1.2 |
| **38** | 0.3 | B-95.ΔA | Mbur(C310W:W379T) | c | 39097.5 | 39096 | 1.5 |
| **35** | 1 | B-95.ΔA | SmbP-Mb(C313W:W382T) | a | 39008.9 | 39009 | 0.1 |
| **35** | 1 | B-95.ΔA | SmbP-Mb(C313W:W382T) | b | 39049.0 | 39047 | 2 |
| **35** | 1 | B-95.ΔA | SmbP-Mb(C313W:W382T) | c | 39087.2 | 39084 | 3.2 |
| **35** | 1 | B-95.ΔA | Mbur(C310W:W379T) | a | 39008.9 | 39008 | 0.9 |
| **35** | 1 | B-95.ΔA | Mbur(C310W:W379T) | b | 39049.0 | 39048 | 1 |
| **35** | 1 | B-95.ΔA | Mbur(C310W:W379T) | c | 39087.2 | 39086 | 1.2 |

[a] All DE3

#### 2.6 **Correlation of small-scale fluorescence measurements and shake flask sfGFP protein yields**

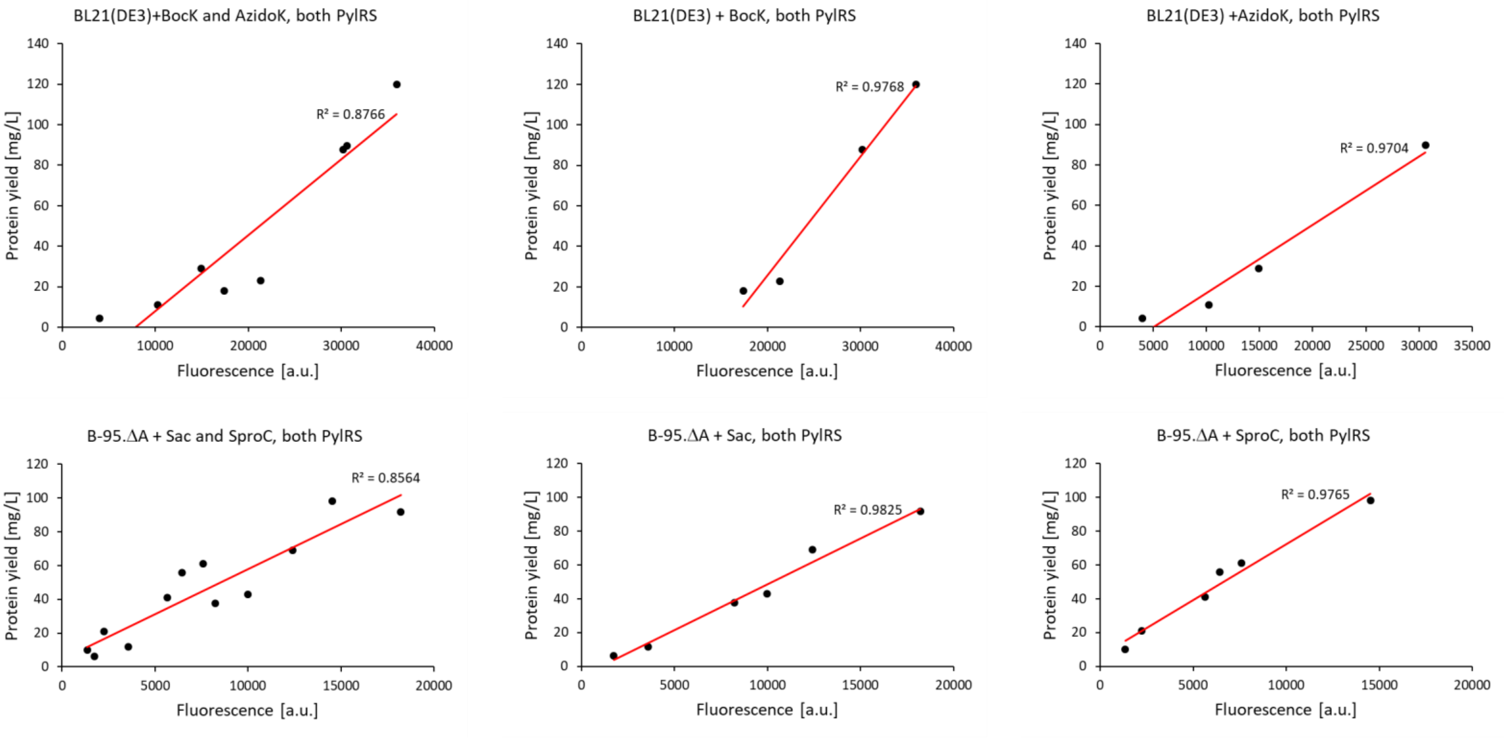

**Figure S92.** Isolated sfGFP protein yields vs. fluorescence.

#### 2.7 **Flow Cytometry Dot plots**

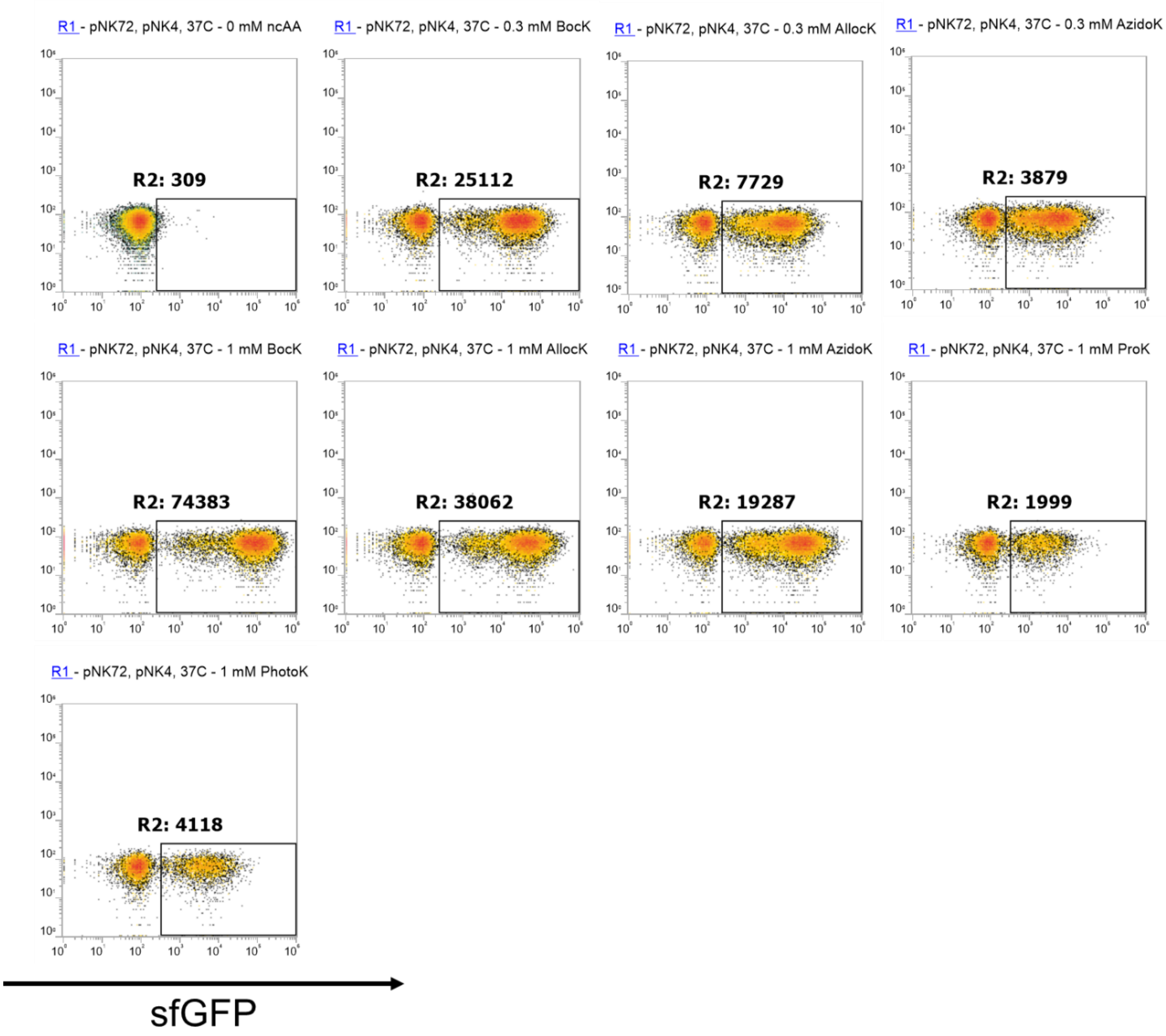

**Figure S93.** Dot plots of the temperature dependent expression of sfGFP(5x amber) at 37°C after 24h with Mbur. pNK72 = Mbur, pNK4 = sfGFP(5x amber).

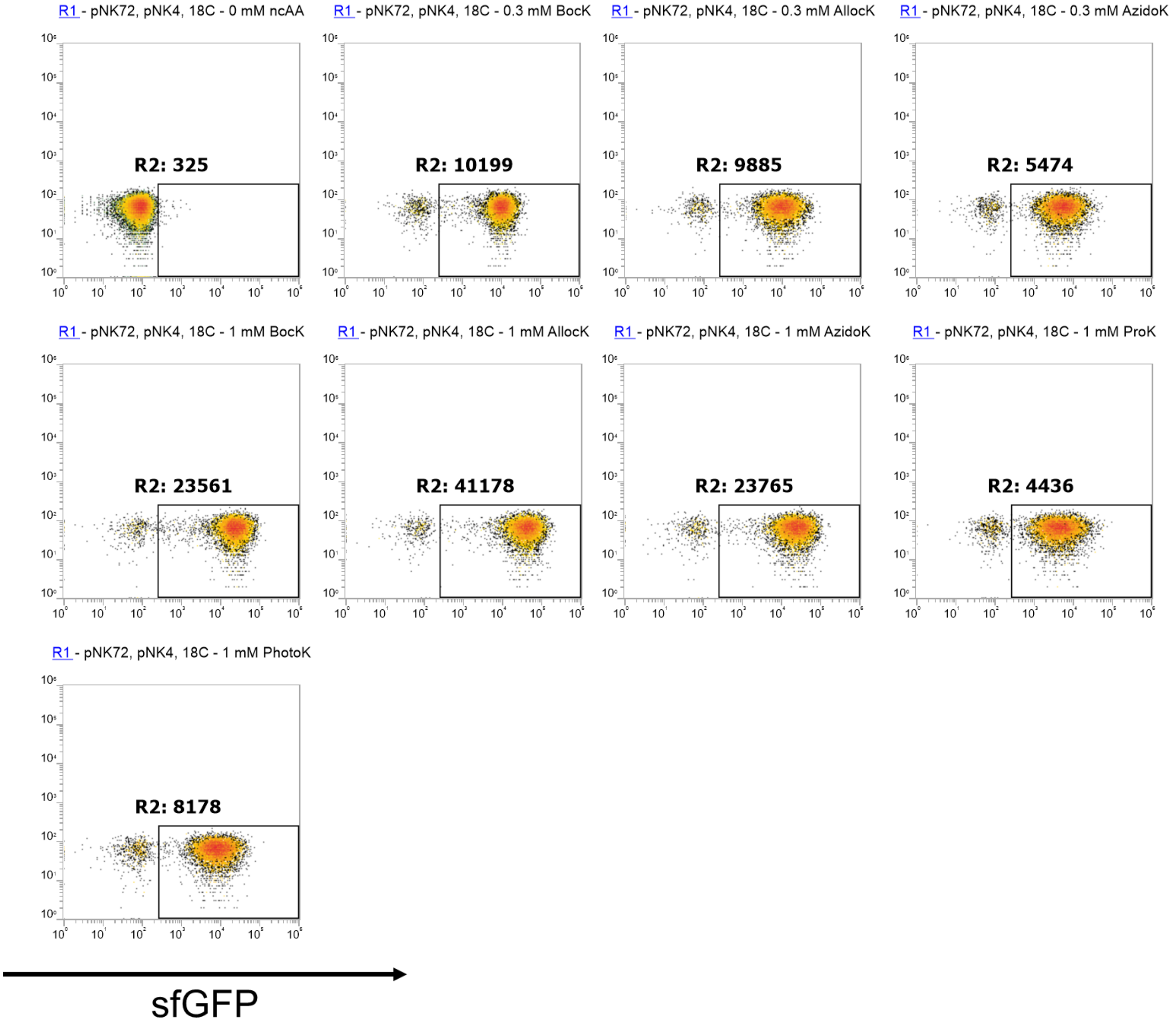

**Figure S94.** Dot plots of the temperature dependent expression of sfGFP(5x amber) at 18°C after 48h with Mbur. pNK72 = Mbur, pNK4 = sfGFP(5x amber).

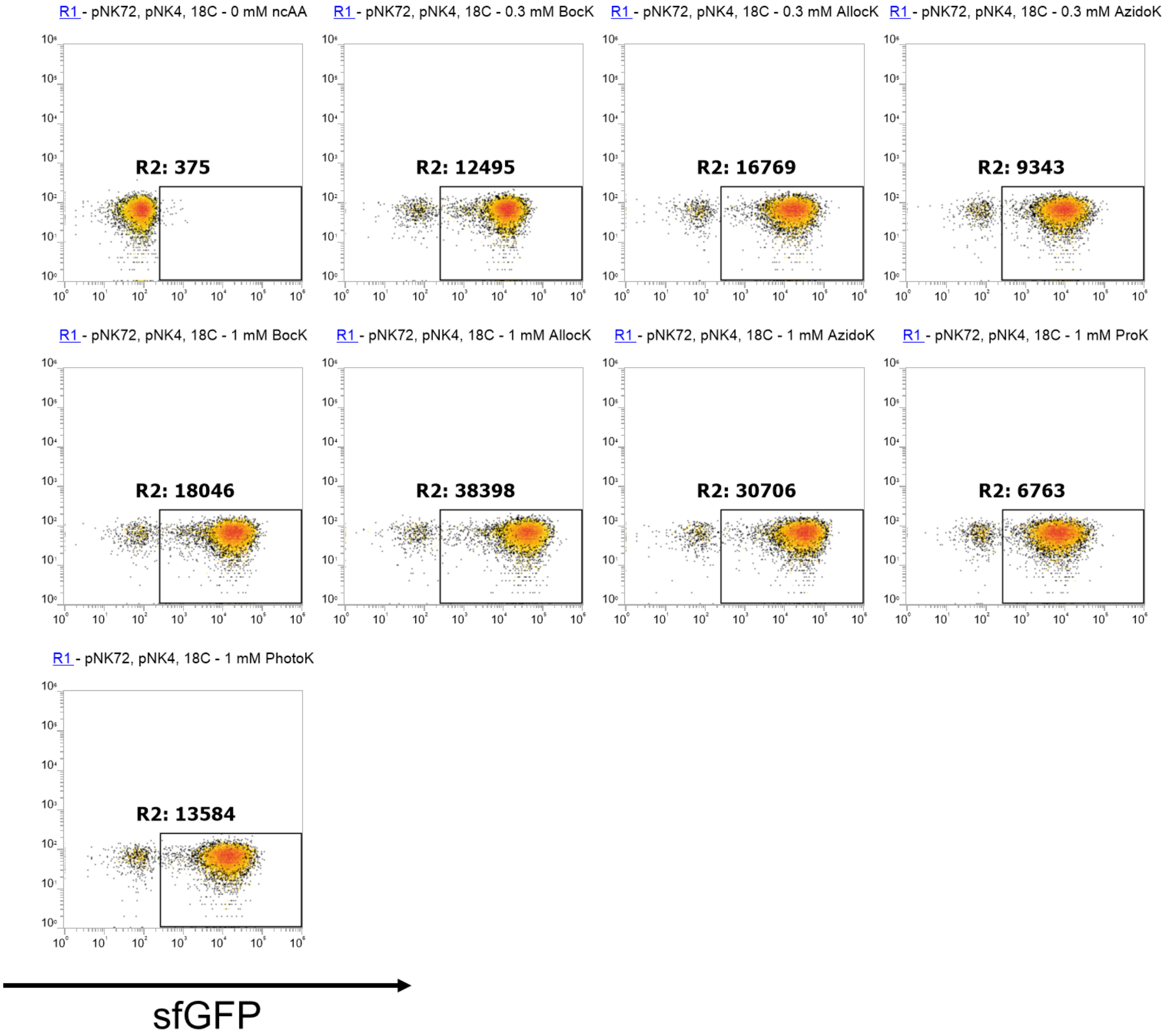

**Figure S95.** Dot plots of the temperature dependent expression of sfGFP(5x amber) at 18°C after 72h with Mbur. pNK72 = Mbur, pNK4 = sfGFP(5x amber).

#### 2.8 **Protein yields**

**Table S2.** Setup of protein production platform with calculated and observed molecular weights of reporter proteinscorresponding to a) SUMO-sfGFP(1x amber), b) SUMO-sfGFP(3x amber), c) SUMO-sfGFP(5x amber), d) SUMO-sfGFP(wild-type). The masses were determined by ESI-MS of the intact proteins (**Figure S60-S88**).

| ncAA | [mM] | E. coli strain[a] | PylRS construct | reporter | protein yield [mg/L][b] | protein yield [mg/L][b] | protein yield [mg/L][b] |
|---|---|---|---|---|---|---|---|
| | | | | | Sample 1 | Sample 2 | Sample 3 |
| **2** | 0.05 | BL21 | Mm | a[c] | 18.5 | 16.2 | 19.1 |
| **2** | 0.05 | BL21 | Mbur | a[c] | 20 | 23.7 | 24.9 |
| **2** | 1 | BL21 | Mm | a | 97.2 | 85.4 | 80.3 |
| **2** | 1 | BL21 | Mbur | a | 123.6 | 108.8 | 127.1 |
| **3** | 0.05 | BL21 | Mm | a[c] | 4.3 | 3.7 | 4.9 |
| **3** | 0.05 | BL21 | Mbur | a[c] | 11.2 | 12.1 | 9.8 |
| **3** | 1 | BL21 | Mm | a | 33 | 28.5 | 25.2 |
| **3** | 1 | BL21 | Mbur | a | 84.9 | 95.8 | 88.3 |
| **9** | 3 | BL21 | Mm(N346A:C348A) | a | 11.9 | 8.7 | 7.6 |
| **9** | 3 | BL21 | Mbur(N308A:C310A) | a | 19.9 | 23.4 | 25.8 |
| **11** | 3 | BL21 | Mm(N346A:C348A) | a | 26.4 | 29.3 | 21.2 |
| **11** | 3 | BL21 | Mbur(N308A:C310A) | a | 52 | 59.4 | 58.1 |
| **13** | 1 | BL21 | Mm(N346A:C348A) | a | 4.6 | 3.7 | 3.9 |
| **13** | 1 | BL21 | Mbur(N308A:C310A) | a | 15.1 | 18.9 | 17.7 |
| **8** | 3 | BL21 | Mm(N346A:C348A) | a | 30.2 | 33.6 | 25.4 |
| **8** | 3 | BL21 | Mbur(N308A:C310A) | a | 51.1 | 58.4 | 60.3 |
| **38** | 0.3 | B-95.ΔA | SmbP-Mb(N313W:W382T) | d | 89.8 | 97.6 | 101.3 |
| **38** | 0.3 | B-95.ΔA | SmbP-Mb(N313W:W382T) | a | 42.2 | 40.1 | 46.9 |
| **38** | 0.3 | B-95.ΔA | SmbP-Mb(N313W:W382T) | b | 13.8 | 11.4 | 9.9 |
| **38** | 0.3 | B-95.ΔA | SmbP-Mb(N313W:W382T) | c | 6.7 | 7.5 | 4.2 |
| **38** | 0.3 | B-95.ΔA | Mbur(C310W:W379T) | d | 113.2 | 118.3 | 108.6 |
| **38** | 0.3 | B-95.ΔA | Mbur(C310W:W379T) | a | 86.2 | 98.9 | 90.1 |
| **38** | 0.3 | B-95.ΔA | Mbur(C310W:W379T) | b | 69.1 | 74.2 | 63.4 |
| **38** | 0.3 | B-95.ΔA | Mbur(C310W:W379T) | c | 38.6 | 33.7 | 40.9 |
| **35** | 1 | B-95.ΔA | SmbP-Mb(N313W:W382T) | a | 60.1 | 54.2 | 52.6 |
| **35** | 1 | B-95.ΔA | SmbP-Mb(N313W:W382T) | b | 19.6 | 24.7 | 18.8 |
| **35** | 1 | B-95.ΔA | SmbP-Mb(N313W:W382T) | c | 10.9 | 8.1 | 11.4 |
| **35** | 1 | B-95.ΔA | Mbur(C310W:W379T) | a | 104.9 | 97.6 | 92.3 |

| | | | | | | | |
|---|---|---|---|---|---|---|---|
| **35** | 1 | B-95.ΔA | Mbur(C310W:W379T) | b | 59.3 | 65.8 | 57.9 |
| **35** | 1 | B-95.ΔA | Mbur(C310W:W379T) | c | 41 | 44.3 | 37.8 |

[a] All DE3. [b] Yield per liter of cell culture; sfGFP absorption was measured directly in the elution fraction before dialysis.

#### 2.9 **Organisms from where the +N PylRS were derived for experiments and *in silico* analysis**

**Table S3: Genomes Used for extraction of PylRS and $tRNA^{Pyl}$ sequences**

| Organism | PylRS abbreviation | Strain | Accession number |
|---|---|---|---|
| *Methanimicrococcus blatticola* | | DSM 13328 | GCF_004363215.1 |
| *Methanococcoides alaskense* | Mala | DSM 17273 | SAMN18250245 |
| *Methanococcoides burtonii* | Mbur | DSM 6242 | GCF_000013725.1 |
| *Methanococcoides methylutens* | | DSM 2657 | GCF_000765475.1 |
| *Methanococcoides vulcani* | | SLH 33 | GCF_900111645.1 |
| *Methanohalobium evestigatum* | | Z-7303 | GCF_000196655.1 |
| *Methanohalophilus euhalobius* | | WG1_MB | SAMN08777283 |
| *Methanohalophilus halophilus* | Mhal | Z-7982 | GCF_001889405.1 |
| *Methanohalophilus levihalophilus* | | DSM 28452 | GCF_017874375.1 |
| *Methanohalophilus mahii* | | DSM 5219 | GCF_000025865.1 |
| *Methanohalophilus portucalensis* | | FDF-1T | GCF_002761295.1 |
| *Methanohalophilus profundi* | | SLHTYRO | GCF_004137855.1 |
| *Methanolobus bombayensis* | | DSM 7082 | GCF_017873415.1 |
| *Methanolobus halotolerans* | | SY-01 | GCF_004745425.1 |
| *Methanolobus profundi* | | Mob M | GCF_900114835.1 |
| *Methanolobus psychrophilus* | Mp | R15 | GCF_000306725.1 |
| *Methanolobus psychrotolerans* | | YSF-03 | GCF_002243045.1 |
| *Methanolobus tindarius* | | DSM 2278 | GCF_000504205.1 |
| *Methanolobus vulcani* | | PL 12/M | GCF_900100715.1 |
| *Methanolobus zinderi* | | DSM 21339 | GCF_013388255.1 |
| *Methanomethylovorans hollandica* | | DSM 15978 | GCF_000328665.1 |
| *Methanosalsum zhilinae* | | DSM 4017 | GCF_000217995.1 |
| *Methanosarcina acetivorans* | | C2A | GCF_000007345.1 |
| *Methanosarcina barkeri* | Mb | 227 | GCF_000970065.1 |
| *Methanosarcina flavescens* | | E03.2 | GCF_001304615.2 |
| *Methanosarcina horonobensis* | | HB-1 | GCF_000970285.1 |
| *Methanosarcina lacustris* | Ml | Z-7289 | GCF_000970265.1 |
| *Methanosarcina mazei* | Mm | S-6 | GCF_000007065.1 |
| *Methanosarcina siciliae* | | T4/M | GCF_000970085.1 |
| *Methanosarcina soligelidi* | | SMA-21 | GCF_000744315.1 |

| | | | |
|---|---|---|---|
| *Methanosarcina spelaei* | | MC-15 | GCF_002287235.1 |
| *Methanosarcina thermophila* | Mt(TM-1) | TM-1 | GCF_000969885.1 |
| *Methanosarcina vacuolata* | | Z-761 | GCF_000969905.1 |
| *Methermicoccus shengliensis* | | DSM 18856 | GCA_013330515.1 |
| *Candidatus Methanomethylophilus alvus* | | Mx1201 | GCF_000300255.2 |
| *Candidatus Methanomassiliicoccus intestinalis* | | MGYG-HGUT-02160 | GCF_902383905.1 |
| *Methanomassiliicoccus luminyensis* | | MGYG-HGUT-02161 | GCF_902383895.1 |

For the source and considerations to the *Mt*PylRS sequence, see supplement **chapter 1.3**.

The ΔN variants were taken from a recent publication[15] and the sequences can be found in the supplementary sequence excel sheet.

### 3. Materials

#### 3.1 Non-Canonical Amino Acids

Unless indicated otherwise, all standard chemicals were purchase from Carl Roth GmbH (Karlsruhe, Germany), Merck (Waltham, MA, USA), VWR International GmbH (Waltham, MA, USA) or Sigma-Aldrich (Taufkirchen, Germany).

**Table S4. Amino acids used in this work.**

| No. | Name | Abbreviation | Cas No. | Company |
|---|---|---|---|---|
| **1** | *N*ε-Allyloxycarbonyl-L-lysine | AllocK | 6298-03-9 | Fluorochem |
| **2** | *N*ε-tert-Butoxycarbonyl-L-lysine | BocK | 2418-95-3 | Budisa Group |
| **3** | *N*ε-Propargyloxycarbonyl-L-lysine | ProK | 1428330-91-9 | Iris Biotech |
| **4** | *N*ε-((2-Azidoethoxy)carbonyl)-L-lysine | AzidoK | 1994331-17-7 | Iris Biotech |
| **5** | 3'-azibutyl-Nε-carbamoyl-L-lysine | PhotoK | 1253643-88-7 | Iris Biotech |
| **6** | *N*ε-benzyloxycarbonyl-L-lysine | BenzK | 1155-64-2 | TCI Deutschland |
| **7** | *O*-methyl-L-tyrosine | O-methyl-Y | 6230-11-1 | Fluorochem |
| **8** | *O*-tert-butyl-L-tyrosine | O-tert-Butyl-Y | 18822-59-8 | Fluorochem |
| **9** | *O*-propargyl-L-tyrosine | O-prop-Y | 610794-20-2 | Iris Biotech |
| **10** | 4-azido-L-phenylalanine | azido-F | 33173-53-4 | Fluorochem |
| **11** | *O*-allyl-L-tyrosine | O-allyl-Y | 107903-42-4 | Iris Biotech |
| **12** | 4-cyano-L-phenylalanine | cyano-F | 167479-78-9 | Alfa Aesar |
| **13** | *O*-$CF_3$-L-tyrosine | O-$CF_3$-Y | 921609-34-9 | Fluorochem |
| **14** | 4-ethynyl-L-phenylalanine | ethynyl-F | 278605-15-5 | Sigma-Aldrich (Merck) |
| **15** | (S)-2-amino-3-(3-(hydroxymethyl)-4-nitrophenyl)propanoic acid | p-oNB-alanin | | Budisa Group |
| **16** | 4-benzoyl-L-phenylalanin | Bpa | 104504-45-2 | Bachem AG |
| **17** | *o*-(2-nitrobenzyl)-3,4-dihydroxyphenylalanine | *m*-oNB-Dopa | | Budisa Group |
| **18** | Sulfotyrosine | sTyr | 956-46-7 | Bachem |
| **19** | (S)-2-Amino-3-(4-((fluorosulfonyl)oxy)phenyl)propanoic acid hydrochloride | FSY | 2227199-79-1 | A2B Chem LLC |
| **20** | O-Phospho-L-tyrosine | pTyr | 21820-51-9 | TCI Chemicals |
| **21** | (S)-2-Amino-3-(4-boronophenyl)propanoic acid | 4-B(OH)-Phe | 76410-58-7 | AmBeeed |
| **22** | 2-Amino-3-(4-(carboxymethyl)phenyl)propanoic acid hydrochloride | CPF | 1803572-24-8 | AmBeeed |
| **23** | (*S*)-2-aminobutyric acid | C4 | 1492-24-6 | TCI Deutschland |
| **24** | (*S*)-2-aminopentanoic acid | C5 | 6600-40-4 | TCI Deutschland |
| **25** | (*S*)-2-aminohexanoic acid | C6 | 327-57-1 | TCI Deutschland |
| **26** | (*S*)-2-aminoheptanoic acid | C7 | 44902-02-5 | Fluorochem |
| **27** | (*S*)-2-aminooctanoic acid | C8 | 116783-26-7 | Fluorochem |
| **28** | (*S*)-2-aminopent-4-enoic acid | C5 alken | 16338-48-0 | Fluorochem |
| **29** | (*S*)-2-aminohex-5-enoic acid | C6 alken | 90989-12-1 | Fluorochem |

| | | | | |
|---|---|---|---|---|
| **30** | (*S*)-2-amino-3-azidopropanoic acid hydrochloride | azido-ala | 105661-40-3 | Iris Biotech |
| **31** | (*S*)-2-amino-4-azidobutanoic acid hydrochloride | AHA | 942518-29-8 | Carl Roth |
| **32** | (*S*)-2-amino-5-azidopentanoic acid hydrochloride | azido-ornithin | 1782935-10-7 | Iris Biotech |
| **33** | (*S*)-2-aminopent-4-ynoic acid | propG | 23235-01-0 | Fluorochem |
| **34** | (*S*)-2-aminohex-5-ynoic acid | Hpg | 98891-36-2 | Toronto Research Chemicals |
| **35** | (*S*)-2-aminohept-6-ynoic acid | Bis-Hpg | 835627-45-7 | Chiralix |
| **36** | (*S*)-2-amino-4-methylpent-4-enoic acid | 4,5-DHL | 87392-13-0 | Fluorochem |
| **37** | (*S*)-2-amino-3-cyanopropanoic acid | CA | 6232-19-5 | Iris Biotech |
| **38** | (*S*)-2-amino-4-cyanobutanoic acid | CHA | 6232-22-0 | Iris Biotech |
| **39** | (*S*)-2-amino-3-cyclopropylpropanoic acid | cyclo-ala | 102735-53-5 | Fluorochem |
| **40** | *S*-propargyl-L-cystein | SproC | 3262-64-4 | Fluorochem |
| **41** | L-ethionine | Eth | 13073-35-3 | Sigma-Aldrich (Merck) |
| **42** | L-methionine sulfoxide | Met-sulfoxide | 3226-65-1 | Sigma-Aldrich (Merck) |
| **43** | *S*-allyl-L-cystein | Sac | 21593-77-1 | TCI Deutschland |

#### 3.2 **Oligonucleotides**

All oligonucleotides were purchased from Sigma-Aldrich (Taufkirchen, Germany), resuspended in dd$H_2O$ to a final concentration of 100 µM and stored at -20 °C. Working concentration of primers were adjusted to 10 µM. Primers shorter than 50 bp were generally purchased in desalted form. Primers between 50-80 bp were ordered in in cartridge-purified form, while longer ones were obtained in HPLC-purified grade. Primers are not listed because far over hundred were used and the utility of this information is limited at best.
